# *The Tabulae Paralytica:* Multimodal single-cell and spatial atlases of spinal cord injury

**DOI:** 10.1101/2023.06.23.544348

**Authors:** Michael A. Skinnider, Matthieu Gautier, Alan Yue Yang Teo, Claudia Kathe, Thomas H. Hutson, Achilleas Laskaratos, Alexandra de Coucy, Nicola Regazzi, Viviana Aureli, Nicholas D. James, Bernard Schneider, Michael V. Sofroniew, Quentin Barraud, Jocelyne Bloch, Mark A. Anderson, Jordan W. Squair, Grégoire Courtine

## Abstract

Here, we introduce the *Tabulae Paralytica*—a compilation of four atlases of spinal cord injury (SCI) comprising a single-nucleus transcriptome atlas of half a million cells; a multiome atlas pairing transcriptomic and epigenomic measurements within the same nuclei; and two spatial transcriptomic atlases of the injured spinal cord spanning four spatial and temporal dimensions. We integrated these atlases into a common framework to dissect the molecular logic that governs the responses to injury within the spinal cord. The *Tabulae Paralytica* exposed new biological principles that dictate the consequences of SCI, including conserved and divergent neuronal responses to injury; the priming of specific neuronal subpopulations to become circuit-reorganizing neurons after injury; an inherent trade-off between neuronal stress responses and the activation of circuit reorganization programs; the necessity of reestablishing a tripartite neuroprotective barrier between immune-privileged and extra-neural environments after SCI; and a catastrophic failure to form this barrier in old mice. We leveraged the *Tabulae Paralytica* to develop a rejuvenative gene therapy that reestablished this tripartite barrier, and restored the natural recovery of walking after paralysis in old mice. The *Tabulae Paralytica* provides an unprecedented window into the pathobiology of SCI, while establishing a framework for integrating multimodal, genome-scale measurements in four dimensions to study biology and medicine.

---

Spinal cord injury (SCI) irreversibly damages neural tissues, leading to permanent and devastating loss of neurological functions^1, 2^. Advances in medical management^3^ and neurotechnologies^4–9^ have improved survival and allow clinicians to address many aspects of neurological dysfunction after SCI. However, decades of investigations culminating in large-scale clinical trials have yet to identify safe and effective therapies to repair the injured spinal cord^1, 10^.

A SCI triggers a cascade of molecular and cellular responses involving inflammatory cell infiltration and cytokine release, apoptosis, demyelination, excitotoxicity, ischemia, and the formation of a fibrotic scar surrounded by an astrocyte border^2, 10–16^. Altering the course of this cascade to repair the injured spinal cord will require a complete understanding of how neural and non-neural cells coordinate the response to SCI over time and throughout the lesion microenvironment^2, 17, 18^. Previous attempts to delineate the molecular logic governing this response initially turned to bulk transcriptomics and proteomics of the entire lesion^15, 19–26^. However, these attempts were technically limited in their ability to resolve cell-type-specific molecular programs triggered by injury, or else focused on isolated aspects of the injury response.

Emerging genome-scale technologies are poised to overcome these limitations. First, single-cell transcriptomics could unveil the molecular programs triggered by SCI within hundreds of thousands of cells composing the injured spinal cord^27–30^. Second, multi-omic methods that couple single-cell transcriptomics to chromatin accessibility profiling within the same cell could access the regulatory programs that govern the response to injury^31–34^. Third, spatial transcriptomics could circumscribe these transcriptional and regulatory programs within the complex cytoarchitecture of the lesion microenvironment^35–42^.

Here, we leveraged single-cell transcriptomics, multiomics, and spatial transcriptomics in mice to establish the *Tabulae Paralytica*, or ‘atlases of spinal cord injury’ (**Fig. 1a**). Together, these atlases, comprising 482,825 individual cells spanning 18 experimental conditions and 71,499 spatial barcodes mapped onto the three-dimensional architecture of the injured spinal cord, provide an unprecedented window into the pathobiology of SCI. We provide an interactive web application to explore these atlases at http://tabulaeparalytica.com.

**Fig. 1.**
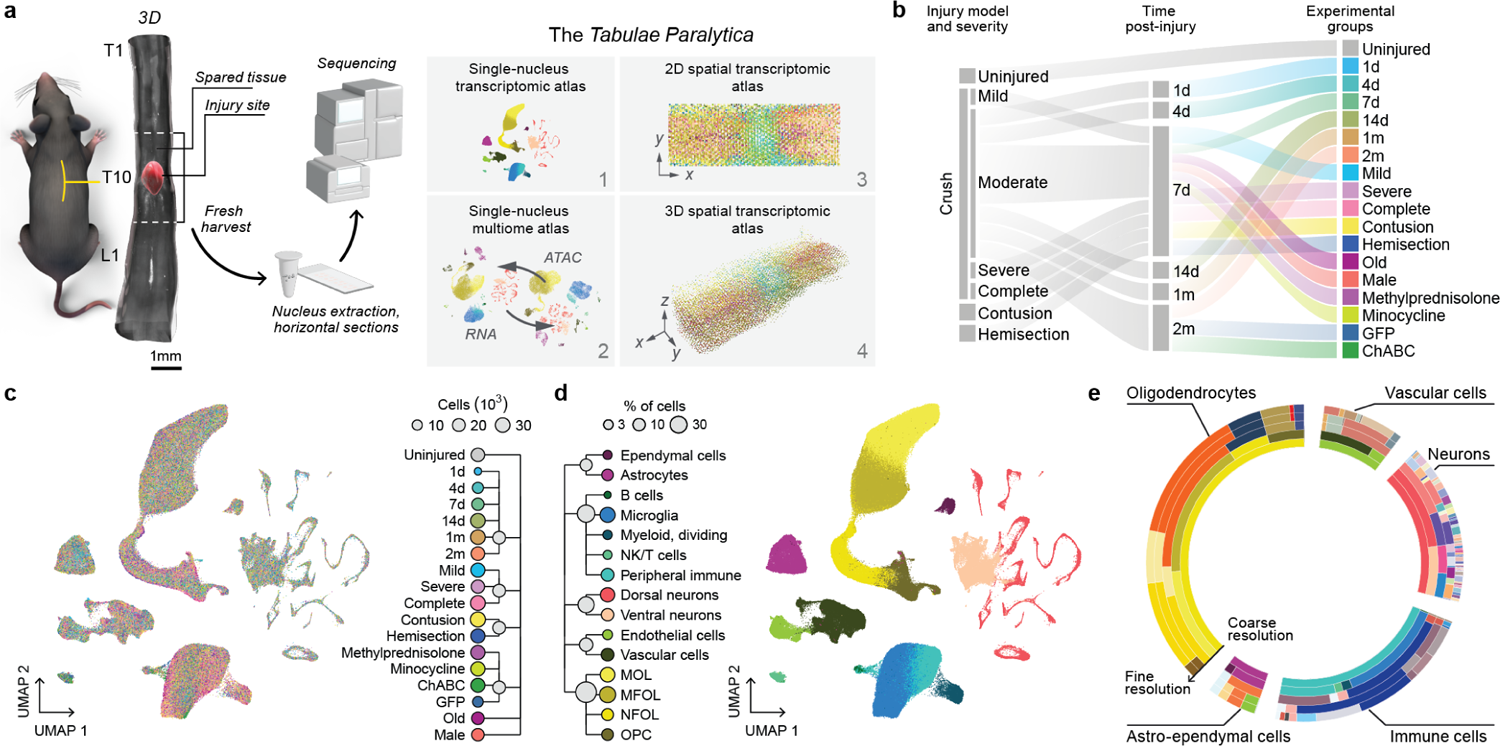
Overview of the *Tabulae Paralytica* and the snRNA-seq atlas. **a**, Schematic overview of the *Tabulae Paralytica*. **b,** Schematic overview of the snRNA-seq atlas. **c,** UMAP visualizations of the 435,099 cells in the snRNA-seq atlas, colored by experimental condition. Dendrogram shows major groups of experimental conditions. **d,** As in **c,** but colored by cell type. Dendrogram shows the first two levels of the clustering tree of spinal cord cell types. **e,** Sunburst plot showing cell type proportions in the snRNA-seq atlas at each level of the clustering tree, from broadest (innermost ring) to most granular (outermost ring).

## Results

### Single-nucleus RNA-seq of the injured spinal cord

Single-cell transcriptomics of injured spinal cord tissues presents unique challenges: many cell types do not survive harsh dissociation protocols, and surviving cells express a dissociation-induced stress signature^43–46^. To overcome these limitations, we optimized single-nucleus RNA-sequencing (snRNA-seq) protocols for the injured spinal cord. We acquired high-quality transcriptomes from all the major cell types from injured and uninjured spinal cords (**Supplementary Fig. 1a-i**). Comparison with 16 published datasets^47–62^ confirmed that our protocol recovered the complete repertoire of spinal cord cell types, even when applied to injured spinal cord tissues (**Supplementary Fig. 1j-q**).

### A comprehensive set of experiments for large-scale snRNA-seq

We next leveraged these optimized protocols to conduct snRNA-seq profiling of the injured spinal cord across a comprehensive set of experimental conditions and injury models that aimed to capture the multifaceted responses to SCI and how pharmacological interventions may alter these responses (**Fig. 1b-c** and **Supplementary Figs. 2** and **3**).

The pathobiological responses activated in the injured spinal cord depend on the severity and mechanism of the initial insult, and evolve over the following days, weeks, and months^1, 3, 18^. To capture these responses, we profiled the spinal cord of uninjured mice at 1, 4, 7, 14, 30, and 60 days after mid-thoracic SCI. Next, we devised a progression of injury severities that led to mild, moderate, severe, and complete functional impairments. Finally, we profiled the spinal cord following SCI induced by different mechanisms of injury, including crush^15, 63^, contusion^64, 65^, and dorsal hemisection^12^.

In humans, immune responses differ across the lifespan and between males and females, with broad implications for disease initiation and progression^66–69^. To evaluate the impact of sex and age on the cell-type-specific molecular programs activated by SCI, we profiled injured spinal cords from male and female mice, and from young and old mice.

Finally, we asked whether single-cell techniques could provide insights into the molecular mechanisms of pharmacotherapies for SCI. To address this question, we profiled the spinal cord of mice treated with three of the most extensively investigated clinical and experimental interventions: methylprednisolone^70–72^, minocycline^73–76^, and chondroitinase ABC (ChABC)^77–84^.

### A single-nucleus transcriptomic atlas of SCI

We exploited this progression of 18 experimental conditions to establish a single-cell atlas of SCI, profiling the spinal cords of mice from each condition by snRNA-seq. We obtained high-quality transcriptomes for a total of 435,099 nuclei from 54 mice (**Fig. 1c-d** and **Supplementary Fig. 4**).

To identify both coarse cell types and more granular sub-types, we subjected the entire dataset to multiple rounds of clustering at increasingly fine-grained resolutions. This procedure identified all the major cell types of the spinal cord and allowed us to establish a comprehensive catalog of 175 more granular subpopulations. We organized these subpopulations into a clustering tree^85^ that recapitulated the known cellular hierarchy of the spinal cord (**Fig. 1e** and Supplementary Fig. 5)^2^,9,47–62,86–88.

Coarse clustering identified cells originating from immune, astroependymal, vascular, oligodendrocyte, and neuronal lineages (**Supplementary Fig. 6**). Within each of these lineages, we first explored the evolution of each subpopulation over time and with increasing injury severity.

The immune compartment comprised 106,619 immune cells spanning both central nervous system-resident and infiltrating immune cells, which are known to coordinate sequential phases of the acute response to SCI (**Fig. 2a** and **Supplementary Fig. 7a-c**)^11^. *P2ry12*-expressing homeostatic microglia were transcriptionally distinguishable from reactive microglia, which expressed *Tgfbr1*. Peripheral lymphocytes comprised B cells, NK cells, and CD8 T cells, whereas myeloid cells subdivided into proliferating and fate-committed subpopulations, as well as neutrophils, macrophages, and dendritic cells. Macrophages encompassed border-associated macrophages (expressing *H2-Aa*) that mediate immune responses in perivascular spaces^89^, as well as distinct inflammatory (*Ifi211*), chemotaxis-inducing (*Cd300a*), and thrombospondin-sensing (*Htr2b*) subpopulations^58^.

**Fig. 2.**
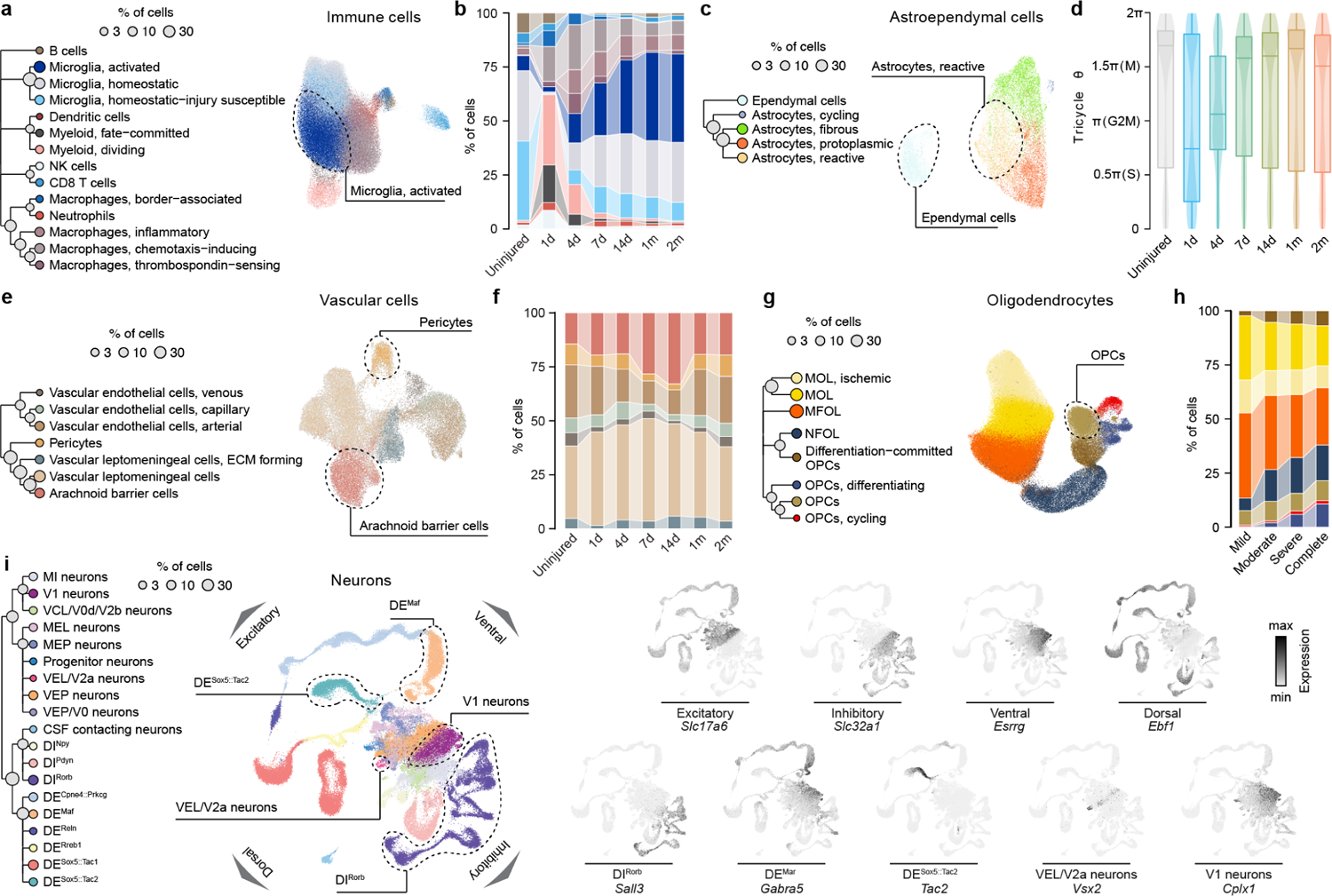
Cell types and subtypes of the uninjured and injured mouse spinal cord. **a**, UMAP visualization of 106,619 immune cells in the snRNA-seq atlas, colored by cell type. Dendrogram shows the bottom four levels of the clustering tree, subset to show immune cell subtypes only. **b,** Proportions of each immune cell subtype across timepoints. **c,** As in **a,** but for 25,211 astroependymal cells. **d,** Cell cycle stages assigned to astroependymal cells at each timepoint post-injury. **e,** As in **a,** but for 40,620 vascular cells. **f,** Proportions of each vascular cell subtype across timepoints. **g,** As in **a,** but for 182,334 oligodendrocytes. **h,** Proportions of each oligodendrocyte subtype across injury severities. **i,** Left, as in **a,** but for 80,315 neurons. Right, UMAP visualization showing expression of key marker genes for select neuronal subtypes.

Our single-cell atlas recapitulated the known evolution of the immune response over the first two months following SCI^11^. Extensive infiltration of peripheral immune cells peaked between 7 and 14 days, paralleling the initiation and slow stabilization of microglial activation (**Fig. 2b**). The relative proportion of homeostatic microglia decreased gradually with injury severity, whereas the proportions of chemotaxis-inducing and inflammatory macrophages expanded (**Supplementary Fig. 7d**). These shifts in the composition of the immune compartment reflected an increasingly profound disruption of the blood-brain barrier.

When a SCI occurs, astrocytes form a barrier that surrounds the fibrotic lesion core to protect viable neural tissue from infiltrating immune cells^15, 90–92^. In our single-cell atlas, the astroependymal compartment comprised 25,211 nuclei, spanning grey matter-resident protoplasmic astrocytes (*Nwd1*, *Gfap*^low^), white matter-resident fibrous astrocytes (*Slc4a4*, *Gfap*^high^), reactive subpopulations that we subdivided based on the expression of *Aldoc*, *Tmem47*, and *Gja1*, and ependymal cells (*Dnah12*) (**Fig. 2c** and **Supplementary Fig. 8a-c**).

The proportion of protoplasmic astrocytes declined gradually over the first few days after injury (**Supplementary Fig. 8d**). The extent of this loss correlated with the severity of the injury (**Supplementary Fig. 8e-f**). This loss contrasted with the reactive astrocyte compartment, which expanded immediately after injury, and persisted into the chronic stage (**Supplementary Fig. 8g**). These responses coincided with the entry of astrocytes into the cell cycle starting at 1 day, with peak proliferation observed at 4 days (**Fig. 2d**). By 7 days, astrocytes had returned to G1/G0, indicating that this proliferation was completed.

The cerebrovasculature comprises an arteriovenous axis of arteries, arterioles, capillaries, venules, and veins^51, 93, 94^. Together, these vessels form the blood-brain barrier that separates the immune-privileged spinal cord parenchyma from the extra-neural environment^95–99^. In our single-cell atlas, the vascular compartment comprised 40,620 cells, which subdivided into endothelial cells, vascular leptomeningeal cells (VLMCs, *Pdgfra*^high^)^51, 93^ and pericytes (**Fig. 2e** and **Supplementary Fig. 9a-c**). Endothelial cells were further sub-divided by their arteriovenous zonation, with distinct arterial (*Emcn*), capillary (*Meox1*), and venous (*Slc38a5*) subpopulations. VLMCs, which are fibroblast-like cells located between astrocyte end-feet and endothelia that express fibrilforming collagens^51, 93^, were further separated into homeostatic and extracellular matrix-forming subpopulations, as well as arachnoid barrier cells (*Slc47a1*) that are involved in cerebrospinal fluid maintenance^51, 100^.

Our single-cell atlas revealed an immediate and severity-dependent disruption of the blood-brain barrier following SCI. This disruption encompassed a contraction of the endothelial and pericyte compartments, and a concomitant expansion in the proportion of VLMCs (**Fig. 2f** and **Supplementary Fig. 9d**). Moreover, vascular cells exhibited a severity-dependent increase in the expression of genes associated with blood-brain barrier dysfunction that increased over the first four days^101^ (**Supplementary Fig. 9e-f**). By 7 days, proliferation of arachnoid barrier cells marked the onset of barrier formation from the cerebrospinal fluid space (**Fig. 2f**). The formation of the cerebrospinal fluid barrier was followed by the reestablishment of the blood-brain barrier, marked by the proliferation of endothelial cells and pericytes, and resolution of blood-brain barrier dysfunction. The resolution of peripheral immune cell invasion coincided with the reestablishment of these barriers.

The oligodendrocyte lineage comprised 182,334 cells that were distributed along a continuous developmental trajectory, spanning oligodendrocyte precursor cells (OPCs), differentiation-committed oligodendrocyte precursors (COPs), newly formed oligodendrocytes (NFOLs), myelin-forming oligodendrocytes (MFOLs), and mature oligodendrocytes (MOLs) (**Fig. 2g** and **Supplementary Fig. 10a-c**)^51,102^. The proportion of oligodendrocytes decreased at 1 day, consistent with the notion that they are sensitive to the ischemic environment that develops after SCI (**Supplementary Fig. 10d**)^103^. By 4 days, we observed a severity-dependent expansion in the proportion of OPCs, which preceded the reinstatement of a near-normal oligodendrocyte compartment by 7 days (**Fig. 2h** and **Supplementary Fig. 10d**).

The spinal cord encompasses dozens of anatomically, functionally, and transcriptionally distinct neuronal _subpopulations_^2^,9,47–51,53,56,57,59,60,86–88_. The scale of our_ single-cell atlas, which comprised 80,315 single-neuron transcriptomes, allowed us to identify 60 distinct subpopulations of neurons (**Fig. 2i** and **Supplementary Fig. 11**). Neuronal subpopulations were parcellated into dorsal versus ventral, excitatory versus inhibitory, and local (*Nfib*) versus long-projecting (*Zfhx3*) populations^59^. We identified well-studied dorsal excitatory (*Tac1*, *Reln*, *Cck*) and inhibitory (*Rory*, *Npy*, *Gal*) subpopulations, as well as dl5/dIL deep dorsal subpopulations expressing *Lmx1b*^47, 51, 86, 88^. A group of ventral neuron subpopulations included developmentally defined V0, V1, V2, V3 (*Sim1*) neurons^86, 104^; Renshaw cells (*Calb1*, *Chrna2*, *Slco5a1*)^105, 106^; cerebrospinal fluid-contacting neurons (CSF-N; *Pdk1l2*)^51, 107, 108^; and motor neurons (*Isl1*)^109^. V0 neurons subclustered into V0c (*Chat*, *Pitx2*)^110^, V0v (*Evx1*)^111, 112^, V0g (*Slc17a6*, *Pitx2*)^110^, and V0d (*Chat*, *Evx1*^OFF^, *Gabra1*)^111, 112^ subpopulations. V1 (*En1*) neurons subclustered into four subpopulations expressing *Pou6f2*, *Foxp2*, *Mafa*, and *Sp8*, respectively^113^. Within V2 neurons, we identified one inhibitory subpopulation of V2b neurons expressing *Gata2* and *Gata3*^114, 115^, as well as multiple subpopulations of developmentally defined V2a neurons (*Vsx2*)^39, 86, 116–118^ that could be separated into local (*Nfib*) and long-projecting (*Zfhx3*) subpopulations^59^. Together, this atlas establishes a single-cell taxonomy of the mouse spinal cord, and delineates the impact of injury severity and time on the repertoire of cell types within the injured spinal cord.

### Conserved and divergent neuronal responses to SCI

A common feature of many insults to the nervous system is that specific neuronal subpopulations exhibit disproportionate susceptibility or resilience to the insult^119–125^. However, whether different neuronal subtypes within the spinal cord respond differentially to injury remains unknown.

To address this possibility, we compared the proportions of neurons from each subpopulation between injured and uninjured spinal cords. Our single-cell atlas confirmed the expected severity-dependent loss of neurons after injury (**Fig. 3a** and **Supplementary Fig. 12a**). However, there were minimal changes in the relative proportions of each neuronal subpopulation, suggesting that spinal cord neurons are, in general, equally vulnerable to SCI (**Supplementary Fig. 12b**).

The sole exception arose from cerebrospinal fluidcontacting neurons^51, 107, 108^, which exhibited a unique resilience to SCI (**Fig. 3b** and **Supplementary Fig. 12c**). This resilience was consistent across every comparison of injured and uninjured spinal cords, and became more pronounced with increasingly severe injuries (**Supplementary Fig. 12d-e**). Immunohistochemistry validated this resilience of cerebrospinal fluid-contacting neurons (**Supplementary Fig. 12f-g**).

**Fig. 3.**
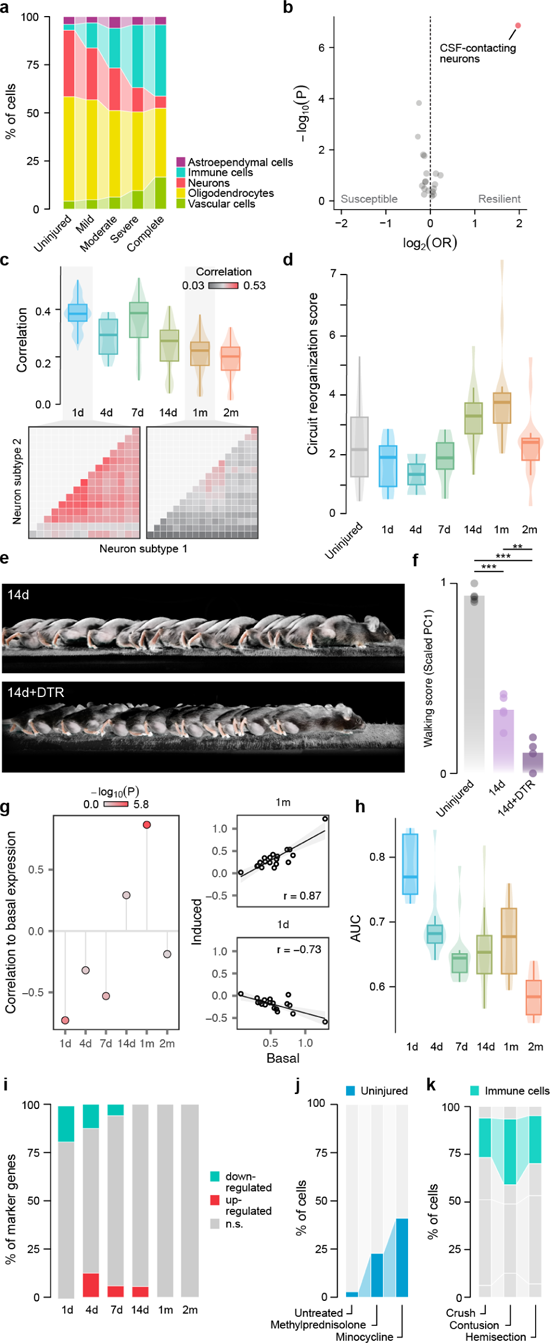
Biological principles governing the response to SCI. **a**, Proportions of each major spinal cord cell type across injury severities. **b,** Susceptible and resilient subtypes of spinal cord neurons. Volcano plot shows log_2_-odds ratios comparing neuron proportions between the uninjured spinal cord and each injured condition at 7 days post-injury (x-axis) versus statistical significance (t-test, y-axis). **c,** Top, transcriptome-wide correlations of DE signatures between each pair of neuron subtypes, across timepoints. Bottom, correlation matrices highlighting conserved DE at 1 day and divergent DE at 1 month. **d,** Expression of the circuit reorganization module in local Vsx2-expressing neurons at each timepoint post-injury. **e,** Chronophotography of walking in Vsx2^Cre^ mice after spontaneous recovery and in mice that received viral injections of AAV5-CAG-FLEX-DTR to induce cell type-specific neuronal ablation prior to SCI. **f,** Walking performance of uninjured mice (*n* = 5), mice after spontaneous recovery (*n* = 5), and in mice with *Vsx2*^ON^ neuron ablation in the lower thoracic spinal cord (*n* = 4). **g,** Left, correlations between basal and injury-induced expression of the circuit reorganization module across neuron subtypes, at each timepoint post-injury. Right, scatterplots showing the correlation between basal and induced expression across neuron subtypes at 1 day and 1 month. **h,** Intensity of the transcriptional perturbation within each neuronal sub-type, as quantified by Augur, across timepoints. **i,** Proportion of neuronal marker genes that are up- and down-regulated at each timepoint. **j,** Proportion of neurons assigned an uninjured transcriptional phenotype in mice treated with methylprednisolone or minocycline, as compared to neurons from the untreated spinal cord. **k,** Proportions of immune cells in the spinal cord across injury models.

We asked whether cerebrospinal fluid-contacting neurons express unique transcriptional programs in response to injury that could explain this resilience (**Supplementary Fig. 12h**). Relative to other neuronal subpopulations, these neurons upregulated genes associated with cell adhesion (*Cntnap5c*), angiogenesis (*Rhoj*), and acute tissue remodeling (*Timp3*).

In contrast, other neuronal subpopulations exhibited a homogenous degree of vulnerability to SCI. We hypothesized that this homogeneity may coincide with the activation of shared transcriptional programs in response to injury. Indeed, we found that SCI initially triggered molecular responses that were broadly conserved across all neuronal subpopulations (**Fig. 3c** and **Supplementary Fig. 13a**). These responses gradually diverged over the following two months, as individual neuronal subpopulations activated increasingly distinct transcriptional programs (**Fig. 3c**).

This homogeneity compelled us to characterize this conserved early response of neurons to SCI (**Supplementary Fig. 13b**). We found that upregulation of immune response pathways (*Fkbp3*, *Stat3*), apoptotic programs (*Pdcd5*, *Bex3*, *Parp3*), and mitochondrial membrane disruption (*Atpif1*, *Ndufa7*, *Fis1*) were the hallmarks of this response. Conversely, neurons downregulated core neuronal functions, including neurotransmitter release (*Lyn*), ion channel expression (*Kcqn1*, *Slc24a5*), and cell adhesion (*Ctnn1*, *Calr*, *Add3*, *Ctnna3*).

We next explored the gradual divergence of neuronal responses at later time points. This divergence coincided with the known timescale at which circuit reorganization mediates the natural recovery of neurological functions after SCI^2^. We therefore reasoned that this divergence might reflect subpopulation-specific circuit reorganization. Indeed, we found that only a few neuronal subpopulations upregulated genes associated with projection growth and morpho-genesis, which occurred between 14 days and 2 months after injury (**Fig. 3d** and **Supplementary Fig. 13c**).

Within these subpopulations, local *Vsx2*^ON^ (*Nfib*) neurons exhibited the greatest upregulation of genes associated with circuit reorganization (**Fig. 3d** and **Supplementary Fig. 13d-e**). To link this transcriptional response to neurological recovery, we ablated *Vsx2*^ON^ neurons in the thoracic spinal cord two weeks prior to a moderate SCI. Ablating *Vsx2*^ON^ neurons prevented the natural recovery of walking in these mice (**Fig. 3e-f** and **Supplementary Fig. 13f-h**).

Local *Vsx2* neurons also exhibited the highest expression of genes associated with circuit reorganization in the uninjured spinal cord (**Supplementary Fig. 13i**). This observation raised the intriguing possibility that specific neuronal subpopulations may be intrinsically primed to serve as circuit-reorganizing cells after injury^2^. To study this possibility, we correlated the expression of circuit reorganization programs in each subpopulation of uninjured neurons with the upregulation of the same programs after injury. We identified a striking correlation between basal and injury-induced circuit reorganization programs between 14 days and 1 month post-injury, when these programs were maximally upregulated (**Fig. 3g** and **Supplementary Fig. 13j**). This time course coincides precisely with the temporal window of opportunity for the circuit reorganization that mediates natural recovery after SCI^2, 126^. Together, these findings imply that specific neuronal subpopulations are endowed with the inherent potential to become circuit-reorganizing cells that support neurological recovery.

Our analyses thus far exposed a temporal continuum between early-conserved and late-diverging neuronal responses following SCI. We sought to quantify the relative intensity of these time-dependent responses. To enable this quantification, we assessed the relative degree of transcriptional perturbation within each neuronal subpopulation over the course of recovery after SCI using Augur^60, 127^. Augur is a machinelearning framework that quantifies the relative magnitude of the transcriptional response within any given cell type to an arbitrary perturbation, a procedure we refer to as cell type prioritization. This prioritization revealed a pronounced neuronal response at 1 day that decreased in intensity over the subsequent days, and thereafter remained constant (**Fig. 3h**).

Based on these observations, we propose a model in which all neurons undergo a profound and broadly conserved transcriptional response immediately after injury that leads to a dichotomous outcome of survival versus apoptosis. Over the subsequent weeks, the surviving neurons exhibit gradually divergent transcriptional responses to injury, whereby only specific subpopulations upregulate molecular programs associated with circuit reorganization. The degree of this injury-induced upregulation is encoded in the basal transcriptional state of each neuronal subpopulation, suggesting that specific subpopulations are primed to serve as circuit-reorganizing neurons following injury^2^.

### Surviving neurons remain differentiated after CNS injury

Single-cell studies have shown that neurons in the injured peripheral nervous system undergo dedifferentiation and loss of transcriptional identity following axonal injury^128, 129^. We asked whether similar biological principles dictate neuronal responses in the injured CNS.

Contrary to single-cell analyses of peripheral neurons, we failed to identify a separate cluster of dedifferentiated neurons within the injured spinal cord at any timepoint (**Fig. 2i**). We reasoned that differential expression (DE) of neuronal marker genes could identify more subtle loss of transcriptional identity. However, we found that the vast majority of subpopulation-specific marker genes were neither up- nor downregulated across the entire time course of SCI (**Fig. 3i** and **Supplementary Fig. 14a**). This observation was robust to the statistical threshold used to identify neuronal marker genes (**Supplementary Fig. 14b-c**).

Collectively, these observations raise the possibility that transient loss of neuron transcriptional identity after injury may be a mechanism by which the peripheral nervous system maintains the distinct capacity to regrow severed nerves^130, 131^. However, the central nervous system fails to recruit this mechanism after injury.

### Growth-facilitating and inhibiting molecule expression

Following SCI, neural and non-neural cells express multiple families of molecules that can facilitate or inhibit axon growth and circuit reorganization^15, 63, 132–134^. These molecular pathways have historically been the main targets for interventions that aim to promote spinal cord repair^10^, but the identities of the cells that produce these molecules are not well characterized.

Our single-cell atlas provides a resource to explore the production of inhibitory and facilitating molecules across the entire repertoire of cell types in the injured spinal cord. For example, we and others previously showed that laminins provide a permissive substrate for axon growth^15, 63, 135^, but the cells that express these molecules have not been identified. We found that laminins (*Lama1*, *Lama2*, *Lama4*, *Lama5*) were expressed primarily by VLMCs, identifying these cells as a target to upregulate the expression of axon growth-supporting molecules for spinal cord repair (**Supplementary Fig. 15a-d**). A second example comes from the family of chondroitin sulfate proteoglycans (CSPGs), which are known to play contrasting roles in inhibiting or supporting axon growth^15, 63, 77–79, 136^. Among inhibitory CSPGs, we found that OPCs are responsible for the expression of *Acan*, *Vcan*, and *Ncan*, whereas astrocytes were the dominant producers of *Bcan* (**Supplementary Fig. 15e-h**). OPCs were also the dominant producer of axon growth-supportive CSPGs, such as *Cspg4* and *Cspg5* (**Supplementary Fig. 15i-j**).

Various studies reported that the enzyme ChABC can digest CSPGs^80, 81, 137–139^, which are a main constituent of perineuronal nets^82, 140^, and that this digestion may open opportunities for circuit reorganization^10, 80, 81, 141^. We employed cell type prioritization to identify the cell types that were most transcriptionally perturbed by ChABC treatment, which pointed to CSPG-producing cell types including OPCs and VLMCs (**Supplementary Fig. 15k**). We then asked how ChABC influences genes associated with circuit reorganization. After ChABC administration, neurons upregulated genes associated with focal adhesion, as previously described^142^, including *Itgav*, *Sorbs3*, *Itgb5*, and *Ilk* (**Supplementary Fig. 15l-m**). Despite these changes, however, high-resolution kinematics showed no significant benefit for the recovery of walking (**Supplementary Fig. 3d**).

Collectively, these results illustrate how our single-cell atlas can be used as a resource to identify the cell-types that produce growth-promoting or inhibitory molecules following SCI, and dissect cell-type-specific responses to potential therapies that target these molecules.

### Immunomodulation does not confer neuroprotection after SCI

Early therapeutic approaches to SCI sought to inhibit immune responses to injury, with the aim of conferring neuroprotection^71^. Preclinical studies suggested neuroprotective actions of methylprednisolone^70^ and minocycline^73–75^, which led to large-scale clinical trials^71, 76^. However, these trials failed to demonstrate the effectiveness of these treatments to mediate functional recovery^70, 72, 143^.

We asked whether our single-cell atlas could reconcile the paradox between the established immunomodulatory activity of these drugs and their failure to ameliorate neurological function. Cell type prioritization^60, 127^ confirmed that both methylprednisolone and minocycline triggered a profound transcriptional perturbation of the entire immune lineage (**Supplementary Fig. 16a-b**). However, this immunomodulation did not coincide with an increase in the survival of neurons (**Supplementary Fig. 16c-e**). This failure to protect neurons from apoptosis coincided with the lack of detectable neurological recovery, despite high-resolution kinematic analyses.

Although these agents failed to improve the survival of neurons after SCI, we reasoned that they might induce more subtle transcriptional changes in the surviving neurons. Specifically, we hypothesized that these agents might repress the molecular programs activated by injury within neurons, thus promoting a shift towards an uninjured transcriptional phenotype. To test this hypothesis, we trained a machine-learning model to classify individual neurons as originating from injured versus uninjured spinal cords (**Supplementary Fig. 16f**). Applying this model to cells from mice treated with methylprednisolone or minocycline exposed a threefold increase in the proportion of neurons that were classified as uninjured (**Fig. 3j**). Consistent with this prediction, surviving neurons treated with methylprednisolone downregulated transcriptional programs associated with innate and adaptive immune responses and cellular stress (**Supplementary Fig. 16g-h**).

Together, these findings suggest that methylprednisolone and minocycline modulate the immune responses to SCI, which in turn shifts the surviving neurons towards their basal transcriptional states. However, these agents fail to alter the early, dichotomous outcome of survival versus apoptosis, and therefore fail to prevent neuronal death or improve neurological recovery.

### Sexually dimorphic responses to SCI are subtle

Sexual dimorphism in immune responses underlies differences in the prevalence of autoimmune disease between males and females^66, 67^. Consequently, we hypothesized that transcriptional programs activated by SCI may also be sexually dimorphic. We applied cell type prioritization to evaluate whether specific cell types are differentially perturbed by SCI in male versus female mice. As anticipated, this analysis prioritized immune cell types as having the most prominent sex-dependent transcriptional differences (**Supplementary Fig. 17a**). Moreover, comparison of immune cell proportions revealed an increase in NK cells within female mice, consistent with established sex differences in the adaptive immune response to neurotrauma^66, 144^ (**Supplementary Fig. 17b-c**).

Despite these dimorphic immune responses, however, high-resolution kinematics failed to demonstrate any sexual dimorphism in neurological recovery (**Supplementary Fig. 3f**). Moreover, the average magnitude of the transcriptional perturbation between male and female mice was among the most subtle in our single-cell atlas (**Supplementary Fig. 17d**). Accordingly, we failed to detect sex differences in the overall proportion of surviving neurons (**Supplementary Fig. 17e-f**). Taken together, these results suggest that sexual dimorphism does not impact early neuronal death or survival, and consequently, has no detectable influence on neurological recovery^145^.

### Cellular divergence between animal models of SCI

Preclinical studies of SCI require the selection of a relevant paradigm from a large repertoire of potential injury models^146–148^. One important difference between these models is whether they explicitly open the meninges, which is thought to promote excessive immune cell infiltration. Unexpectedly, however, we found that the degree of peripheral immune invasion was broadly conserved across the injury models included in our single-cell atlas (**Fig. 3k** and **Supplementary Fig. 18a-c**). We validated this finding by morphometrically quantifying Cd45-expressing cells, finding the number of these to be essentially identical across injury models (**Supplementary Fig. 18d-e**). These observations suggest that crush and contusion injuries dismantle the blood-brain barrier and cause extensive peripheral immune invasion that is not contingent on explicit meningeal disruption caused by hemisection injuries.

A second potential difference between preclinical paradigms is their relevance to human injuries. The most common mechanism of spinal cord damage in humans occurs through burst fractures and distraction injuries that impact the ventral spinal cord^149^. To understand whether our profiled injury models lead to differential perturbations of neurons along the dorsoventral axis, we employed cell type prioritization to quantify transcriptional responses in each neuronal subpopulation. Compared to crush injury, we found that dorsal hemisection and contusion injuries preferentially perturbed neurons in the dorsal spinal cord (**Supplementary Fig. 18f**). Conversely, increasingly severe crush injuries induced balanced perturbations in neurons located in the dorsal and ventral spinal cord (**Supplementary Fig. 18g**). These differences between rodent models of SCI must be considered when selecting an injury paradigm for preclinical studies.

### Catastrophic failure of tripartite barrier formation in old mice

Aging causes multifaceted changes in gene expression that culminate in dysregulated transcriptional responses to disease and biological perturbations^150–155^, but whose cellular and functional consequences after SCI remain poorly understood. Remarkably, we found that the transcriptional differences between young and old mice after SCI were nearly as profound as those between injured and uninjured mice (**Fig. 4a**). The magnitude of this transcriptional perturbation was mirrored by extensive functional impairments in old mice compared to young mice (**Fig. 4b-c** and **Supplementary Fig. 3e**). We therefore sought to elucidate the mechanisms underlying these transcriptional and functional differences.

**Fig. 4.**
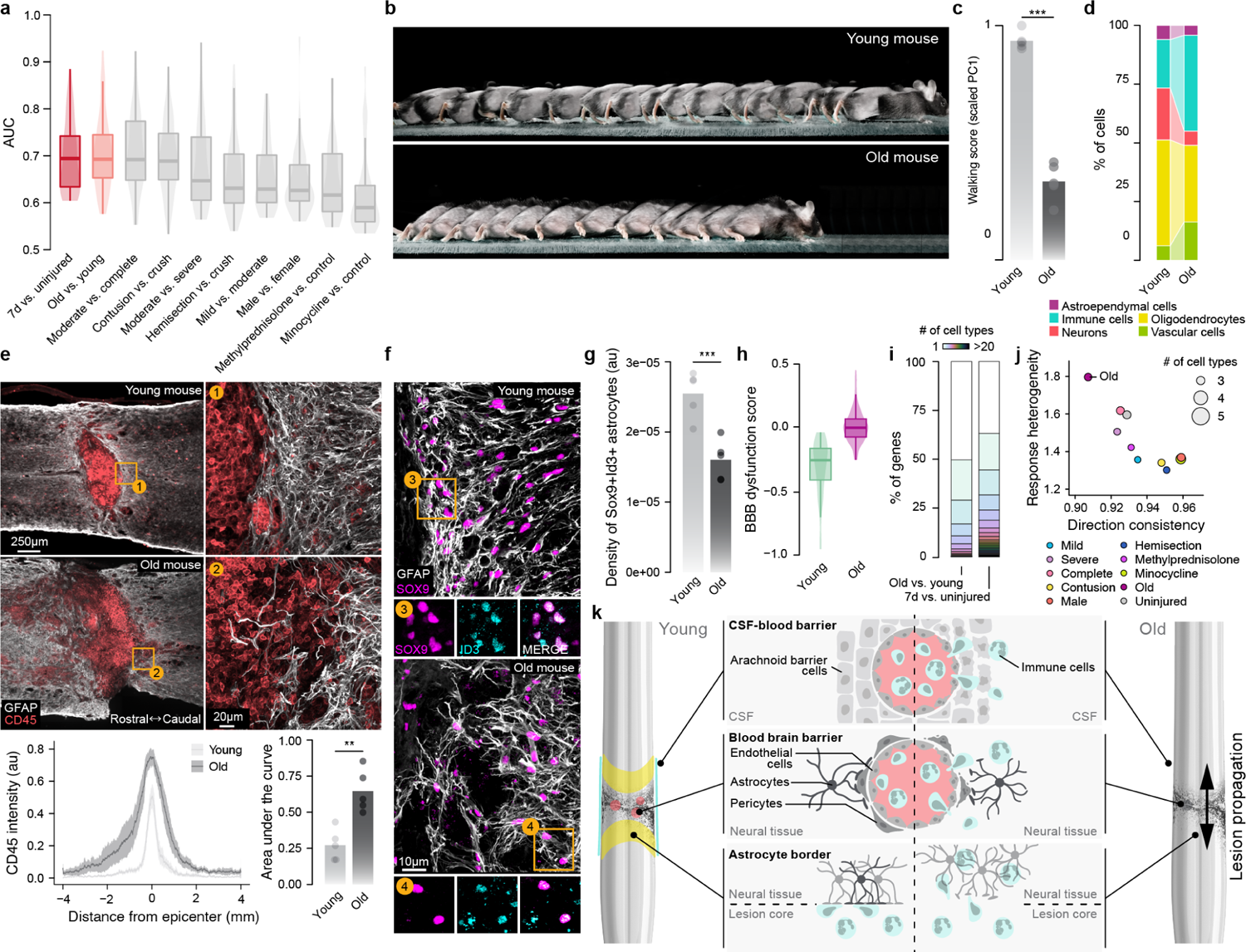
Catastrophic failure to reestablish a tripartite neuroprotective barrier in old mice. **a**, Intensity of the transcriptional perturbation across all cell types of the spinal cord, as quantified by Augur, for all comparisons involving the injured spinal cord at 7 days post-injury. Comparisons between injured and uninjured mice, and old and young injured mice, are highlighted. **b-c,** Chronophotography, **b,** and walking performance (*n* = 5 each), **c,** of young and old mice after spontaneous recovery from moderate crush SCI. **d,** Proportions of each major spinal cord cell type in young versus old mice after SCI. **e,** Composite tiled scans of GFAP and CD45 in horizontal sections from representative old and young mice. Bottom left, line graph demonstrates CD45 intensity at specific distances rostral and caudal to lesion centers. Bottom right, bar graph indicates the area under the curve (independent samples two-tailed t-test; *n* = 5 per group; t = 4.57; p = 0.002). **f,** Horizontal sections from representative old and young mice identifying a lack of Sox9^ON^Id3^ON^ cells in the astrocyte border region in old mice compared to young mice. **g,** Density of Sox9^ON^Id3^ON^ cells in the astrocyte border region (independent samples two-tailed t-test; *n* = 5 per group; t = 4.84; p = 0.001). **h,** Expression of the BBB dysfunction module in capillary endothelial cells from young and old mice at seven days post-injury. **i,** Cell type specificity of DE genes in comparisons of young vs. old mice or injured vs. uninjured mice. **j,** Heterogeneity of cell-type-specific differential expression in experimental comparisons involving the injured spinal cord at seven days after SCI. Aging is characterized by greater discoordination of gene expression than any other condition in the snRNA-seq atlas, as reflected by increased response heterogeneity and decreased direction consistency. **k,** Schematic overview of the cell types comprising the tripartite neuroprotective barrier.

We first compared the proportions of cell types in the injured spinal cord of young and old mice. This comparison revealed a profound reduction in the proportion of neurons surviving the injury in old mice, which was accompanied by a dramatic increase in the proportion of immune cells (**Fig. 4d**). Anatomical assessments confirmed that old mice developed larger lesions compared to young mice, despite identical mechanisms of injury (**Fig. 4e**).

We next asked how age impacted the transcriptional responses to SCI within individual cell types. Cell type prioritization^60, 127^ revealed abnormal responses in infiltrating immune cells from old mice, including dividing myeloid progenitors, NK cells, and T cells (**Supplementary Fig. 19a**). However, Augur also detected abnormal responses within cell types involved in the formation of the blood-brain barrier and the astrocyte barrier, including extracellular matrix-forming VLMCs, capillary endothelial cells, and OPCs. Consistent with this observation, we detected an age-dependent decrease in the proportion of *Id3*-expressing astrocytes, which form the astrocyte lesion border^156^, and of arachnoid barrier cells, which form the cerebrospinal fluid barrier (**Supplementary Fig. 19b-c**). Immunohistochemistry for Sox9 and Id3 confirmed these findings (**Fig. 4f,g**). Within the vascular compartment, we identified an age-dependent upregulation of gene programs associated with dysfunction of the blood-brain barrier, and downregulation of the specialized gene programs that enable vascular cells to establish the blood-brain barrier (**Fig. 4h** and **Supplementary Fig. 19d**).

To dissect the transcriptional programs that are dysregulated in old mice, we performed DE analysis^87^ of all the cell types in the spinal cord. In contrast to young mice, we observed that many genes were DE within just a single cell type in old mice (**Fig. 4i**). Moreover, other genes were upregulated in some cell types, but downregulated in others (**Supplementary Fig. 19e-f**). To quantify the coordination of the transcriptional responses to SCI across cell types, we devised statistical measures that aimed to capture both the variability of DE and changes in the direction of DE across cell types. These quantifications revealed that transcriptional responses to injury were profoundly discoordinated across the cell types of the spinal cord in old mice, relative to every other experimental comparison (**Fig. 4j** and **Supplementary Fig. 19e-f**).

Together, these findings suggest that old mice fail to deploy the coordinated, multicellular response to SCI that naturally occurs in young mice. This failure manifests in the disruption of three essential neuroprotective barriers between the immune-privileged and extra-neural environments of the injured spinal cord: (i) the blood-brain barrier; (ii) the CSF-brain barrier; and (iii) the border-forming astrocyte barrier (**Fig. 4k**). The consequence of this tripartite barrier formation failure is a dramatic increase in peripheral immune cell invasion, which culminates in the uncontrolled expansion of the lesion, the catastrophic loss of neurons adjacent to the injury site, and a resulting inability to coordinate the recovery of neurological functions.

### A single-nucleus multi-omic atlas of the injured and uninjured spinal cord

Our snRNA-seq atlas exposed the transcriptional programs triggered by SCI across the entire repertoire of cells in the spinal cord. However, we recognized that this atlas was intrinsically limited in its ability to expose the regulatory mechanisms that underlie these transcriptional programs. To overcome this limitation, we compiled the second atlas of the *Tabulae Paralytica*: a multi-omic atlas of the injured spinal cord (**Fig. 5a**).

**Fig. 5.**
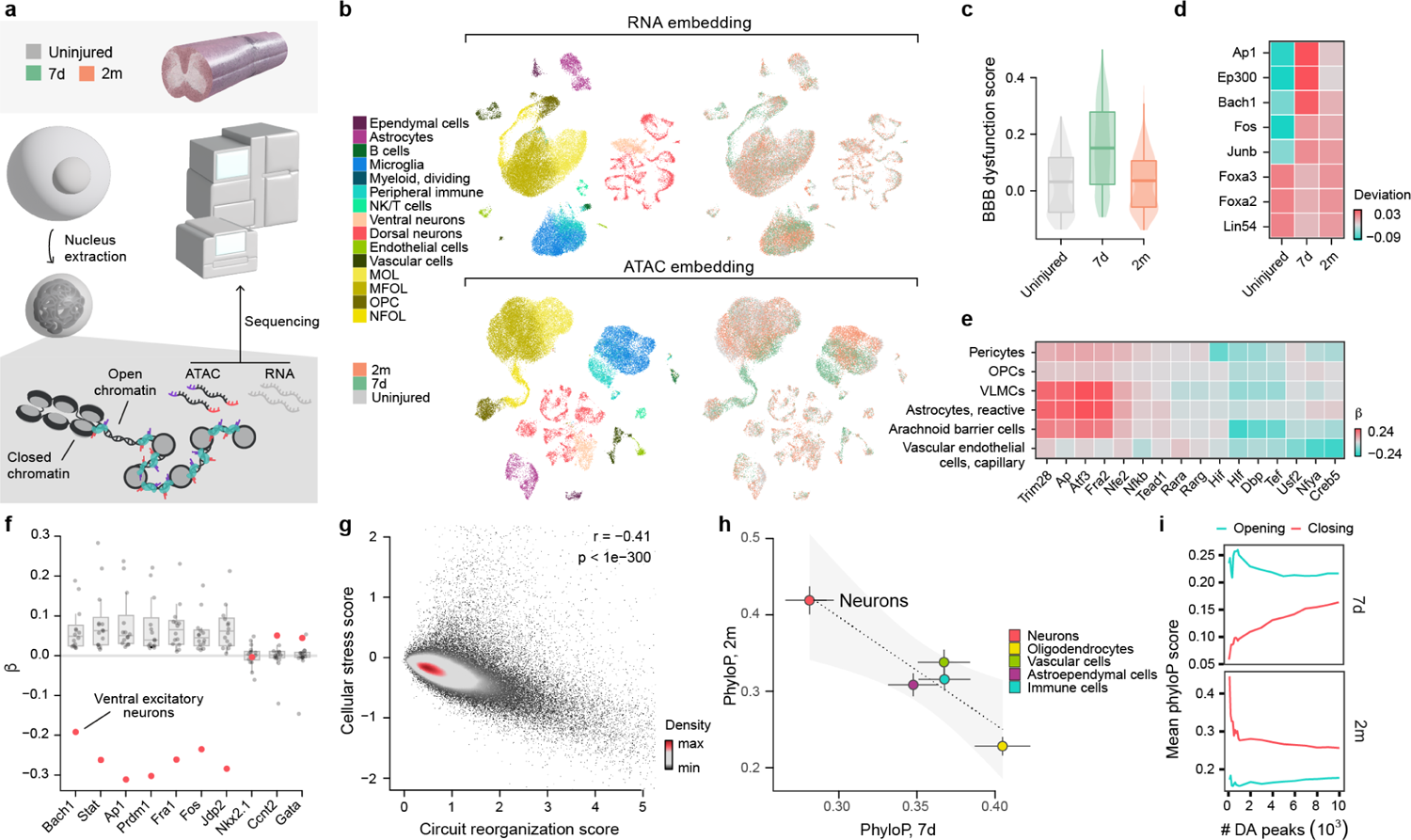
A multi-omic atlas of SCI. **a**, Schematic overview of the multiome atlas. **b,** UMAP visualizations of the 40,256 cells in the multiome atlas, based on the RNA, top, or ATAC, bottom, modalities, and colored by cell types, left, or experimental conditions, right. **c,** Expression of the blood-brain barrier dysfunction module score in vascular cells across timepoints in the multiome atlas. **d,** Transcription factors for which motif accessibility in the ATAC modality correlated with expression of the blood-brain barrier dysfunction module in the RNA modality across vascular cells in the multiome atlas. **e,** Selected transcription factors with significant differences in motif accessibility at 7 days in cell types that establish the tripartite barrier. **f,** Transcription factors exhibiting significant variability in transcription factor binding across neuron subtypes at 2 months. Motif accessibility in ventral excitatory interneurons is highlighted. **g,** Anticorrelated expression of the circuit reorganization module, x-axis, versus that of the cellular stress module, y-axis, in 80,315 neurons from the snRNA-seq atlas. **h,** Mean phyloP scores of differentially accessible peaks in major cell types of the spinal cord at 7 days, x-axis, and 2 months, y-axis. Error bars show the standard error of the mean for each comparison. **i,** Mean phyloP scores of differentially accessible peaks that are opening or closing in neurons at 7 days or 2 months post-injury.

We deployed single-nucleus multi-omics to measure both RNA and accessible chromatin within the same individual cells, using the assay for transposase-accessible chromatin by sequencing (ATAC-seq). We leveraged these methodologies to profile the uninjured and injured spinal cords of mice at 7 days and 2 months post-injury. After quality control of both modalities^157^, we obtained a dataset measuring gene expression and chromatin accessibility in 47,726 nuclei (**Fig. 5b** and **Supplementary Fig. 20**).

We aimed to link the multi-omic atlas to the cellular taxonomy of the spinal cord that our snRNA-seq atlas had established. To overcome challenges in cell type annotation within snATAC-seq data, we adapted an automated cell type annotation approach^158^ to hierarchically assign cell types and subtypes to each cell in the multiome atlas based on the RNA modality. We validated the accuracy of this approach through cross-validation in the snRNA-seq atlas, and established that cell types were recovered at similar frequencies in both atlases (**Supplementary Figs. 21-22**).

We then leveraged this taxonomy to call peaks within each cell type and subtype at increasingly granular resolutions on the clustering tree (**Supplementary Fig. 23**)^157^, and identified differentially accessible transcriptional factors within each subpopulation using chromVAR (**Supplementary Figs. 24-25**)^159^. This analysis highlighted well-studied transcriptional factors that play canonical roles in specifying the identity of spinal cord cell types, including astrocytes (*Nr1d1*, *Nfib*^160^), B cells (*Pax2*, *Pax5*^161^), Microglia (*Irf1*^162^), pericytes (*Ebf1*^163^), and endothelial cells (*Lef1*^164^). However, our data also pointed to less-characterized cell-type-specific TFs, including *Zbtb3* (ventral interneurons), *Nf1* (vascular leptomeningeal cells), *Zbtb7b* (oligodendrocyte precursor cells), *Rfx5* (ependymal cells), *Ccnt2* (CSF-contacting neurons), and *Mlx* (immune cells).

### Regulatory programs underlying tripartite neuroprotective barrier formation

Since our snRNA-seq atlas exposed a number of biological principles that dictate the multifaceted responses to SCI in different cell types of the spinal cord, we sought to leverage our multiome atlas to understand the regulatory programs that orchestrate these transcriptional responses.

We first aimed to dissect the gene regulatory programs involved in the reestablishment of the tripartite neuroprotective barrier after SCI. Because our multiome atlas recapitulated the upregulation of gene programs associated with blood-brain barrier dysfunction that we had observed in the snRNA-seq atlas (**Fig. 5c**), we leveraged the ATAC modality to identify the transcriptional factors that underlie this dysfunction. To exploit the link between RNA and ATAC modalities, we correlated the accessibility of transcriptional factor binding motifs to the expression of these gene programs within the same cell (**Supplementary Fig. 26a**). Within vascular cells, the expression of the blood-brain barrier dysfunction program^101^ was correlated with the accessibility of transcription factors associated with cellular stress and inflammation (*Ap1*, *Junb*, *Bach1*, *Fos*) and hypoxia-induced VEGF stimulation (*EP300*), and anticorrelated with the accessibility of transcription factors driving cellular proliferation (*Foxa2*, *Foxa3*, *Lin54*; **Fig. 5d** and **Supplementary Fig. 26b**).

These responses were mirrored by shared and cell-type-specific regulatory programs in the other cellular subpopulations that coordinate the formation of the tripartite barrier (**Fig. 5e** and **Supplementary Fig. 26c**). VLMCs, pericytes, and arachnoid barrier cells exhibited decreased accessibility of transcription factors known to modulate the permeability of the blood-brain or cerebrospinal fluid barriers (*Rarg*, *Hif1*), whereas arachnoid barrier cells and pericytes exhibited decreased accessibility of transcription factors that govern barrier efflux of metabolites and that regulate neuronal activity (*Dbp*, *Tef*). Finally, border-forming astrocytes exhibited increased motif accessibility of multiple transcription factors associated with acute responses to stress or hypoxia, including *Junb*, *Bach1*, *Fos*, and *EP300*^160^.

### Evolutionary divergence in circuit reorganization

Our snRNA-seq atlas established that SCI triggers an immediate transcriptional response that is conserved across all neuronal subpopulations. Conversely, we found that transcriptional responses gradually diverged between neuronal sub-populations over time after SCI (**Fig. 3c**). We therefore next sought to understand the regulatory programs that govern neuronal responses to SCI as well as their associated functional consequences and potential origins.

In the multiome atlas, we observed that the early-conserved transcriptional response was mirrored by conserved regulatory programs that involved increased accessibility of transcription factors associated with cellular stress (*Myc*, *Nfe2*) and apoptosis (*Tfap2*) (**Supplementary Fig. 27a**). To characterize late-diverging regulatory programs, we devised a permutation-based statistical approach that evaluated variability in transcription factor binding across all subpopulations of neurons (**Supplementary Fig. 27b**). We found that this variability originated from divergent regulatory responses within a subpopulation of ventral excitatory interneurons (**Fig. 5f** and **Supplementary Fig. 27c**). Whereas every other subpopulation of neurons exhibited increased accessibility of transcription factors associated with cellular stress responses (*Ap1*, *Stat*, *Bach1*, *Fos*), this subpopulation of ventral excitatory interneurons instead exhibited decreased accessibility of these transcription factors.

The results from our snRNA-seq atlas revealed that ventral excitatory interneurons express genes implicated in circuit reorganization at high levels (**Supplementary Fig. 13c**). The distinctive lack of cellular stress responses within these neurons led us to hypothesize that, in general, neurons face an inherent trade-off between the expression of cellular stress response programs and transcriptional programs associated with circuit reorganization. To test this hypothesis, we reexamined our snRNA-seq atlas and confirmed the existence of an anticorrelation between the expression of programs related to stress response versus circuit reorganization (**Fig. 5g**). These observations suggest a model whereby the ability of different neuronal subpopulations to participate in circuit reorganization is intrinsically linked to the intensity of their response to cellular stress.

Neuronal responses to injury vary dramatically across the tree of life, to the extent that neurons from lower vertebrates can demonstrate spontaneous regeneration whereas neurons from adult mammals fail to regenerate after SCI^16^. This divergence compelled us to characterize the evolutionary conservation of the genomic regions that become differentially accessible following SCI. We used phyloP^165^ to quantify the sequence conservation of these regions, and identified profound differences in the evolutionary conservation of differentially accessible peaks within neurons, as compared to glia (**Fig. 5h** and **Supplementary Fig. 28a-c**). Inspecting these differences more closely, we discovered dichotomous patterns of evolutionary conservation for peaks that opened versus closed in neurons after SCI (**Fig. 5i** and **Supplementary Fig. 28d-e**). In the acute phase of SCI, neurons displayed increased accessibility of evolutionarily conserved genomic regions, which was mirrored by decreased accessibility of evolutionarily accelerated genomic regions. These trends were reversed two months after SCI, when evolutionarily accelerated regions displayed increased accessibility.

These observations led us to ask whether the opening of evolutionarily accelerated regions is necessary for neurons to participate in the recovery of neurological functions. To test this possibility, we computed the mean evolutionary conservation of accessible genomic regions within individual neurons. Across all subpopulations of neurons, we found that the same subpopulation of ventral excitatory interneurons demonstrated the most pronounced opening of evolutionarily accelerated regions in response to injury (**Supplementary Fig. 28f**).

We conclude that neurons in the spinal cord face an inherent tradeoff between the activation of cellular stress responses, and the opening of evolutionarily accelerated genomic regions to express transcriptional programs associated with circuit reorganization.

### A spatial transcriptomic atlas of SCI

Interrogation of our snRNA-seq and multiome atlases identified cell-type-specific transcriptional and regulatory programs triggered by SCI. However, these transcriptional and regulatory programs were identified in dissociated cells, and therefore, could not be visualized within the complex microenvironment of the injury. To overcome this limitation, we resolved these programs within the cytoarchitecture of the spinal cord using spatial transcriptomics (**Fig. 6a**).

**Fig. 6.**
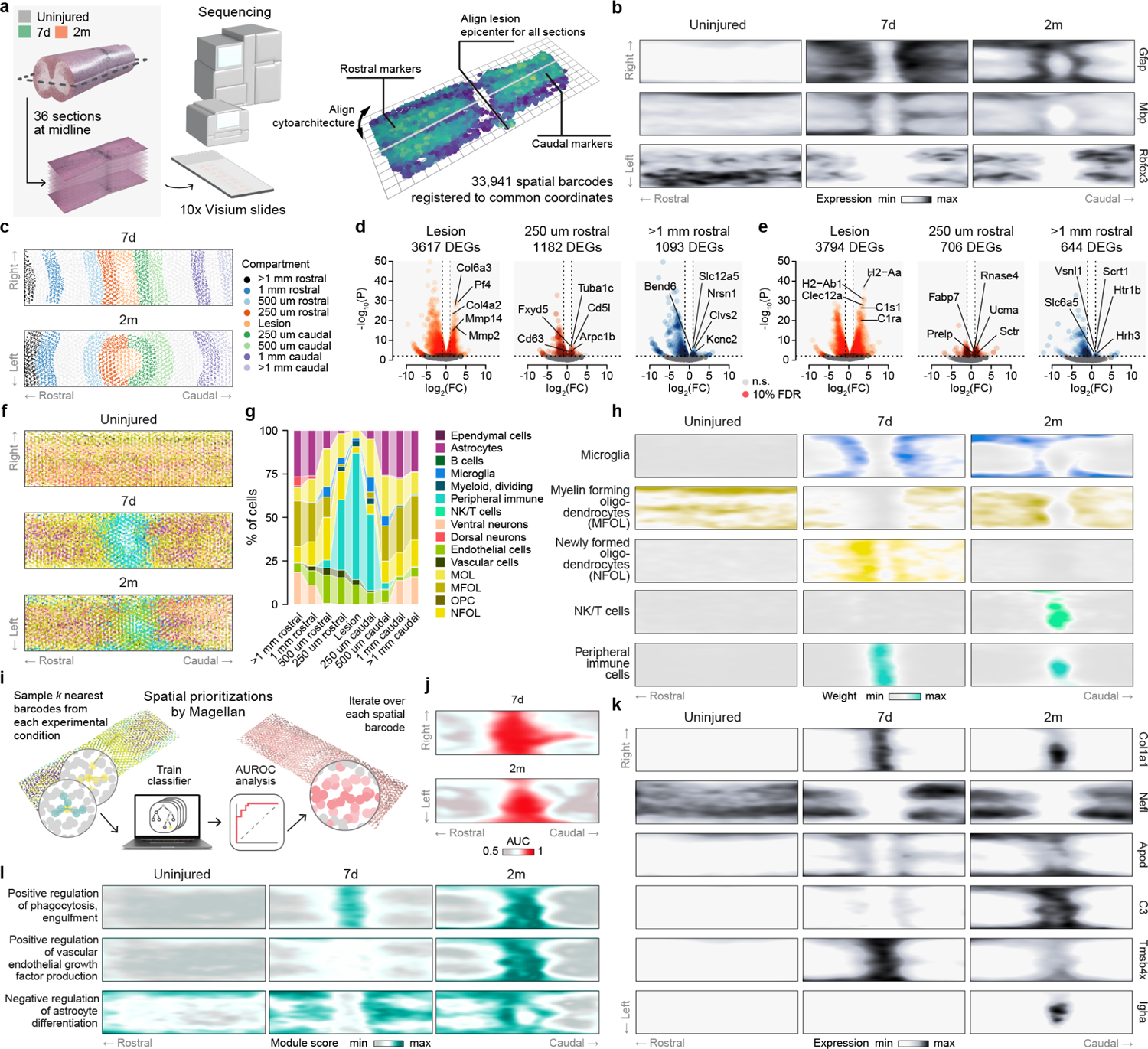
A spatial transcriptomic atlas of SCI. **a**, Schematic overview of the 2D spatial transcriptomic atlas. **b,** Computational histology of key marker genes on the two-dimensional common coordinate system of the spinal cord. **c,** Legend showing the position of each lesion compartment on the coordinate system of the spinal cord. **d-e,** Volcano plots showing compartment-specific genes at 7 days, **d,** and 2 months, **e,** for a subset of the lesion compartments discussed in the text. **f,** Cell type deconvolution of spatial barcodes in the 2D spatial transcriptomic atlas. **g,** Cellular composition of each lesion compartment, as determined by cell type deconvolution. **h,** Deconvolution weights assigned by RCTD to selected cell types. **i-j,** Spatial prioritizations assigned by Magellan to each spatial barcode at 7 days, **i,** and 2 months, **j. k-l,** Expression of selected genes, **k,** and gene modules, **l,** prioritized by their correlation to spatial prioritizations assigned by Magellan at each spatial barcode.

We profiled the spinal cords of uninjured and injured mice at 7 days and 2 months after SCI, and obtained 33,941 high-quality spatial barcodes from 36 coronal sections (**Supplementary Fig. 29**). To permit direct comparison across experimental conditions, we registered all 36 sections to a common coordinate system^166^ (**Fig. 6a-b**).

The coordinated, multicellular response to SCI establishes a thin astrocyte barrier that separates the fibrotic lesion core from surrounding immune-privileged neural tissue^17, 18^. The requirements to promote neural repair are known to differ between these distinct lesion compartments, but the underlying molecular logic remains incompletely understood^2, 10^. We leveraged our spatial atlas to uncover the molecular programs that are shared between, or specific to, each of these compartments.

To identify molecular differences between lesion compartments, we demarcated spatial barcodes corresponding to the fibrotic scar, the astrocyte barrier, and the adjacent neural tissue (**Fig. 6c** and **Supplementary Fig. 30**). We then performed DE analysis to identify genes specific to each lesion compartment (**Fig. 6d-e** and **Supplementary Fig. 31**). This analysis showed that the fibrotic scar was differentiated from other lesion compartments by upregulation of extracellular matrix molecules (*Col6a3*, *Col4a2*) and matrix metalloproteinases (*Mmp2*, *Mmp14*), and a sustained expression of cytokines (*Cd74*, *Pf4*) and complement proteins (*C1s1*, *C1ra*). The astrocyte barrier was differentiated by genes associated with cytoskeletal organization (*Tuba1c*, *Arpc1b*) and cell proliferation (*Fxyd5*, *Id3*) at 7 days, and by genes related to radial glial fiber development (*Fabp7*), extracellular matrix molecules (*Ucma*, *Prelp*), synaptic reorganization (*Sctr*), and neuronal survival (*Rnase4*) at 2 months after SCI. Spared reactive neural tissue was differentiated by genes associated with neuronal functions, including solute transport (*Slc12a5*, *Kcnc2*), neuron growth and projection formation (*Nrsn1*, *Bend6*), and synapse maturation (*Scrt1*, *Nptx1*), whereas levels of growth factors and their receptors demonstrated little change.

We next sought to dissect the cellular composition of each lesion compartment by deconvolving the cell types within each spatial barcode^167^ (**Fig. 6f-h** and **Supplementary Fig. 32**). We found that the fibrotic lesion core was composed almost exclusively of immune and vascular cells (**Fig. 6g**). The density of astroependymal cells abruptly increased towards the edge of the lesion, and thereafter gradually decreased with increasing distance from the lesion core. As the distance from the lesion core increased, the proportions of each cell type approximated those observed in uninjured spinal cords, mirroring immunohistochemical observations^17, 18^.

To quantify the relative degree of transcriptional perturbation throughout the lesion microenvironment, we applied Magellan (**Fig. 6i**)^9^. Magellan is a machine-learning framework that quantifies the relative magnitude of the transcriptional response at any given spatial locus to an arbitrary perturbation, a procedure we refer to as spatial prioritization (**Supplementary Fig. 33a**). This prioritization recovered the profound transcriptional perturbation occurring at the lesion core during the first 7 days after injury, with a gradient of decreasing intensity that spread radially throughout the spared but reactive neural tissue adjacent to the fibrotic scar (**Fig. 6j** and **Supplementary Fig. 33b**). Spatial prioritization also captured the contraction of the injury border after 2 months of recovery from SCI (**Fig. 6j** and **Supplementary Fig. 33c-d**).

These results illustrate how spatial prioritization accurately recovered the two-dimensional architecture of the evolving injury. We therefore asked whether spatial prioritization could also provide a resource to identify the molecular programs that elaborate this architecture, without any *a priori* definition of the lesion compartments. To answer this question, we tested for correlation between the spatial prioritization scores assigned to each barcode by Magellan and the expression of individual genes (**Supplementary Fig. 34a-c**). As anticipated, this approach recovered many of the genes associated with canonical lesion compartments, including extracellular matrix molecules at the lesion core (*Col1a1*, *Col13a1*) and neuronal genes (*Nefl*, *Nefh*) in spared but reactive neural tissue (**Fig. 6k** and **Supplementary Fig. 34d-f**). Similarly, we tested for correlation between the spatial prioritizations assigned by Magellan and the average expression of all genes associated with a given Gene Ontology (GO) term. This analysis recapitulated the multifaceted innate and adaptive immune responses within the lesion site (**Fig. 6l** and **Supplementary Fig. 35**).

We then asked whether the spatial transcriptomic atlas could identify genes whose correlation to the perturbation response differed between 7 days and 2 months after SCI. To answer this question, we tested for differential correlation between spatial prioritization scores and gene expression at 7 days and 2 months post-SCI (**Supplementary Fig. 36a**). This analysis highlighted genes (*Gfap*, *Aqp4*, *Apod*) coinciding with the location of the astrocyte barrier, reflecting the contraction of this border that takes place between 7 days and 2 months after injury (**Fig. 6k** and **Supplementary Fig. 36b**). Beyond the astrocyte barrier, differential prioritization also pointed to temporal evolution in the innate and adaptive immune responses, including immediate microglial activation (*Wfdc17*, *Spp1*) and lymphocyte homing (*Stab1*), which contrasted with delayed complement activation (*C3*) and monocyte maturation (*Ms4a7*) (**Fig. 6k** and **Supplementary Fig. 36b**). Repeating this differential prioritization at the level of GO terms highlighted the spatial evolution of astrocyte differentiation, vascular endothelial growth factor production, and phagocytosis (**Fig. 6l** and **Supplementary Fig. 37**).

Spatial prioritization also allowed us to uncover less appreciated aspects of the biology of an SCI. For example, we identified chronic activation of immunoglobulin factors (*Ighg2c*, *Jchain*, *Igha*) within the lesion core, which likely contribute to maintaining host defenses, and antigen binding within the fibrotic core (**Supplementary Fig. 36b**). Spatial prioritization also highlighted a robust expression of *Dbi*^168^ along the lesion border (**Supplementary Fig. 36b**). Since *Dbi* modulates the activity of the neurotransmitter gamma-aminobutyric acid (GABA), the expression of this gene may be involved in the reported reduction of neuronal activity in the vicinity of lesion borders. Moreover, our analyses identified *Prdx6* as highly associated with the lesion border, suggesting the expression of this antioxidant enzyme may protect the surrounding reactive neural tissue from oxidative injury (**Supplementary Fig. 36b**). Last, spatial prioritization identified distinct subcompartments of the lesion core itself. We observed that genes associated with iron metabolism (*Flt1*) expressed diffusely throughout the lesion, but genes associated with fat metabolism (*Plin2*) confined to the innermost aspects of the lesion core, and genes associated with actin sequestration (*Tms4bx*) extending out along the lesion edges (**Fig. 6k** and **Supplementary Fig. 34f**).

Together, these results establish a resource to explore the multicellular responses to SCI across the cytoarchitecture of the spinal cord, and validate the ability of spatial prioritization to resolve both well-documented and novel aspects of these responses.

### A four-dimensional spatiotemporal atlas of SCI

The ability to visualize the central nervous system in three dimensions using tissue clearing technologies has opened new possibilities to study the anatomy and function of the nervous system^169^. Analogously, we reasoned that expanding our spatial transcriptomic atlas into a third spatial dimension would enable a more complete description of the biology of SCI. We further surmised that our snRNA-seq and multiome atlases could be overlaid onto this three-dimensional model of the spinal cord in order to resolve the spatiotemporal distribution of the transcriptional and regulatory responses across the compendium of experimental conditions included in our *Tabulae*.

To develop a four-dimensional spatial atlas of the spinal cord, we collected 16 tissue sections that were equally spaced along the dorsoventral axis of spinal cords from uninjured and injured mice at 7 days and 2 months after SCI (**Fig. 7a**). The distance between each section was approximately 50 *µ*m, which ensured a dense coverage of the third spatial dimension. After quality control and registration to a common three-dimensional coordinate system, we obtained a dataset comprising 37,558 spatial barcodes from 48 sections (**Supplementary Fig. 38**).

**Fig. 7.**
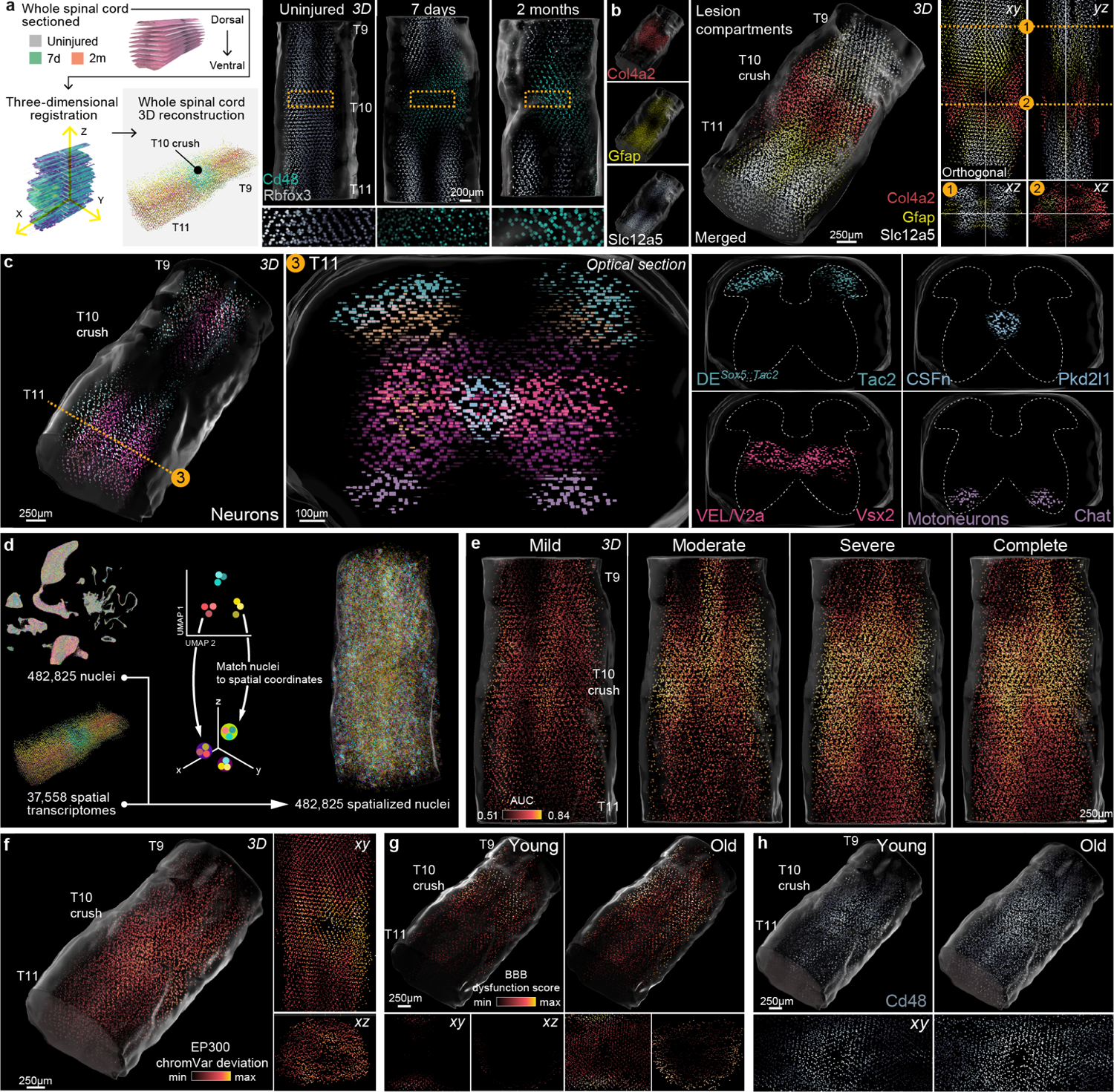
A four-dimensional spatiotemporal atlas of SCI. **a**, Left, schematic overview of the 3D spatial transcriptomic atlas. Right, expression of marker genes for neurons (*Rbfox3*) and immune cells (*Cd48*) across three dimensions in the uninjured and injured mouse spinal cord. **b,** Expression of marker genes associated with distinct lesion compartments across the 3D spatial transcriptomic atlas at 7 days, including the fibrotic scar (*Col4a2*), the astrocyte barrier (*Gfap*) and the surrounding spared but reactive neural tissue (*Slc12a5*). **c,** Spatial localization of selected neuronal subpopulations defined by snRNA-seq across the 3D spatial transcriptomic atlas at 7 days. **d,** Left, UMAP representation of 435,099 single-nucleus transcriptomes from the snRNA-seq atlases, colored by experimental condition. Right, spatial coordinates assigned to each single-nucleus transcriptome within the 3D spatial transcriptomic atlas. **e,** Three-dimensional spatial prioritization of spatialized cells from the snRNA-seq atlas, across injury severities. **f,** Accessibility of the EP300 binding motif within spatialized cells from the multiome atlas at 7 days, visualized on the 3D spatial transcriptomic atlas. **g,** Expression of the blood-brain barrier dysfunction module in vascular cells from young and old mice at 7 days, visualized on the 3D spatial transcrip-tomic atlas. **h,** Expression of *Cd48* in spatialized cells from young and old mice at 7 days, visualized on the 3D spatial transcriptomic atlas.

To validate the construction of our four-dimensional atlas, we first confirmed that our dataset resolved the established spatial distributions of cell types in the uninjured and injured spinal cord^17, 18^. Three-dimensional projections of gene expression exposed the distinct lesion compartments, including the fibrotic scar (*Col4a2*), the astrocyte barrier (*Gfap*), and spared but reactive neural tissue (*Slc12a5*; **Fig. 7b** and **Supplementary Fig. 39**). Moreover, neurons and immune cells occupied their appropriate locations in the injured spinal cord, whereby the fibrotic scar was invaded by peripheral immune cells and devoid of neurons. We next inspected the expression of well-studied inhibitory and facilitating molecules (**Supplementary Fig. 40**), finding the inhibitory CSPG aggrecan (*Acan*) was expressed within spared neural tissue that surrounded the lesion core, whereas brevican (*Bcan*) was expressed in both the lesion core and neural tissue. The growth-permissive molecule *Cspg4* was expressed predominantly in the lesion core, whereas *Cspg5* was expressed within the astrocyte barrier. Finally, the permissive extracellular matrix molecule *Lama1* was upregulated both within the lesion core and by border-forming astrocytes.

To increase the resolution of this spatiotemporal atlas, we again leveraged our snRNA-seq atlas to deconvolve the cellular composition of each spatial barcode (**Supplementary Figs. 41-42**). This procedure resolved cellular subpopulations within highly specific locations, such as ependymal cells and border-forming macrophages. Moreover, dorsal and ventral neurons were appropriately separated along the coronal plane, and the locations of specific neuronal subpopulations such as cerebrospinal fluid-contacting neurons, *Vsx2*-expressing neurons, and motor neurons were correctly resolved (**Fig. 7c**).

We next aimed to integrate all of the *Tabulae* into a single, unified framework. Using Tangram^170^, we embedded single-nucleus transcriptomes and epigenomes onto our four-dimensional atlas of the mouse spinal cord, generating a unified dataset of 554,324 single-nucleus or spatial barcodes that were each associated with a full transcriptome, an experimental condition, and x-, y-, and z-coordinates (**Fig. 7d**).

We then applied Magellan to the integrated spatial dataset. This spatial prioritization reflected the severity-dependent increase in transcriptional perturbation within increasing injury severity (**Fig. 7e**). Consistent with these observations, Magellan captured severity-dependent changes in peripheral immune cell invasion, astrocytic demarcation of the lesion, and neuronal death (**Supplementary Fig. 43**). Moreover, we spatialized the expression of conserved-early neuronal responses, as well as late-diverging expression of programs associated with circuit reorganization (**Supplementary Fig. 44**). We then linked these changes in cell type composition and gene expression to transcription factor accessibility by spatializing the accessibility of transcription factors involved in the establishment of the tripartite barrier (**Fig. 7f** and **Supplementary Fig. 45**).

Our snRNA-seq atlas identified a profound transcriptional perturbation across spinal cord cell types in old mice following SCI. We therefore sought to understand the spatial distribution of this perturbation. Magellan revealed that old mice developed an expanded and poorly circumscribed territory of transcriptional perturbation as compared to young mice, reflecting their failure to re-establish the tripartite neuroprotective barrier. This failure was reflected by global upregulation of blood-brain barrier dysfunction module score, and downregulation of gene expression programs associated with blood-brain barrier identity (**Fig. 7g** and **Supplementary Fig. 46**). Three-dimensional visualization of genes associated with peripheral immune invasion exposed the failure to demarcate the lesion in old mice (**Fig. 7h**). Collectively, these results establish the feasibility of constructing an integrated transcriptomic and epigenomic atlas of healthy and perturbed tissues across four spatiotemporal dimensions.

### Reestablishing the tripartite barrier restores walking in old mice

The *Tabulae Paralytica* documented the spatially- and temporally-dependent activation of transcriptional and regulatory mechanisms that are triggered after SCI in order to reestablish a tripartite neuroprotective barrier. Conversely, the catastrophic failure to reestablish this tripartite barrier in old mice results in poorly circumscribed lesions, massive neuronal death, and impaired recovery of neurological functions (**Fig. 4** and **Supplementary Fig. 19**). These observations led us to hypothesize that interventions that accelerate wound repair by promoting the formation of the tripartite barrier could restore neurological functions in old mice.

Because we found that the number of *Id3*-expressing, border-forming astrocytes was decreased in old mice (**Fig. 4f-g**), we reasoned that increasing their production would accelerate the formation of the astrocyte barrier, limiting lesion size and preserving neurological function. We previously found that the delivery of EGF and FGF2 increased both the proliferation and absolute number of border-forming astrocytes^63^. Moreover, it is established that the delivery of Vegf accelerates endothelial cell proliferation and reformation of vascular networks^171^. We therefore engineered lentiviruses to overexpress Egf, Fgf2, and Vegf and, as a proof-of-principle test, delivered these vectors to the lower thoracic spinal cord two days prior to SCI (**Fig. 8a** and **Supplementary Fig. 47a**). This procedure increased the production of border-forming astrocytes, reduced the number of CD45^+^ infiltrating immune cells, and resulted in smaller and more circumscribed lesions (**Fig. 8b-c** and **Supplementary Fig. 47b-c**). In addition, and remarkably, treated old mice exhibited a natural recovery of walking resembling that of young mice subjected to the same severity of SCI (**Fig. 8d-f** and **Supplementary Fig. 47d-e**). Together, these findings demonstrate that interventions that augment the tripartite neuroprotective barrier and thereby maintain the immune-privileged environment of the spinal cord can prevent the catastrophic neural damage resulting from SCI in aged subjects.

**Fig. 8.**
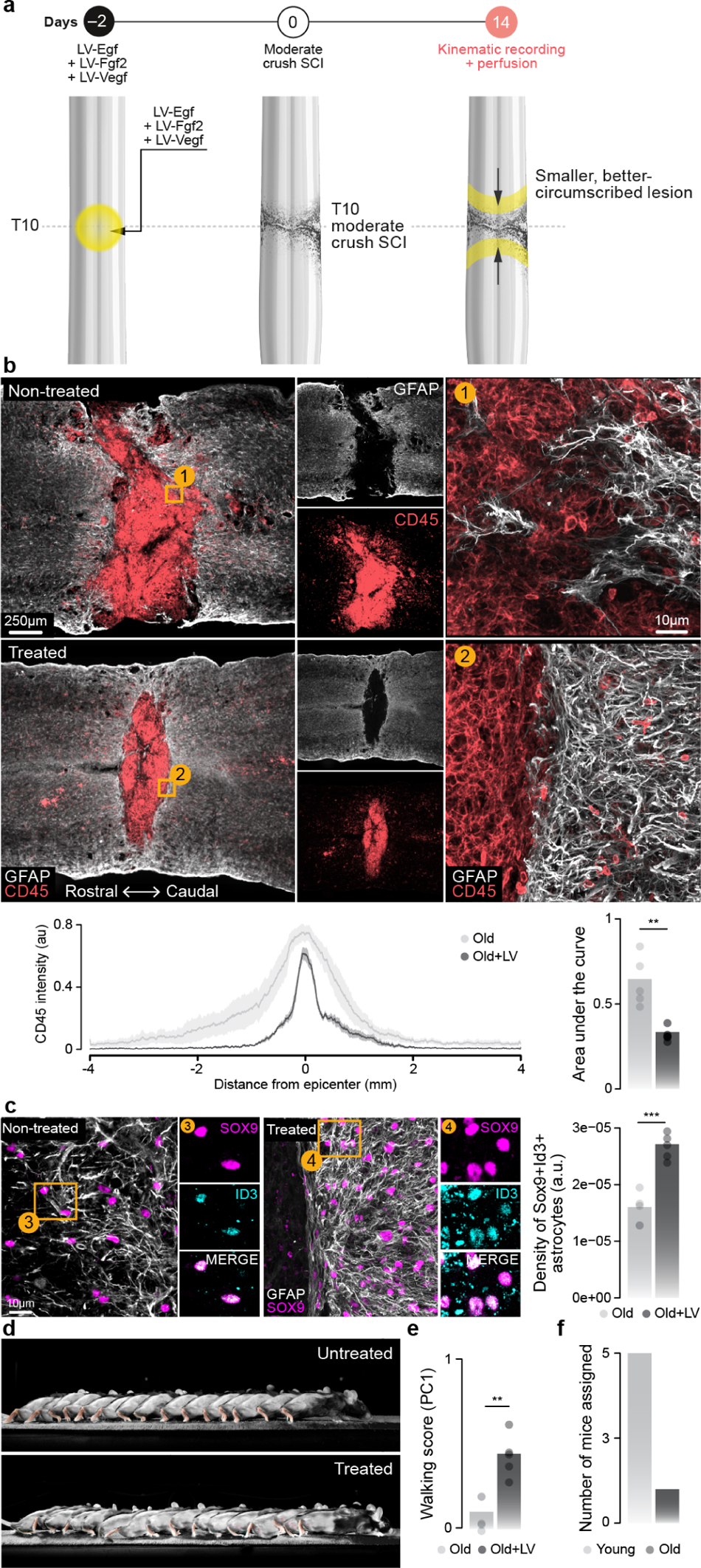
A rejuvenative gene therapy reestablishes the tripartite barrier to restore walking. **a**, Experimental design of a gene therapy intervention to promote the formation of the tripartite barrier. **b,** Composite tiled scans of GFAP and CD45 in horizontal sections from representative old and treated mice. Line graph demonstrates CD45 intensity at specific distances rostral and caudal to lesion centers. Bar graph shows the area under the curve (independent samples two-tailed t-test; *n* = 5 per group; t = 4.57; p = 0.002). **c,** Horizontal sections from representative old and treated mice identifying a restoration of Sox9^ON^Id3^ON^ cells in the astrocyte border region in treated mice. Right, bar graph indicates the density of Sox9^ON^Id3^ON^ cells in the astrocyte border region (independent samples two-tailed t-test; *n* = 5 per group; t = 6.84; p = 0.0002). **d,** Chronophotography of walking in old mice without (top) and with (bottom) a gene therapy intervention to promote the formation of the tripartite barrier. **e,** Walking performance of old mice with and without treatment (independent samples two-tailed t-test; *n* = 5 per group; t = 4.85; p = 0.001). **f,** Experimental conditions assigned to old mice that received gene therapy by a machine-learning model trained on kinematics data from untreated animals. Mice were almost exclusively assigned to the young mouse group, indicating that the walking patterns of treated old mice most resemble those of young mice.

## Discussion

SCI triggers a coordinated cascade of cellular and molecular responses, whose spatiotemporal complexity has thus far prevented the development of safe and effective therapies to repair the injured spinal cord. To help unravel this complexity, we established the *Tabulae Paralytica*—a resource comprising multimodal single-cell and spatial atlases of SCI. We profiled single-nucleus RNA expression in more than 400,000 cells, spanning 18 experimental conditions that captured the most commonly studied manipulations in basic and translational research on SCI that could be made accessible for these experiments. We simultaneously profiled the dynamics of chromatin accessibility and gene expression in a further 40,000 cells to expose the regulatory programs that direct the response to injury. To circumscribe these responses within the cytoarchitecture of the injured spinal cord, we generated a spatial transcriptomic atlas of the injury that we extended into four spatial and temporal dimensions. We merged these atlases into a combined atlas to provide an unprecedented window into the genome-wide molecular cascade that unfolds after an injury to the spinal cord—from epigenetic regulation, to transcriptional programs within individual cells, to spatially- and temporally-dependent multicellular responses—that we overlaid onto a four-dimensional model of the spinal cord.

The *Tabulae Paralytica* provide a resource that not only allowed us to resolve outstanding questions within the field of SCI, but also exposed previously unknown biological principles. For example, we revealed a dichotomy between the early-conserved and late-diverging neuronal responses to SCI, the latter of which reflect differential capacity for circuit reorganization across neuronal subpopulations. We found that this capacity is encoded in the basal transcriptional repertoires of each subpopulation prior to SCI, implying that specific neuronal subpopulations are primed to become circuit-reorganizing neurons after injury. Moreover, we show that the ability of neuronal subpopulations to contribute to recovery entails an inherent trade-off between the activation of cellular stress responses versus that of transcriptional programs involved in circuit reorganization. We also demonstrate that the profound impairment of neurological recovery after SCI in old mice reflects a failure to coordinate the formation of a tripartite neuroprotective barrier between immune-privileged neural tissue and extra-neural lesion environments. This failure leads to overwhelming immune infiltration and catastrophic neuronal death into neural tissue, precluding neurological recovery. We built on this discovery to develop a new gene therapy strategy that reactivated border-forming cells, prevented lesion expansion, and improved neurological recovery in old mice. This mechanism-based therapeutic strategy opens new avenues to accelerate wound repair and increase neurological recovery in humans with SCI.

The *Tabulae Paralytica* embody a number of technological and conceptual advances that demonstrate how genome-wide single-cell and spatial technologies can deliver new insights into uninjured and perturbed tissues. On a conceptual level, we demonstrate how the increasing scale of single-cell technologies enables a comprehensive interrogation of the experimental manipulations relevant to any given disease within a single study. On a technical level, we establish the possibility of extending spatial transcriptomics into three and even four spatial and temporal dimensions within a common coordinate framework. Moreover, we show that the integration of multi-omic single-cell atlases allows us to overlay patterns of chromatin accessibility onto a four-dimensional spatial model. Finally, our findings illustrate the power of cell type and spatial prioritization, as implemented by Augur and Magellan, to resolve the molecular basis of diseases or biological perturbations using single-cell and spatial genomics.

The *Tabulae Paralytica*, or ‘atlases of spinal cord injury,’ will serve as (i) a foundational resource to understand the pathobiology of SCI; (ii) a reference of cellular and molecular responses to predict and interrogate the consequences of new therapeutic strategies; (iii) a conceptual and technical framework to advance spatially resolved single-cell studies of disease and biological perturbations; and (iv) a translational resource to uncover new biological mechanisms of SCI that can be exploited to develop therapies to repair the injured human spinal cord.

## Methods

### Mouse model and experimental conditions

Adult male or female C57BL/6 mice (15-25 g body weight, 8-15 weeks of age) or transgenic mice were used for all experiments. Aged mice were purchased from JAX at 60 weeks of age (stock no. 000664). Vsx2^Cre^ (MMMRRC 36672, also called Chx10^Cre^) transgenic mouse strain was bred and maintained on a mixed genetic background (C57BL/6). Housing, surgery, behavioral experiments and euthanasia were all performed in compliance with the Swiss Veterinary Law guidelines. Manual bladder voiding and all other animal care was performed twice daily throughout the entire experiment. All procedures and surgeries were approved by the Veterinary Office of the Canton of Geneva (Switzerland; authorizations GE/145/2). Spinal cord crushes were performed as previously described^15, 63^. For time-course experiments, animals were euthanised at 1 day, 4 days, 7 days, 14 days, 1 month, or 2 months post-injury. Crush injuries were performed at multiple severities by including spacers within No. 5 Dumont forceps (Fine Science Tools) such that when closed, there was a maximal distance of 1 mm, 0.5 mm, 0.25 mm, or 0 mm (no spacer) with a tip width of 0.5 mm. Dorsal hemisection SCIs were performed as previously described^12^. For dorsal hemisection SCI, a laminectomy was made at the mid-thoracic level (T10) and the dorsal half of the spinal cord was cut using a microscapel. Contusion SCIs were performed as previously described^9, 64^.

Minocycline was administered with intraperitoneal injections as previously described^73–75^, with a loading dose of 50 mg/kg at 1 h post-injury and 24 h post-injury, followed by maintenance doses of 25 mg/kg every 24 h for the next five days. Methylprednisolone was administered intramuscularly as previously described^172^, with a loading dose of 60 mg/kg at 1 h post-injury then an additional 30 mg/kg dose every 6 h for 24 h. Chondroitinase ABC was delivered via lentiviral injections as previously described^173^. Briefly, the *Proteus vulgaris* ChABC gene was previously modified to make a mammalian-compatible engineered ChABC gene^138^. The modified ChABC cDNA was subcloned into a lentiviral transfer vector (termed LV-ChABC) with the mouse phosphoglycerate kinase promoter^138^. The final viral titer was 479 *µ*g/mL of P24, corresponding to ∼10^6^ TU/*µ*L. A control lentiviral vector (termed LV-GFP) was generated from the same transfer vector containing the cDNA coding for GFP, with a viral titer of 346 *µ*g/mL of P24.

### Viral vectors and vector production

Viruses used in this study were either acquired commercially or produced at the EPFL core facility. The following AAV plasmids were used and detailed sequence information is available as detailed or upon request: AAV-CAG-flex-human Diphtheria Toxin Receptor (DTR, plasmid gifted by Prof. S. Arber), and produced as AAV5 at the EPFL Bertarelli Foundation Platform in Gene Therapy and SIN-cPPT-PGK-FGF2-WPRE,SIN-cPPT-PGK-EGF-WPRE SIN-cPPT-PGK-VEGF-WPRE, SIN-cPPT-GFAP-GDNF-WPRE, and LV-PGK-ChABC (gifted by Prof. E. Bradbury). Injection volumes, coordinates and experimental designs are described below.

### Biological repair intervention in aging mice

General surgical procedures have been previously described in detail^15, 63, 64^. Surgeries were performed at EPFL under aseptic conditions and under 1-2% isoflurane in 0.5-1 L/min flow of oxygen as general anesthesia, using an operating microscope (Zeiss) and rodent stereotaxic apparatus (David Kopf) as previously described^63, 64^. LV injections were made two days before SCI to allow time for expression, and were targeted over the intended spinal cord segment to be injured. LVs were injected into four sites (two sets of bilateral injections, 0.30 *µ*L/injection [all vectors diluted to 600 *µ*g P24/mL in sterile saline]) 0.6 mm below the surface at 0.15 *µ*L per minute using glass micropipettes connected via high-pressure tubing (Kopf) to 10 *µ*L syringes under the control of a microinfusion pump. Moderate crush SCIs were introduced at the level of T10/T11 after laminectomy of a single vertebra by using No. 5 Dumont forceps (Fine Science Tools) with a spacer so that when closed a 0.5 mm space remained, and with a tip width of 0.5 mm to completely compress the entire spinal cord laterally from both sides for 5 s. After surgeries, mice were allowed to wake up in an incubator. Analgesia, buprenorphine (Essex Chemie AG, Switzerland, 0.01-0.05 mg/kg s.c.) or carprofen (5 mg/kg s.c.), was given twice daily for 2-3 days after surgery. Animals were randomly assigned numbers and thereafter were evaluated blind to experimental conditions. Fourteen days after SCI, all mice were evaluated in an open field and all animals exhibiting any hindlimb movements were not studied further.

### Neuron subpopulation-specific ablation

For ablation experiments with diphtheria toxin, Vsx2^Cre^ mice were subjected to crush SCI as described above. Three sets of bilateral injections of AAV5-CAG-FLEX-DTR^174^ were made over the T9, T10, and T11 spinal segments (0.25 *µ*L per injection) at a depth of 0.6 mm below the dorsal surface and separated by 1 mm. Two weeks after spinal infusions, mice received intraperitoneal injections of diphtheria toxin (Sigma, D0564) diluted in saline (100 *µ*g/kg) to ablate Vsx2 neurons. Kinematics were evaluated in all mice before ablation, one week, and two-weeks post-ablation.

### Behavioural assessments

Behavioral procedures have been previously described in detail^64, 174, 175^. Briefly, during overground walking, bilateral leg kinematics were captured with twelve infrared cameras of a Vicon Motion Systems (Oxford, UK) that tracked reflective markers attached to the crest, hip, knee, ankle joints, and distal toes. The limbs were modelled as an interconnected chain of segments and a total of 80 gait parameters were calculated from the recordings. To evaluate differences between experimental conditions, as well as to identify the most relevant parameters to account for these differences, we implemented a multistep multifactorial analysis based on principal component analysis, as previously described in detail^9, 64, 174^, and coupled to automated, markless tracking software^176^.

### Perfusions

Mice were perfused at the end of the experiments. Mice were deeply anesthetized by an intraperitoneal injection of 0.2 mL sodium pentobarbital (50 mg/mL). Mice were transcardially perfused with PBS followed by 4% paraformaldehyde in PBS. Tissue was removed and post-fixed overnight in 4% paraformaldehyde before being transferred to PBS or cryoprotected in 30% sucrose in PBS.

### Immunohistochemistry

Immunohistochemistry was performed as previously described^15, 63, 64^. Perfused post-mortem tissue was cryoprotected in 30% sucrose in PBS for 48 h before being embedded in cryomatrix (Tissue Tek O.C.T, Sakura Finetek Europe B.V.) and freezing. 30 *µ*m thick transverse or horizontal sections of the spinal cord were cut on a cryostat (Leica), immediately mounted on glass slides and dried or in free floating wells containing PBS plus 0.03% sodium azide. Primary antibodies were: rabbit anti-GFAP (1:1000; Dako); guinea pig anti-NeuN (1:300; Millipore); chicken anti-RFP (1:500, Novus Biologicals); rabbit anti-Chx10 (also known as Vsx2) (1:500, Novus Biologicals); rat anti-CD45 (1:100, BD Biosciences); goat anti-Sox9 (1:200, Novus Biologicals); rabbit anti-Id3 (1:500; Cell Signalling Technology); rabbit anti-PKD1L2 (1:1000; Merck Millipore); mouse anti-GFAP (1:1000; MARK). Fluorescent secondary antibodies were conjugated to Alexa 488 (green), or Alexa 405 (blue), or Alexa 555 (red), or Alexa 647 (far red) (ThermoFisher Scientific, USA). The nuclear stain was 4’,6’-diamidino-2-phenylindole dihydrochloride (DAPI; 2 ng/mL; Molecular Probes). Sections were imaged digitally using a slide scanner (Olympus VS-120 Slide scanner) or confocal microscope (Zeiss LSM880 + Airy fast module with ZEN 2 Black software). Images were digitally processed using ImageJ (ImageJ NIH) software or Imaris (Bitplane, version 9.0.0).

### Tissue clearing (CLARITY)

Samples were incubated in X-CLARITY hydrogel solution (Logos Biosystems Inc., South Korea) for 24 h at 4*^◦^*C with gentle shaking. Samples were then degassed and polymerized using the X-CLARITY Polymerisation System (Logos Biosystems Inc., South Korea), followed by washes in 0.001 M PBS for 5 minutes at room temperature. Samples were next placed in the X-CLARITY Tissue Clearing System (Logos Biosystems Inc., South Korea), set to 1.5 A, 100 RPM, 37*^◦^*C, for 29 h. Clearing solution was made in-house with 4% sodium dodecyl sulfate (SDS), 200 mM boric acid with dH_2_O, pH adjusted to 8.5. Following this, samples were washed for at least 24 h at room temperature with gentle shaking in 0.1 M PBS solution containing 0.1% Triton X-100 to remove excess SDS. Finally, samples were incubated in 40 g of Histodenz dissolved in 30 mL of 0.02 M PB, pH 7.5, 0.01% sodium azide (refractive index 1.465) for at least 24 h at room temperature with gentle shaking prior to imaging.

### 3D imaging

Imaging of cleared tissue was performed using a customized mesoSPIM^177^ and CLARITY-optimized light-sheet microscope (COLM)^178^. A custom-built sample holder was used to secure the central nervous system in a chamber filled with RIMS. Samples were imaged using either a 1.25*×* or 2.5*×* objective at the mesoSPIM and a 4*×* or 10*×* objective at the COLM with one or two light sheets illuminating the sample from both the left and right sides. The voxel resolution in the x-, y- and z directions was 5.3 *µ*m *×* 5.3 *µ*m *×* 5 *µ*m for the 1.25*×* acquisition and 2.6 *µ*m *×* 2.6 *µ*m *×* 3 *µ*m for the 2.5*×* acquisition. The voxel resolution of the COLM was 1.4 *µ*m *×* 1.4 *µ*m by 5 *µ*m. Images were generated as 16-bit TIFF files and then stitched using Arivis Vision4D (Arivis AG, Munich, Germany). 3D reconstructions and optical sections of raw images were generated using Imaris (Bitplane, version 9.0.0) software.

### Histological analysis

To quantify immune invasion after different models of SCI, we measured the percentage of CD45 immunopositive area after binarizing the fluorescent signal using the image analysis software Fiji. To assess the formation of astrocyte scar borders after SCI in young and old mice, we counted the number of Sox9^ON^ cells using the image analysis software QuPath and the cell detection functionality with default settings (version 0.4.3). We then classified Sox9^ON^ cells as either Id3^ON^ or Id3^OFF^ by setting a mean signal intensity threshold. We used the same approach to count the number of NeuN^ON^ and PKD1L2^ON^ neurons, to assess the resilience of CSF-contacting neurons after SCI. To quantify the number of *Vsx2*^ON^ neurons after neuronal subpopulation-specific ablation with DTR we used the image analysis software Imaris (Bitplane, version 9.0.0).

### Chronophotography

Chronophotography was used to generate a representative series of still pictures arranged in a single photograph to illustrate the locomotor abilities of mice. Videos at 25 fps or photographs at 15 fps were recorded while mice were performing locomotor tasks such as quadrupedal walking on the runway. Images from these recordings were chosen to best illustrate the different consecutive phases of walking of the hindlimbs, i.e. stance phases and swing phases. The frequency of chosen pictures varied due to the varying velocity of the mice. The series of pictures were assembled in Photoshop while blending out non-essential details.

### snRNA sequencing library preparation

Single-nucleus dissociation of the mouse lumbar spinal cord was performed according to our established procedures^9, 60^. Following euthanasia by isoflurane inhalation and cervical dislocation, the lumbar spinal cord site was immediately dissected and frozen on dry ice. We denounced spinal cords in 250 *µ*L sucrose buffer (0.32 M sucrose, 10 mM HEPES [pH 8.0], 5 mM CaCl_2_, 3 mM Mg acetate, 0.1 mM EDTA, 1 mM DTT) and 0.1% Triton X-100 with the Kontes Dounce Tissue Grinder. 1.1 mL of sucrose buffer was then added and filtered through a 40 *µ*m cell strainer. The lysate was centrifuged at 3200 g for 5 min at 4*^◦^*C. The supernatant was decanted, and 1 mL of sucrose buffer added to the pellet and incubated for 1 min. The pellet was homogenized using an Ultra-Turrax and 3 mL of density buffer (1 M sucrose, 10 mM HEPES [pH 8.0], 3 mM Mg acetate, 1 mM DTT) was added below the nuclei layer. The tube was centrifuged at 3200 g at 4*^◦^*C for 10 min and supernatant was immediately poured off. Nuclei on the bottom half of the tube wall were resuspended in 100 *µ*L PBS with 1% BSA for subsequent single-nucleus RNA sequencing or in 10x Nuclei Buffer (catalog no. 2000153, 10x Genomics) for subsequent single-nucleus multiome sequencing. Resuspended nuclei were filtered through a 30 *µ*m strainer, and adjusted to 1,000 nuclei/*µ*L. We carried out snRNA-seq library preparation using the 10x Genomics Chromium Single Cell Gene Expression Kit Version 3.1. The nuclei suspension was added to the Chromium RT mix to achieve loading numbers of 10,000 nuclei. For downstream cDNA synthesis, library preparation and sequencing, the manufacturer’s instructions were followed.

### Multiome sequencing library preparation

We carried out snRNA- and ATAC library preparation using the 10x Genomics Chromium Single Cell Multiome ATAC + Gene Expression Kit. First, the transposition mix was added to the resuspended nuclei followed by a 60 min incubation at 37*^◦^*C. The transposed nuclei were added to the Chromium RT mix to achieve loading numbers of 10,000 nuclei. The manufacturer’s instructions were followed for downstream cDNA synthesis, library construction, indexing and sequencing.

### Spatial transcriptomics library preparation

We carried out two separate experiments to study the cytoarchitecture of the lesion microenvironment after SCI. First, we prepared sections from uninjured mice, 7 days and 2 months after crush SCI (performed with a 0.5 mm spacer as described above). For each experimental condition, we prepared sections from the lesion epicenter of three independent biological replicates. Second, to prepare our four-dimensional spatiotemporal atlas, we collected sections throughout the entire spinal cord of mice from each of the three experimental conditions. The spinal cord injury sites of mice were embedded in OCT and cryosections were generated at 10 *µ*m at –20*^◦^*C. For the four-dimensional atlas, every fifth section was collected throughout the entire dorsoventral axis of each spinal cord. Sections were immediately placed on chilled Visium Tissue Optimization Slides (catalog no. 1000193, 10x Genomics) or Visium Spatial Gene Expression Slides (catalog no. 1000184, 10x Genomics). Tissue sections were then fixed in chilled methanol and stained according to the Visium Spatial Gene Expression User Guide (catalog no. CG000239 Rev A, 10x Genomics) or Visium Spatial Tissue Optimization User Guide (catalog no. CG000238 Rev A, 10x Genomics). For gene expression samples, tissue was permeabilized for 12 min, which was selected as the optimal time based on tissue optimization time-course experiments. Brightfield histology images were taken using a 10*×* objective on a slide scanner (Olympus VS-120 Slide scanner). For tissue optimization experiments, fluorescent images were taken with a TRITC filter using a 10*×* objective and 400 ms exposure time. Libraries were prepared according to the Visium Spatial Gene Expression User Guide.

### Read alignment

Following sequencing on our HiSeq4000 (EPFL Gene Expression Core Facility), snRNA-seq reads were aligned to the latest Ensembl release of the mouse genome (GRCm38.101), and a matrix of unique molecular identifier (UMI) counts was obtained using CellRanger (10x Genomics, version 4.0.0)^30^. For spatial transcriptomics, a spatial expression UMI count matrix was obtained using SpaceRanger (10x Genomics, version 1.0.0). For the multiome dataset, RNA-seq and ATAC-seq data were aligned to the reference genome using CellRanger-ARC (10x Genomics, version 2.0.0), and a matrix of UMI counts was obtained for the RNA modality. The ATAC modality was then processed further using ArchR, as described below.

### snRNA-seq preprocessing and quality control

Droplet-based snRNA-seq experiments are known to be affected by ambient RNA contamination, whereby freely floating RNA molecules are encapsulated along with a cell or nucleus in a single droplet and spuriously attributed to the endogenous expression profile of the encapsulated cell^179^. The presence of ambient RNA is a potential source of batch effects and spurious differential expression. To mitigate this possibility, we used CellBender^180^ to remove ambient RNA molecules and filter empty droplets. CellBender remove-background was run for 50 epochs with a learning rate of 5e-5. Corrected count matrices were then imported into Seurat^181^ for further quality control. Quality control metrics included the number of UMIs per cell, the number of genes detectably expressed per cell, and the proportion of UMI counts arising from mitochondrial genes. For the pilot dataset, cells with between 200 and 40,000 UMIs, and less than 7,500 genes expressed, were retained. For the snRNA-seq and multiome datasets, cells with at least 200 UMIs were retained. Additional quality control was performed for the multiome dataset on the basis of the ATAC modality, as described further below. Low-quality libraries were identified as those with distributions of number of UMIs, number of genes expressed, or proportion of mitochondrial counts that differed markedly from the remainder of the libraries in the dataset, and a total of three low-quality libraries (two from the snRNA-seq dataset and one from the multiome dataset) were removed.

Putative doublets then were identified and filtered using a combination of approaches. We tested the performance of four computational methods for doublet detection in our pilot dataset, including DoubletFinder^182^, scDblFinder^183^, scds^184^, and Scrublet^185^. On the basis of this analysis, we selected scDblFinder and scds as the two methods that (i) did not display an overt bias towards doublet detection for cells of any particular type, (ii) which showed the highest agreement with one another, and (iii) which were also found to be among the top-performing methods in an independent benchmark^186^. We adopted a conservative approach by filtering barcodes from the union of those called doublets by either scDblFinder or scds in both the pilot and snRNA-seq datasets. For the multiome dataset, doublets were instead identified and filtered using ArchR, as described below.

### Integration and cell type annotation

Prior to clustering and cell type annotation, we first performed batch effect correction and data integration across experimental conditions using Harmony^158^. Gene expression counts were normalized to counts per 10,000 and log-transformed, and the top 2,000 variable genes were identified using the ‘vst’ method in Seurat. Gene expression values were then scaled and centered and provided as input to Harmony, which was run with 50 principal components. The integrated Harmony embeddings were then provided as input to *k*-nearest neighbor graph construction and Leiden clustering using the default Seurat workflow^181^, as in our previous studies^9,60, 87^. Cell types were then manually annotated on the basis of marker gene expression, guided by previous studies of the mouse spinal cord^47–57, 59–61, 87^ and other relevant cell atlases of major cell types^93, 102^. Local and projecting neuronal subpopulations were annotated on the basis of *Nfib* and *Zfhx3* expression, respectively^59^. Clusters corresponding to damaged cells or doublets that had survived initial quality control were removed at this stage. In the pilot dataset, we performed an initial round of clustering to identify coarse cell types with a resolution of 0.05, followed by subclustering of neurons (resolution = 0.5) and glia (resolution = 0.1) to annotate more fine-grained subtypes. In the snRNA-seq dataset, we repeated the clustering analysis for multiple values of the resolution parameter (0.01, 0.05, 0.2, 0.5, 2) in order to annotate cell types across multiple resolutions (e.g. neurons *→* ventral neurons *→* ventral excitatory interneurons). We then used the clustree package^85^ to link clusters across adjacent resolutions into a hierarchical clustering tree, as previously described^7,9,60^.

### Pilot dataset and meta-analysis of published spinal cord snRNA-seq datasets

We conducted an initial pilot experiment to confirm that our dissociation procedures enabled the recovery of all the cell types comprising the mouse spinal cord. snRNA-seq libraries were prepared from one uninjured mouse and one mouse 7 days after crush SCI, and deeply sequenced to a target depth of 100,000 reads per nucleus. After preprocessing and quality control as described above, we retained 9,170 nuclei from the injured sample and 9,099 nuclei from the uninjured sample. Following data integration and cell type annotation as described above, we confirmed that we recovered the major cell types of the spinal cord in both the injured and uninjured spinal cords, and that changes in cell type proportions across experimental conditions were concordant with the established pathophysiology of SCI. We then compared the cell type proportions in our pilot dataset to those in 16 published single-cell datasets from the mouse spinal cord^47–62^. Automated cell type annotation of published datasets was performed using the label transfer workflow in Seurat, with our own previously published dataset from the uninjured lumbar spinal cord^60^ used as the reference. We confirmed that the label transfer workflow yielded reliable predictions by comparing automated cell type annotations to manual annotations from a published dataset^50^. This analysis established that our dissociation protocols allowed us to recover the major cell types of the spinal cord in proportions consistent with published single-nucleus RNA-sequencing studies of the whole adult spinal cord. Finally, we took advantage of our deeply sequenced pilot dataset to calibrate our target sequencing depth of our main experiments, and selected a target depth of 75,000 reads per nucleus on the basis of downsampling analysis of the pilot dataset.

### snRNA-seq atlas

For the snRNA-seq dataset, preprocessing, quality control, data integration, and cell type annotation were performed as described above, yielding a dataset comprising 435,099 nuclei from 54 mice spanning 18 experimental conditions. Marker genes were identified for each cluster using the FindMarkers function in Seurat. We visualized the distribution of cell types, experimental conditions, and the expression of marker genes with UMAP embeddings of both the entire dataset as well as each major cell type. Marker gene dotplots were constructed using the DotPlot function in Seurat. Cell type proportions were visualized using sunburst plots^187^ and Sankey diagrams. The cell cycle positions of astroependymal cells were estimated using tricycle^188^. The expression of a previously described gene module^101^ associated with BBB dysfunction was estimated using the Seurat function AddModuleScore. Unless otherwise stated, all cell subtype analyses were performed at level 5 of the clustering tree (corresponding to a resolution of 2).

### Cell type proportions

Testing for differences in cell type proportions within single-cell data can lead to false discoveries because the data is compositional in nature, and consequently, increases the proportion of one cell type can cause an artefactual decrease in the proportions of every other cell type^189^. To avoid this pitfall, we used the propeller method^190^, as implemented in the speckle R package, to test for differences in cell type proportions between experimental conditions, as an independent benchmark showed this to be among the most accurate methods in balancing control of the false discovery rate with statistical power^191^. The details of individual cell type proportion analyses are described below.

### Differential expression

To identify genes differentially expressed between experimental conditions, we performed differential expression (DE) analysis by aggregating expression from all cells of a given type within each replicate into a ‘pseudobulk’ profile, as previously described^87^ and implemented in the Libra R package, (https://github.com/neurorestore/Libra). In our previous work^87^, we demonstrated that this approach allowed us to overcome false discoveries caused by variability between biological replicates^192^. We showed that widely-used single-cell DE methods can conflate this variability with the effect of a biological perturbation, leading to hundreds or even thousands of false discoveries. We therefore instead used the likelihood ratio test implemented in edgeR^193^ to identify DE genes between pseudobulks from each cell type. The details of individual DE analyses are described below.

### GO enrichment analysis

GO term annotations for mouse were obtained from the Gene Ontology Consortium website. GO terms annotated to less than five genes were excluded. The average expression level of genes associated with each GO term in individual cells was calculated using the Seurat function AddModuleScores, which controls for the average expression of randomly selected control features. Linear mixed models were then used to test for differences in GO module scores test across experimental conditions, using the ‘lmerTest’ R package to optimize the restricted maximum likelihood and obtain p-values from the Satterthwaite approximation for degrees of freedom. The details of individual GO enrichment analyses are described below.

### Cell type prioritization

To identify cell types activated in response to each biological perturbation captured in the *Tabulae Paralytica*, we employed a machine-learning method for cell type prioritization that we previously developed, named Augur^9,60, 127^. Briefly, Augur seeks to rank cell types based on the intensity of their transcriptional response to a biological perturbation. The key assumption underlying Augur is that cell types undergoing a profound response to a perturbation should become more separable, within the highly multidimensional space of gene expression, than less affected cell types. To quantify this separability, we framed this problem as a classification task. Augur first withholds a proportion of experimental condition labels, then trains a random forest classifier to predict the condition from which each cell was obtained (for instance, SCI or uninjured). The accuracy with which this prediction can be made from single-cell gene expression measurements is then evaluated in cross-validation, and quantified using the area under the receiver operating characteristic curve (AUC). This process is repeated separately for each cell type. The AUC then provides a quantitative measure of separability that can be used to rank cell types based on the relative magnitude of their response to an arbitrary perturbation. We refer to this process as cell type prioritization. Augur was run with default parameters directly on the UMI count matrix for all comparisons. To evaluate the robustness of cell type prioritizations to the resolution at which neuronal subtypes were defined in the snRNA-seq data, we applied Augur at various clustering resolutions, and visualized the resulting cell type prioritizations both on a hierarchical clustering tree^85^ of cell types and as a progression of UMAPs^9^. The details of individual cell type prioritization analyses are described below.

### Conserved and divergent neuronal responses to SCI

To identify spinal cord neurons that were resilient or susceptible to SCI, we computed the log_2_-odds ratio between the uninjured spinal cord and each experimental condition in which the injured spinal cord was profiled at 7 days post-injury, using neuron subtypes defined at level 4 of the clustering tree (corresponding to a resolution of 0.5), then identified resilient or susceptible neuron subtypes using a t-test on the log_2_-odds ratios. To identify DE genes specific to CSF-contacting neurons at the most acute phase of the injury response, we used edgeR to test for an interaction term between neuronal subtype and experimental condition at 1 day post-injury, using pseudobulk gene expression profiles.

To quantify the degree to which transcriptional responses to injury were conserved across neuron subtypes, we computed the Spearman correlation between log-fold changes estimated by edgeR between each pair of level 4 neuron subtypes. For genes that were not quantified in one of the two subtypes, missing log-fold change values were replaced with zeros.

To characterize the conserved early response of neurons to SCI, we first filtered to genes that were differentially expressed within individual level 4 neuron subtypes at a 10% false discovery rate. We then sorted these genes first by the number of neuron subtypes in which they were DE, and second by the mean absolute log-fold change estimated by edgeR.

To quantify the expression of transcriptional programs associated with projection growth and morphogenesis, we used the average expression of genes associated with the GO term “GO:0031175” (neural projection development) to construct a circuit reorganization score, as described above. We then computed the basal expression of this circuit reorganization score as the median GO module score in the uninjured spinal cord for each level 4 neuron subtype. To quantify upregulation of the circuit reorganization score after injury, we subtracted the basal expression score from the GO module score at each timepoint post-injury to yield induced expression scores. We then calculated the Pearson correlation between basal and induced circuit reorganization scores. We carried out similar analyses for the GO terms “GO:0061564” (axon development) and “GO:0016358” (dendrite development). Cell type prioritization was performed by comparing neurons from each level 4 subtype at each timepoint post-injury to neurons from the uninjured spinal cord.

### Neurons remain differentiated after CNS injury

Individual marker genes for each neuron subtype were manually curated from literature after cross-referencing with other atlases, as described above. DE analysis was performed by comparing neurons from each level 4 subtype at each time-point post-injury to neurons from the uninjured spinal cord using edgeR as described above, with a 5% false discovery rate. We also constructed unbiased lists of the top-*n* marker genes for each level 4 neuron subtype (for *n* = 5, 10, or 50) using the FindMarkers function in Seurat. We used the AddModuleScore function to summarize the average expression of the top-*n* marker genes in each individual neuron, then used a linear mixed model to test for differences across experimental conditions as described above for GO enrichment analyses.

### Facilitating and inhibiting molecule expression in the injured spinal cord

We visualized the expression of key facilitating and inhibiting molecules across the cell types and subtypes of the spinal cord using clustering trees, with the scaled mean expression for each cell type or subtype calculated as in the Seurat function DotPlot. To identify genes coordinately up- or downregulated across level 4 neuron subtypes in response to ChABC treatment, we used edgeR to perform DE analysis as described above, and performed a one-sample t-test on log-fold change estimates from edgeR^9^. We then used linear mixed models to perform GO enrichment analysis of ChABC treatment for each level 4 neuron subtype, as described above, and performed a one-sample t-test on coefficients estimated by the mixed models.

### Cellular divergence between animal models of SCI

Cell type proportions were compared using propeller, as described above, both for coarse cell types and for the most fine-grained subtypes of immune cells. Cell type prioritization was performed by comparing neurons from each level 4 sub-type between each pair of animal models (crush, contusion, or hemisection). Separately, we tested for differences in the AUCs of dorsal and ventral level 4 neuron subtypes by comparing neurons from mild, moderate, severe, or complete injuries to neurons from the uninjured spinal cord.

### Immunomodulation does not confer neuroprotection after spinal cord injury

Cell type prioritization was performed by comparing cell types at each resolution of the clustering tree from drug-treated and untreated but injured spinal cords. The proportions of coarse cell types were compared using propeller as described above.

To dissect more subtle transcriptional effects of neuroprotective agents on surviving neurons, we developed a machine-learning approach to identify neurons displaying an uninjured transcriptional phenotype. For each experimental condition involving injured and untreated mice (i.e., excluding the uninjured and drug-treated conditions), we trained a random forest model on scaled and log-normalized gene expression data to distinguish cells from that condition (“injured” cells) to cells from the uninjured spinal cord (“uninjured” cells). We then applied each of these models in turn to neurons from the methylprednisolone and minocycline conditions, in order to predict whether they displayed an injured or uninjured phenotype. The modal prediction across all models was then assigned to each neuron. To further characterize the transcriptional programs induced by neuroprotective agents, we then used linear mixed models to perform GO enrichment analysis of methylprednisolone or minocycline treatment for each level 4 neuron sub-type, as described above, and performed a one-sample t-test on coefficients estimated by the mixed models.

### Sexually dimorphic responses to SCI are subtle

Cell type prioritization was performed by comparing cell types at each resolution of the clustering tree from male and female spinal cords. The range of AUC values assigned by Augur in cross-validation was then compared to that observed in other comparisons involving the injured spinal cord at 7 days post-injury. Cell type proportions were compared using propeller, as described above, both for coarse cell types and for the most fine-grained subtypes of immune cells.

### Catastrophic failure of tripartite barrier formation in old mice

Cell type prioritization was performed by comparing cell types at each resolution of the clustering tree from young and old mice, and the range of AUC values assigned by Augur in cross-validation was again compared to that observed in other comparisons involving the injured spinal cord at 7 days post-injury. The proportion of *Id3*-expressing astrocytes was compared between young and old mice using a *χ*^2^ test. Gene modules associated with blood-brain barrier endothelial cell identity and peripheral endothelial cell identity were obtained from the literature^101^, and their expression in individual vascular cells was calculated using the Seurat function AddModuleScore.

DE analysis was performed as described above by comparing cells from young and old mice after SCI, for cell subtypes at level 4 of the clustering tree (resolution = 0.5) and with a false discovery rate of 5%. To quantify the heterogeneity of gene expression across cell types, we calculated two summary statistics. First, we defined the direction consistency as the proportion of cell types in which the sign of the log-fold change was the same as the modal sign. For example, if a gene was upregulated in eight of ten cell types and downregulated in the other two, the direction consistency would be 80%. Second, we defined the response heterogeneity as the standard deviation of the log-fold change across cell types.

### snATAC-seq preprocessing and quality control

Preprocessing and quality control of the ATAC modality within our multiome dataset was carried out using CellRanger-ARC and ArchR^157^. Reads were mapped to the reference genome with CellRanger-ARC, and arrow files were created from the resulting fragment files. Nuclei were first filtered based on the RNA modality as described above, and subsequently additional quality control was performed in ArchR. We initially ran ArchR with very lenient filtering in order to determine optimal quality control parameters (minimum TSS enrichment score = 0, minimum fragments per cell = 100), and selected optimal parameters based on the joint distribution of these parameters. Arrow files were subsequently regenerated after filtering nuclei to those with a minimum TSS enrichment score of 4 and a minimum of 4,000 fragments per cell. Doublet detection and filtering was performed using the ArchR functions addDoubletScores and filterDoublets, both with default parameters. These steps afforded matrices of 40,526 nuclei that passed quality control in both the RNA and ATAC modalities.

To link cell types in the multiome dataset to the cellular taxonomy derived from our snRNA-seq atlas, we devised a hierarchical label transfer strategy using Symphony^158^. Briefly, we first used Symphony to perform automated cell type assignment in the multiome dataset at the highest level of the clustering tree (level 1, resolution = 0.01). We then used Symphony to perform automated cell type assignment at the second level of the clustering tree (resolution = 0.05), considering only subtypes of the assigned coarse cell types as potential matches for each nucleus. This process was repeated iteratively for each level of the clustering tree. We validated the accuracy of this strategy using a leave-library-out cross-validation approach within the snRNA-seq atlas, in which entire libraries were withheld from the atlas and automated cell type assignment was compared to the manual cell type assignment derived from the entire dataset. We found that the hierarchical approach improved the accuracy of automated cell type assignment relative to a non-hierarchical version of the same approach, in which all cell subtypes at any given level were considered as potential matches, particularly at more granular levels of the clustering tree. For cell type assignment in the multiome dataset, we ran Symphony using the hierarchical approach with 100 soft cluster centroids, 100 principal components, and 20 nearest-neighbors, then made additional manual adjustments to cell type annotations for a handful of cell subtypes that showed discordant marker gene expression.

Peak calling in the snATAC-seq dataset was then carried out using the default ArchR workflow, including peak calling with MACS2^194^ on pseudobulk replicates from each cell type, followed by peak merging across cell types using an iterative overlap removal procedure. We repeated this process for cell type definitions at each level of the clustering tree and found that peak calling at more granular resolutions allowed us to preferentially detect distal regulatory elements. Unless otherwise noted, downstream analyses were carried out on the peak matrix called with coarse cell type definitions (level 1, resolution = 0.01).

### Transcription factor activities

Transcription factor deviations were estimated by chromVAR^159^, using motif sets from the chromVAR package (ENCODE, HOMER, and CisBP) as well as the 2020 version of JASPAR^195^. Transcription factor motifs associated with cell type identity were identified using a Wilcoxon rank-sum test, as in the Seurat function FindMarkers. Linear mixed models were used to identify transcription factor motifs differentially active in cells from injured spinal cords, using the ‘lmerTest’ R package to optimize the restricted maximum likelihood and obtain p-values from the Satterthwaite approximation for degrees of freedom, and a false discovery rate of 10%.

To identify transcription factors that were up- or downregulated across all level 4 neuron subtypes at 7 days post-injury, we performed a one-sample t-test on coefficients estimated by the mixed models. To identify transcription factors with discordant patterns of up- or downregulation at 2 months post-injury, we devised a permutation-based statistical approach. Neuron subtype assignments at level 4 of the clustering tree were randomized within each experimental condition, and differential activity testing was performed using linear mixed models in the permuted data. This process was repeated 100 times, and the standard deviation of model coefficients was calculated for the observed and permuted datasets. The resulting z statistics were then converted to p-values using a standard normal distribution and significantly divergent motifs were identified using a 10% false discovery rate.

To identify transcription factors associated with dysfunction of the tripartite barrier after SCI, BBB dysfunction module scores^101^ were first estimated from the RNA modality of the multiome data, as in the snRNA-seq atlas. chromVAR deviations in the ATAC modality were then correlated to the resulting module scores, using the Pearson correlation and restricting this analysis to vascular cells. Linear mixed models were then used to identify motifs that were differentially accessible at 7 days in level 4 subtypes associated with the tripartite barrier, including vascular leptomeningeal cells, capillary endothelial cells, pericytes, arachnoid barrier cells, reactive astrocytes, and OPCs. Analyses of differentially active transcription factors in neurons or blood-brain barrier cell types were carried out using chromVAR deviation matrices derived from peak matrices at the relevant resolution of the clustering tree, as described above.

### Differential accessibility

To identify differentially accessible peaks, we extended the workflow for pseudobulk DE analysis in Libra to peak count matrices derived by ArchR. Cells of each type were aggregated within replicates to form pseudobulks, and then testing for differential accessibility was performed using the likelihood ratio test implemented in DESeq2^196^. The evolutionary conservation of each peak was quantified as the mean phyloP conservation score from the 60-way vertebrate data set^165^ of all bases within the peak. The evolutionary conservation of all peaks open within a given cell was further summarized by taking the mean phyloP score across all accessible peaks in that cell.

### Evolutionary divergence in circuit reorganization

The expression of cellular stress response programs in the snRNA-seq atlas was estimated by using the Seurat function AddModuleScore to summarize the mean expression of genes associated with the GO term GO:0033554 (“cellular response to stress”). The resulting score was then correlated with the circuit reorganization score described above across all neurons.

### Spatial transcriptomics preprocessing and quality control

Following read alignment and count matrix generation with SpaceRanger as described above, Seurat^181^ was used to calculate quality control metrics for each spatial barcode, including the number of genes detected, number of UMIs, and proportion of reads aligned to mitochondrial genes. Low-quality barcodes were filtered by removing those with less than 3,000 or more than 45,000 UMIs. Low-quality sections were identified as those with distributions of number of UMIs, number of genes expressed, or proportion of mitochondrial counts that differed markedly from the remainder of the sections in the dataset, and removed. In the two-dimensional spatial dataset, these steps afforded a UMI count matrix comprising 33,941 spatial barcodes from nine biological replicates (three from each experimental condition). In the three-dimensional spatial dataset, these steps afforded a UMI count matrix comprising 37,558 spatial barcodes from three biological replicates (one from each experimental condition).

### Registration to a common coordinate framework

We aligned all spatial transcriptomics sections into a common coordinate system using a custom image analysis pipeline that includes preprocessing, registration and combination of histological images from different sections, aspects of which have been previously described^9^. In brief, we implemented image preprocessing in Fiji, and registration procedures in R, using the image analysis package ‘imager’. Segmentation of the histological sections and associated spatial barcodes from background was achieved using a custom macro in Fiji. Segmented sections were then aligned using imager. Image registration was performed manually using the tissue structure to guide registration, as captured by (i) histological images, (ii) quality control statistics (e.g., % of mitochondrial counts), (iii) marker genes for coarse cell types and dorsoventral or rostrocaudal transcription factors (e.g., *Ebf1*, *Esrrg*, *Hox* genes), and (iv) unsupervised clustering of the spatial barcodes, as implemented within Seurat.

### Visualization

Quality control metrics and marker gene expression were smoothed prior to visualization on the two-dimensional spinal cord using locally weighted regression, as implemented in the RCTD package^167^. Visualization of the three-dimensional spinal cord was achieved with Imaris (Bitplane, version 9.0.0). Briefly, the three-dimensional spatial transcriptomics data was binned along the z-dimension into slices of 10 *µ*m. Within each slice, quantitative values (quality control metrics, gene expression, gene module scores, and chromVAR deviations) were smoothed using three-dimensional locally weighted regression. When multiple quantitative values were assigned to a single spatial coordinate (for example, when performing spatial prioritization on snRNA-seq barcodes embedded via Tangram), the mean value at each coordinate was assigned, with the exception of the expression of individual genes, for which the maximum value at each coordinate was assigned instead. Each barcode was then assigned a size of 3 pixels, and the resulting slices were exported as 16 bit grayscale TIFF files using imager for import into Imaris. Separate reconstructions of the three-dimensional spinal cord volume were performed for each experimental condition in the spatiotemporal atlas (that is, uninjured, 7 days, and 2 months).

### Differential expression

To identify genes differentially expressed between regions in the injured spinal cord within the two-dimensional spatial dataset, we extended the workflow for pseudobulk DE analysis in Libra to spatial count matrices derived by SpaceRanger. Cells from each region were aggregated within replicates to form pseudobulks, and then testing for DE was performed using the likelihood ratio test implemented in edgeR^193^. DE analysis was performed separately for spinal cord regions at 7 days and 2 months post-injury. DE gene expression was visualized on the two-dimensional spinal cord using two-dimensional locally weighted regression, as implemented in the RCTD package^167^.

Cell type deconvolution. To integrate our snRNA-seq atlas with the two- and three-dimensional spatial atlases, we used RCTD^167^ to deconvolve spatial barcodes into a mixture of one or more cell types, while accounting for technical differences between single-nucleus and spatial transcriptomes. RCTD was run with doublet mode disabled, allowing each barcode to potentially contain more than two cell types, separately for cell type definitions at level 1 and 2 of the clustering tree. We recovered smoothed patterns of cell type abundance by two-dimensional locally weighted regression of deconvolution weights, as described by the authors of RCTD^167^. Separately, a single cell type was assigned to each spatial barcode by taking the maximum deconvolution weight assigned by RCTD for that barcode. For cell type definitions at level 2 of the clustering tree, only subtypes of the assigned level 1 cell types were considered as potential matches for each spatial barcode.

### Spatial prioritization with Magellan

To characterize the spatial response to SCI in an unbiased manner, we employed a machine-learning method spatial prioritization that we recently developed, named Magellan^9^. Magellan builds on the concept of transcriptional separability that provides a basis for cell type prioritization in Augur, as described above. However, in spatial transcriptomics data, the analytical level of interest is not necessarily a cell type, but rather a coordinate within a two- or three-dimensional tissue. To approach the data at this level, we sought to evaluate the transcriptional separability between barcodes from two experimental conditions at each point within a common coordinate system. We reasoned that we could achieve this by evaluating the separability of barcodes from each condition within small, overlapping tiles, layered across the spatial coordinate system. Briefly, for each barcode in a spatial transcriptomics dataset, Magellan selects the *k* nearest neighbors from each experimental condition within common coordinate space, where *k* is set to 20 by default. Then, Magellan withholds the experimental condition labels for a proportion of these neighbors, and trains a random forest classifier to predict the experimental condition given the remaining barcodes as input. The accuracy of these predictions is evaluated in the withheld barcodes, and the process is repeated in three-fold cross-validation. As in Augur, the accuracy is quantified using the AUC. The cross-validation is repeated several times (by default, 50 times) in order to converge at a robust estimate of the AUC. The entire procedure is repeated for each barcode in the dataset, providing a spatial map of the AUC over the coordinate system of the spatial transcriptomes.

Magellan was used to perform spatial prioritization in the two-dimensional spatial dataset by comparing registered spatial transcriptomes from each pair of experimental conditions (uninjured, 7 days, 2 months). To visualize the intensity of the perturbation response, the spatial AUC was smoothed by two-dimensional locally weighted regression. In addition, we performed a one-dimensional locally weighted regression to visualize the intensity of the perturbation response along the rostrocaudal axis of the spinal cord.

To more carefully dissect the transcriptional basis of the perturbation response detected by Magellan, we tested for correlation between gene expression and the AUC of spatial prioritization. Briefly, we filtered the UMI count matrix within each comparison to include only genes detected in at least 100 spatial barcodes, and then computed Pearson correlations between scaled and log-normalized gene expression vectors and the AUCs returned for each barcode by Magellan. We further identified genes that were differentially correlated with the AUCs at 7 days and 2 months by testing for differential correlations using the Fisher z-transformation, adapting code from the DGCA R package^197^. We extended this concept by computing module scores for GO terms for each spatial barcode with the Seurat function AddModuleScore, as described above, and testing for significant correlations between GO module scores and the AUCs returned by Magellan. As in the DE analysis, the expression of genes or GO modules correlated or anticorrelated with the AUC of spatial prioritization was visualized on the two-dimensional spinal cord using two-dimensional locally weighted regression.

### Integration of the *Tabulae Paralytica*

To integrate all of the four *Tabulae* into a single framework, we leveraged Tangram^170^ to to embed singlenucleus transcriptomes and epigenomes onto the common coordinate system established by our four-dimensional atlas of the mouse spinal cord. Alignment of single-cell barcodes into the spatiotemporal atlas was performed separately for each experimental condition in the snRNA-seq and multiome atlases, using the most similar condition in the spatiotemporal atlas as a reference (e.g., aligning cells from 14 days to the spatiotemporal atlas at 7 days and cells from 1 month to the spatiotemporal atlas at 2 months). Tangram was run with the top 500 highly variable genes for each cell type and using cell type definitions at level 4 of the clustering tree.

This procedure assigned x-, y-, and z-coordinates to each nuclei in the snRNA-seq and multiome atlases. We then employed Magellan to perform three-dimensional spatial prioritization on the spatialized single-cell data, using the coordinates assigned by Tangram for each barcode. Spatialized cells from each injury severity were compared to those from uninjured mice. Separately, spatialized cells from old mice were compared to those from injured young mice at the same timepoint. Moreover, we again tested for correlation between gene expression in spatialized cells and the AUC of three-dimensional spatial prioritization.

Gene modules associated with blood-brain barrier endothelial cell identity and peripheral endothelial cell identity were obtained from the literature^101^, and their expression in spatial barcodes was calculated using the Seurat function AddModuleScore. Similarly, we used the average expression of genes associated with the GO term “GO:0031175” (neural projection development) to construct a circuit reorganization score, as described above for the snRNA-seq atlas, and visualized the expression of this score in spatialized neurons from the snRNA-seq atlas. Last, to summarize the expression of the conserved early response module in neurons, we selected the top 25 genes that were upregulated in the greatest number of level 4 neuron subtypes and with the greatest mean log-fold change at 4 days post-injury, and used the Seurat function AddModuleScore to summarize the expression of this gene module.

Transcription factor accessibility at 7 days was visualized on the four-dimensional atlas by first embedding individual nuclei from multiome atlas onto the three-dimensional coordinate system of the spinal cord, and then visualizing chromVAR deviations from linked epigenomes for each nucleus.

### Statistics, power calculations, group sizes, reproducibility, visualization

Statistical evaluations of repeated measures were conducted by one-way ANOVA with post hoc independent pairwise analysis as per Tukey’s HSD. For all photomicrographs of histological tissue, staining experiments were repeated independently with tissue from at least four, and in most cases six, different animals with similar results.

## Data availability

Sequencing data have been deposited to the Gene Expression Omnibus (GSE234774, snRNA-seq and spatial transcriptomics, and GSE230765, multiome).

## Code availability

Augur, Libra, and Magellan are available from GitHub (https://github.com/neurorestore/Augur, https://github.com/neurorestore/Libra, https://github.com/neurorestore/Magellan).

## Acknowledgements.

This work was supported by the Swiss National Science Foundation (310030_192558 to G.C., PZ00P3_185728 to M.A.A. and PZ00P3_208988 to J.W.S.); the Morton Cure Paralysis Foundation (to M.A.A); the ALARME Foundation (to M.A.A. and G.C); the Dr. Miriam and Sheldon G. Adelson Medical Foundation (to M.V.S.); Wings for Life (M.V.S., M.A.S., and M.M); Holcim-Stiftung Foundation (to J.W.S.); Canadian Institutes for Health Research (to J.W.S.); and the Human Frontiers in Science Program long-term fellowship (LT001278/2017-L to C.K.). We are grateful to Jimmy Ravier and Frederic Merlos for the illustrations; the Advanced Lightsheet Imaging Center (ALICe) at the Wyss Center for Bio and Neuroengineering, Geneva; and Dr. Elizabeth Bradbury for providing ChABC vectors. This work was supported in part using the resources and services of the Gene Expression Core Facility and the Bertarelli Platform for Gene Therapy at the School of Life Sciences of EPFL.

## Author contributions

J.W.S., J.B., M.V.S., M.A.A., M.A.S., and G.C. conceived and designed experiments. J.W.S., M.A.A., C.K., M.G., T.H.H., A.L., A.d.C., N.R., V.A., and N.D.J. conducted experiments. J.W.S., M.A.S., M.G., Q.B., and Y.Y.T. analyzed the data. B.S. contributed essential resources. J.W.S., M.A.S., M.A.A., and G.C. wrote the manuscript. All authors contributed to the editing of the manuscript.

