## Supplementary Figures for "*The Tabulae Paralytica:* Multimodal single-cell and spatial atlases of spinal cord injury"

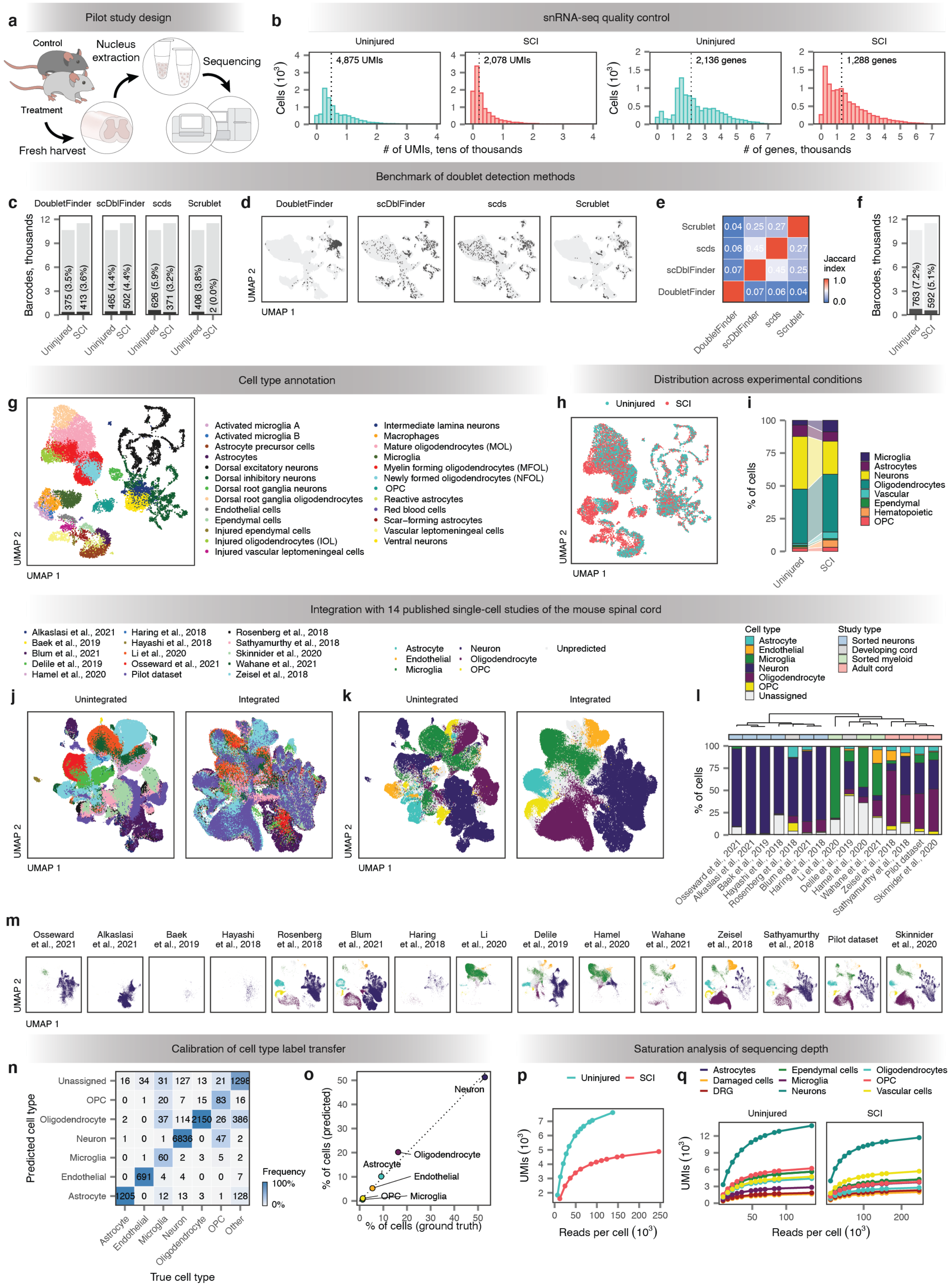

**Supplementary Fig. 1 | Pilot study of snRNA-seq in the injured mouse spinal cord.**
**a,** Schematic overview of experimental design.
**b,** Quality control (number of UMIs per cell and number of genes detected per cell) of single-cell libraries from injured and uninjured mouse spinal cords.
**c-f,** Detection and prevalence of cell doublets in the pilot study.
**c,** Number of doublets detected in each library by four computational methods for doublet detection.
**d,** Visualization of doublets detected by each of four computational methods on a UMAP plot of 22,157 cell barcodes from the mouse spinal cord.
**e,** Overlap between doublets identified by each method, as quantified by the Jaccard index. DoubletFinder calls a largely distinct set of cell barcodes as doublets, as compared to the other three methods.
**f,** Number and proportion of doublets detected in each library by the final method for doublet identification, comprising the union of doublet calls from scDblFinder and scds.
**g,** UMAP visualization of 18,269 cells, revealing 25 transcriptionally distinct clusters of cells in the injured and uninjured mouse spinal cord.
**h,** UMAP visualization of 18,269 cells, colored by the experimental condition of origin for each cell (uninjured or SCI).
**i,** Proportions of eight major cell types of the mouse spinal cord detected in either experimental condition.
**j,** UMAP visualization of 420,786 cells, including the pilot dataset and sixteen published single-cell studies of the mouse spinal cord, colored by the study of origin. Left, before data integration with Seurat; right, after data integration.
**k,** As in **j,** but showing major cell types inferred by label transfer with Seurat.
**l,** Proportions of six major cell types, and unassigned cells, recovered by sixteen published single-cell studies of the mouse spinal cord and the pilot dataset. Datasets are hierarchically clustered by cell type composition and annotated according to the study type.
**m,** As in **k,** but showing cells from each study individually.
**n-o,** Calibration of the label transfer functionality used to assign coarse cell types in published studies.
**n,** Concordance between cell types automatically assigned by label transfer and manually assigned by the authors of the original study in the Sathyamurthy et al. dataset.
**o,** Correlation between the cell type proportions recovered by label transfer and those manually assigned by the authors of the original study in the Sathyamurthy et al. dataset.
**p-q,** Analysis of the minimum sequencing depth required to detect cell-type-specific transcriptional changes in the injured mouse spinal cord.
**p,** Proportion of the complete UMI count matrix recovered after downsampling the number of reads in the injured and uninjured libraries, respectively, as compared to the complete dataset.
**q,** Proportion of the complete UMI count matrix recovered in each major cell type after downsampling the number of reads in the injured and uninjured libraries, respectively, as compared to the complete dataset.

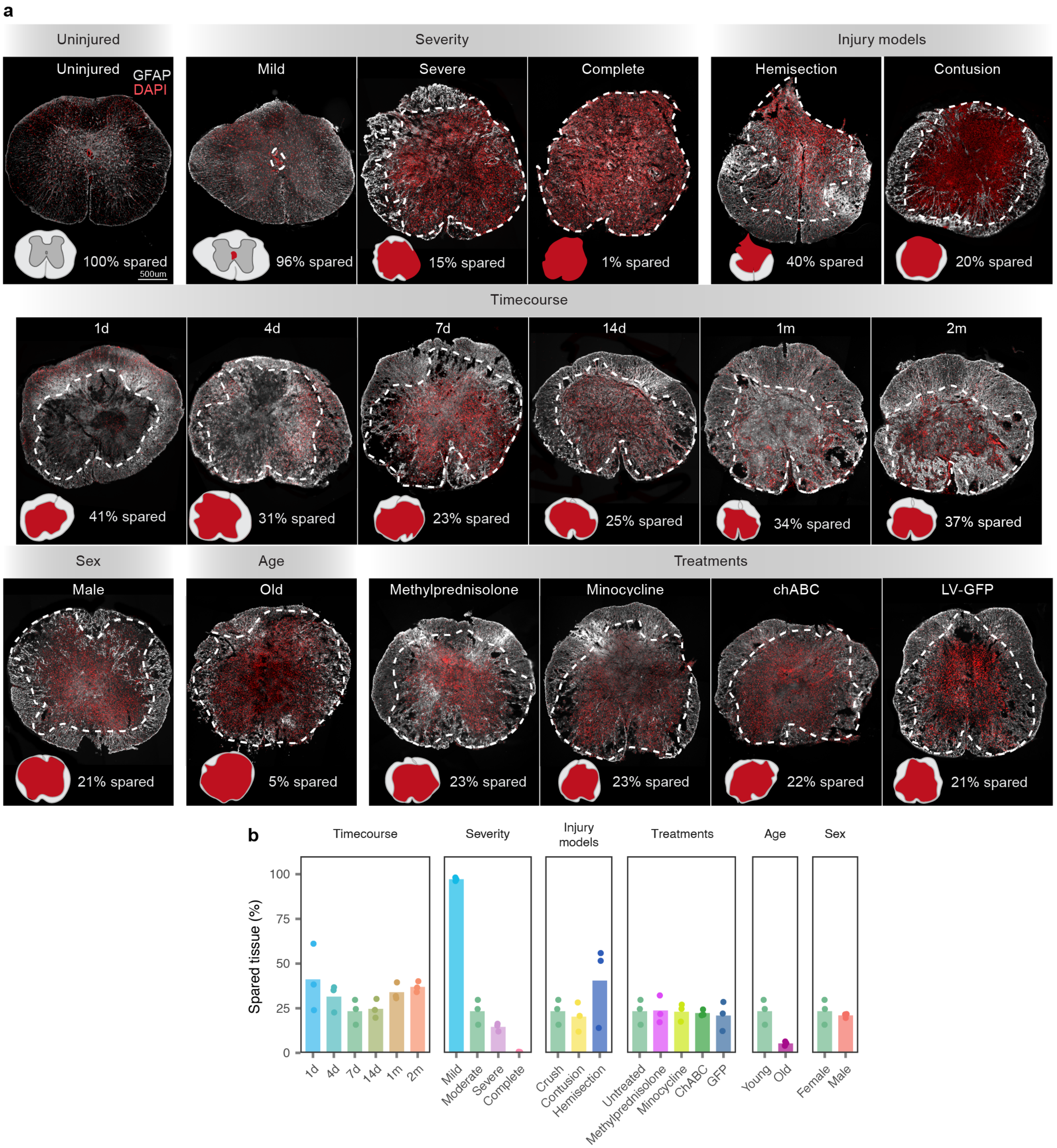


**Supplementary Fig. 2 | Representative histological images across experimental conditions.**

**a,** Histological photomicrographs show two-dimensional views of injured spinal cords stained with glial fibrillary acidic protein (GFAP).

**b,** Quantification of the SCI lesion epicentre. The extent of spared spinal cord tissue was quantified from coronal sections immunolabeled against GFAP.

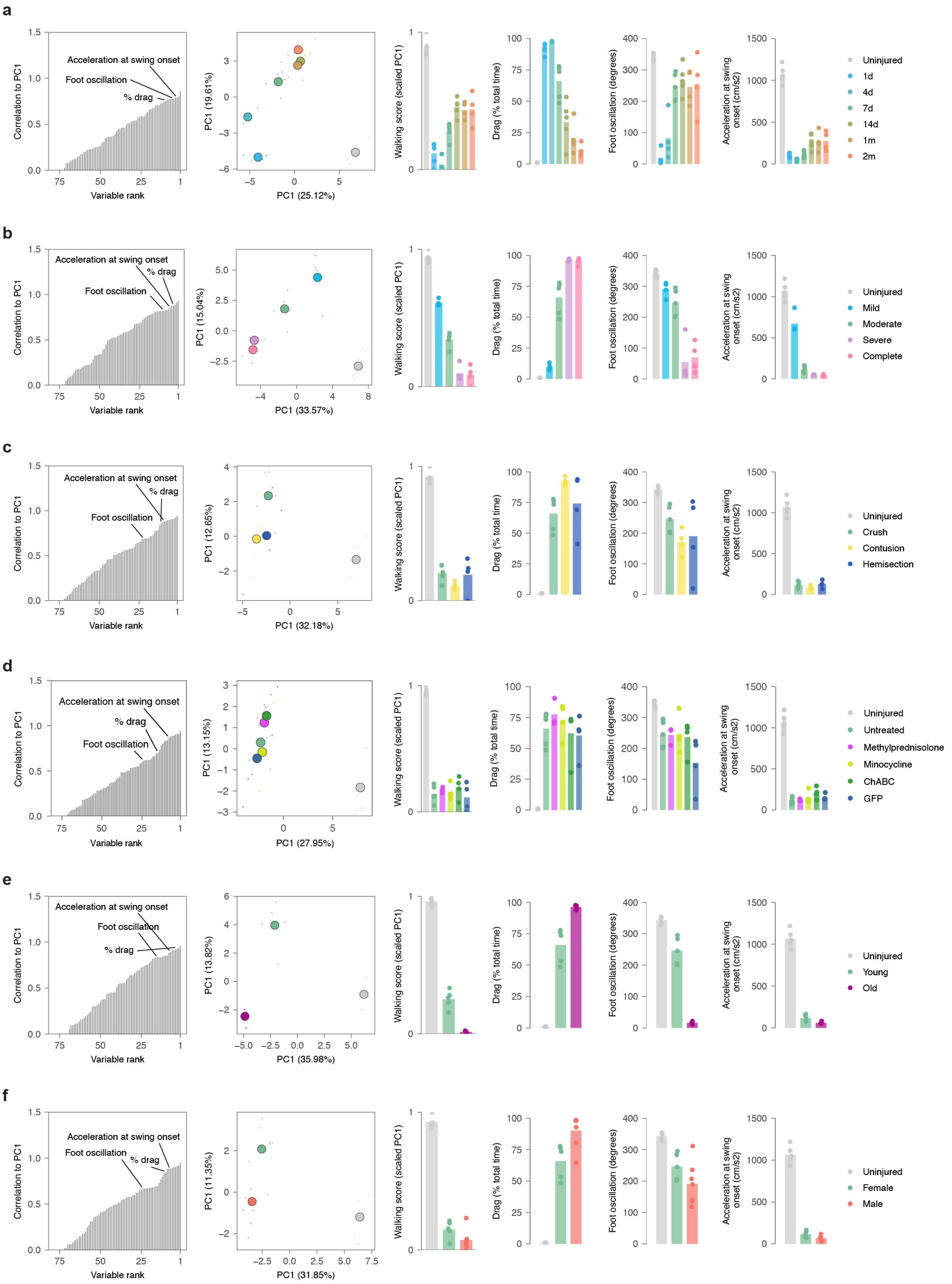

a
Correlation to PC1
Variable rank
Acceleration at swing onset
Foot oscillation
% drag
PC1 (25.12%)
PC1 (19.61%)
Walking score (scaled PC1)
Drag (% total time)
Foot oscillation (degrees)
Acceleration at swing onset (cm/s2)
Uninjured
1d
4d
7d
14d
1m
2m
b
Correlation to PC1
Variable rank
Acceleration at swing onset
% drag
Foot oscillation
PC1 (33.57%)
PC1 (15.04%)
Walking score (scaled PC1)
Drag (% total time)
Foot oscillation (degrees)
Acceleration at swing onset (cm/s2)
Uninjured
Mild
Moderate
Severe
Complete
c
Correlation to PC1
Variable rank
Acceleration at swing onset
% drag
Foot oscillation
PC1 (32.18%)
PC1 (12.65%)
Walking score (scaled PC1)
Drag (% total time)
Foot oscillation (degrees)
Acceleration at swing onset (cm/s2)
Uninjured
Crush
Contusion
Hemisection
d
Correlation to PC1
Variable rank
Acceleration at swing onset
% drag
Foot oscillation
PC1 (27.95%)
PC1 (13.15%)
Walking score (scaled PC1)
Drag (% total time)
Foot oscillation (degrees)
Acceleration at swing onset (cm/s2)
Uninjured
Untreated
Methylprednisolone
Minocycline
ChABC
GFP
e
Correlation to PC1
Variable rank
Acceleration at swing onset
Foot oscillation
% drag
PC1 (35.98%)
PC1 (13.82%)
Walking score (scaled PC1)
Drag (% total time)
Foot oscillation (degrees)
Acceleration at swing onset (cm/s2)
Uninjured
Young
Old
f
Correlation to PC1
Variable rank
Acceleration at swing onset
Foot oscillation
% drag
PC1 (31.85%)
PC1 (11.35%)
Walking score (scaled PC1)
Drag (% total time)
Foot oscillation (degrees)
Acceleration at swing onset (cm/s2)
Uninjured
Female
Male

**Supplementary Fig. 3 | High-resolution kinematic analysis across experimental conditions.**

**a,** Locomotor performance of mice across timepoints after SCI. Locomotor performance was quantified using principal component analysis applied to 55 gait parameters calculated from kinematic recordings. Small points show individual gait cycles (*n* > 10 per mouse, *n* = 4-7 mice per group). Large points show the mean of each experimental group. The first principal component (PC1) distinguished gaits from mice across different experimental groups. Analysis of factor loadings on PC1 revealed that the percentage of paw dragging, the extent of foot oscillation, and acceleration of the limb at swing onset were among the parameters that showed the highest correlation with PC1. Bars report the mean values of these gait parameters.

**b,** As in **a,** for mice subjected to different severities of crush SCI.

**c,** As in **a,** for mice subjected to different injury mechanisms.

**d,** As in **a,** for mice treated with various clinical and experimental interventions.

**e,** As in **a,** for old versus young mice.

**f,** As in **a,** for male versus female mice.

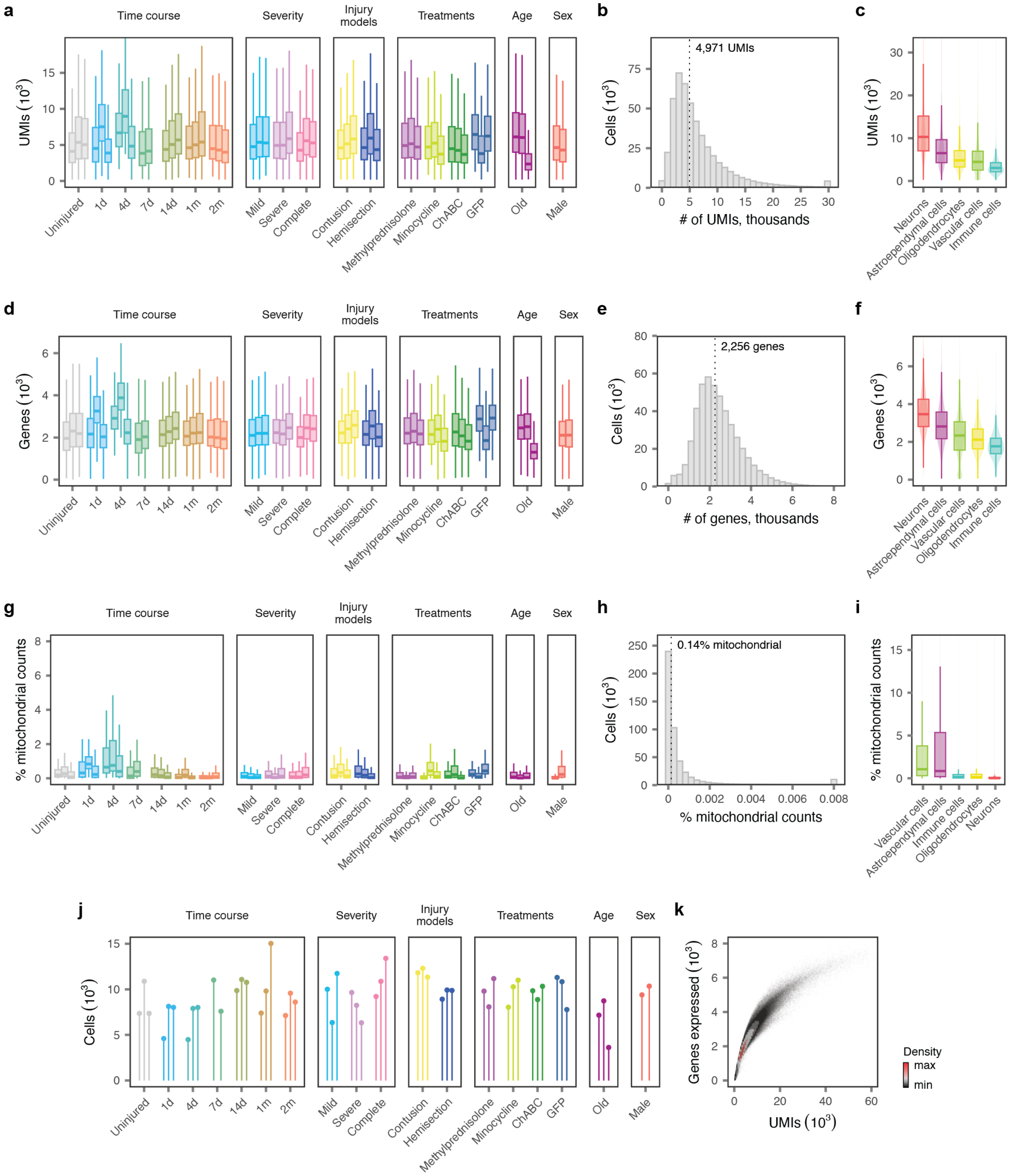

a
Time course
Severity
Injury models
Treatments
Age
Sex
UMIs (10^3)
Uninjured
1d
4d
7d
14d
1m
2m
Mild
Severe
Complete
Contusion
Hemisection
Methylprednisolone
Minocycline
ChABC
GFP
Old
Male
b
4,971 UMIs
Cells (10^3)
### of UMIs, thousands
c
UMIs (10^3)
Neurons
Astroependymal cells
Oligodendrocytes
Vascular cells
Immune cells
d
Genes (10^3)
e
2,256 genes
Cells (10^3)
### of genes, thousands
f
Genes (10^3)
g
% mitochondrial counts
h
0.14% mitochondrial
Cells (10^3)
% mitochondrial counts
i
% mitochondrial counts
j
Cells (10^3)
k
Genes expressed (10^3)
UMIs (10^3)
Density
max
min

**Supplementary Fig. 4 | Quality control of the snRNA-seq atlas.**
**a-i,** Quality control statistics for 435,099 single-nucleus transcriptomes from the uninjured and injured mouse spinal cord.
**a,** Number of unique molecular identifiers (UMIs) per nucleus in individual libraries from each experimental condition.
**b,** Number of UMIs per nucleus, aggregated across all libraries. Inset text shows the median number of UMIs.
**c,** Number of UMIs per nucleus in each major cell type.
**d,** Number of genes detected per nucleus in individual libraries from each experimental condition.
**e,** Number of genes detected per nucleus, aggregated across all libraries. Inset text shows the median number of genes detected.
**f,** Number of genes detected per nucleus in each major cell type.
**g,** Proportion of mitochondrial counts per nucleus in individual libraries from each experimental condition.
**h,** Proportion of mitochondrial counts per nucleus, aggregated across all libraries. Inset text shows the median proportion of mitochondrial counts.
**i,** Proportion of mitochondrial counts per nucleus in each major cell type.
**j,** Number of nuclei passing quality control in each of the 54 libraries comprising the snRNA-seq atlas.
**k,** Relationship between the number of UMIs and the number of genes detected per nucleus across all 435,099 nuclei.

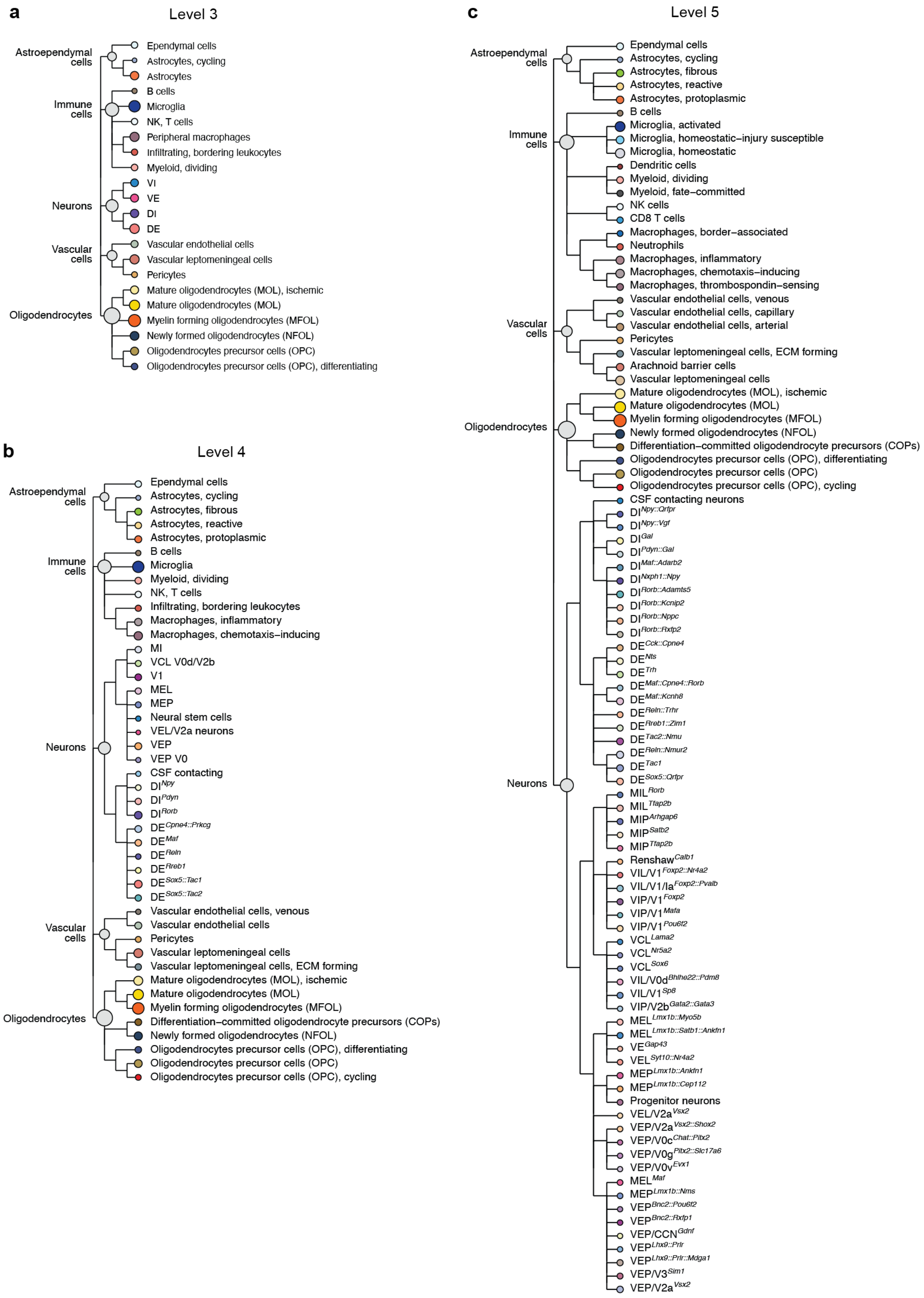


**Supplementary Fig. 5 | Clustering tree of 180 cell types and subtypes in the snRNA-seq atlas.**
**a,** Clustering tree of the mouse spinal cord, revealing the hierarchical relationships between spinal cord cell types across levels 1 to 3 in the taxonomy, with cell types at level 3 highlighted. Text at the top of the tree shows the clades of the clustering tree corresponding to the major cell types of the mouse spinal cord (i.e., level 1 in the taxonomy).
**b,** As in **a,** but showing level 4 in the taxonomy.
**c,** As in **a,** but showing level 5 in the taxonomy.

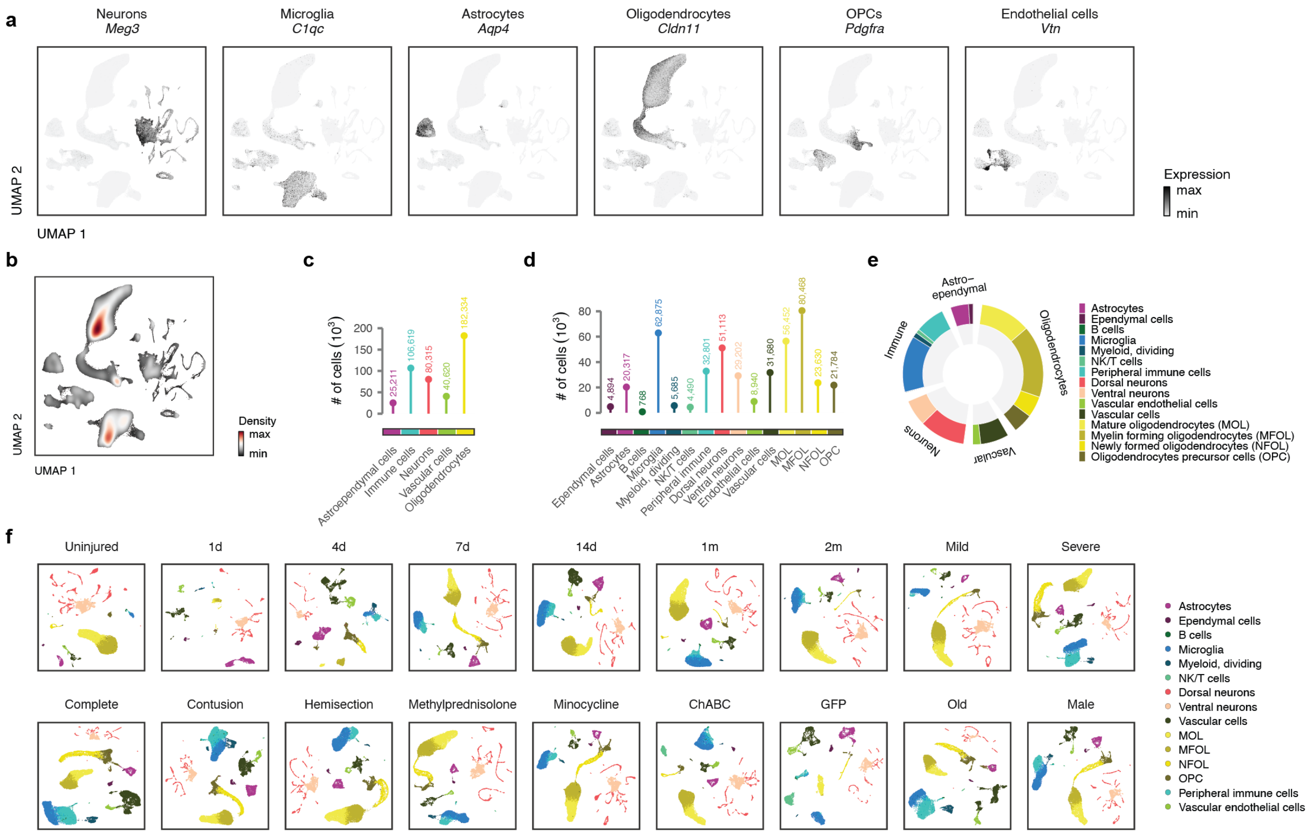


**Supplementary Fig. 6 | Annotation of major cell types in the snRNA-seq atlas.**
**a,** UMAP visualization showing expression of key marker genes for the major cell types of the mouse spinal cord.
**b,** UMAP visualization showing the density of individual nuclei within the UMAP embedding.
**c,** Number of nuclei from each major (level 1) cell type identified across all experimental conditions.
**d,** Number of nuclei from each level 2 cell type identified across all experimental conditions.
**e,** Proportions of nuclei from each level 1 (inner circle) and level 2 (outer circle) across all experimental conditions.
**f,** UMAP visualizations of each individual experimental condition in the snRNA-seq atlas, colored by level 2 cell type.

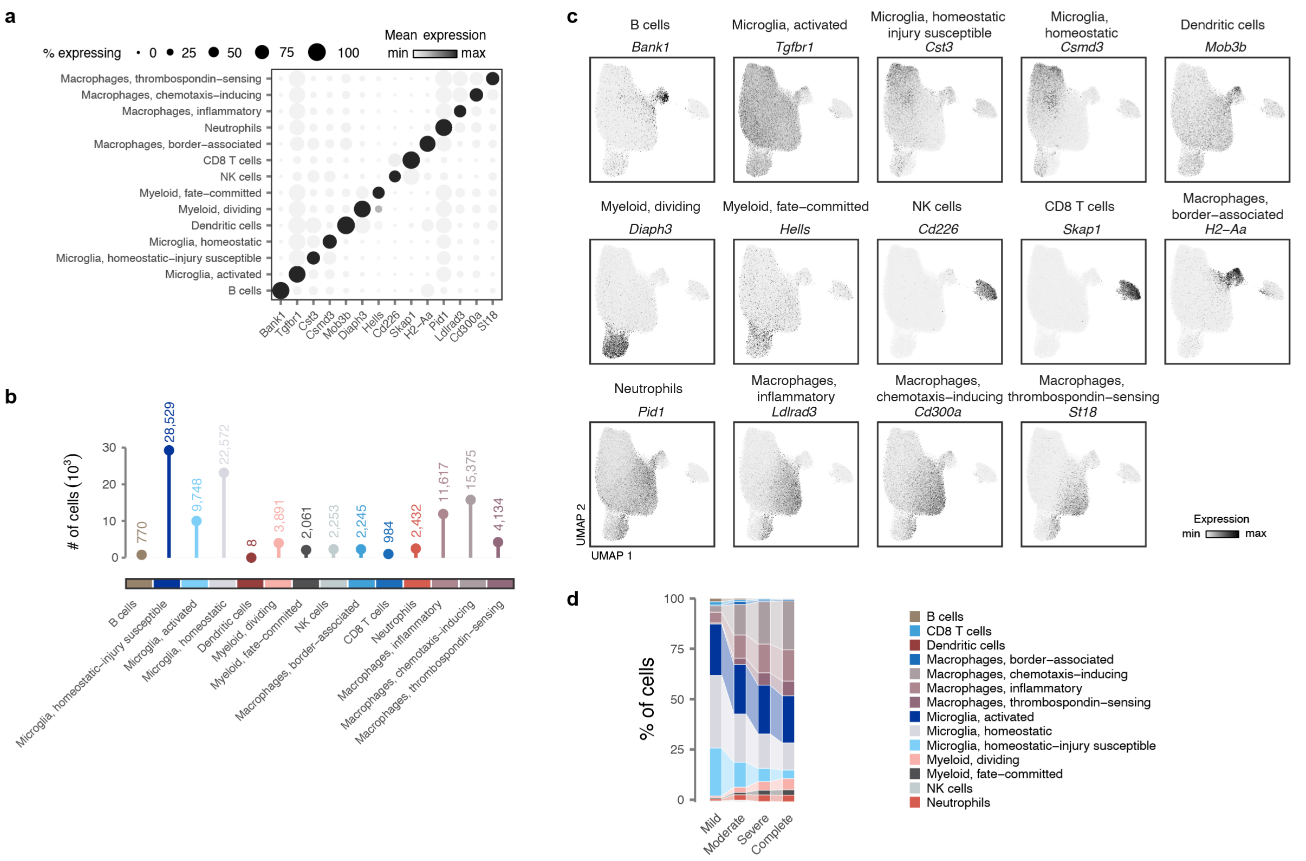


**Supplementary Fig. 7 | Annotation of immune cell subtypes in the snRNA-seq atlas.**
**a,** Dot plot showing expression of key marker genes for immune cell subtypes.
**b,** Number of nuclei from each immune cell subtype identified across all experimental conditions.
**c,** UMAP visualization showing expression of key marker genes for immune cell subtypes.
**d,** Sankey diagram showing the proportions of each astroependymal cell subtype across injury severities.

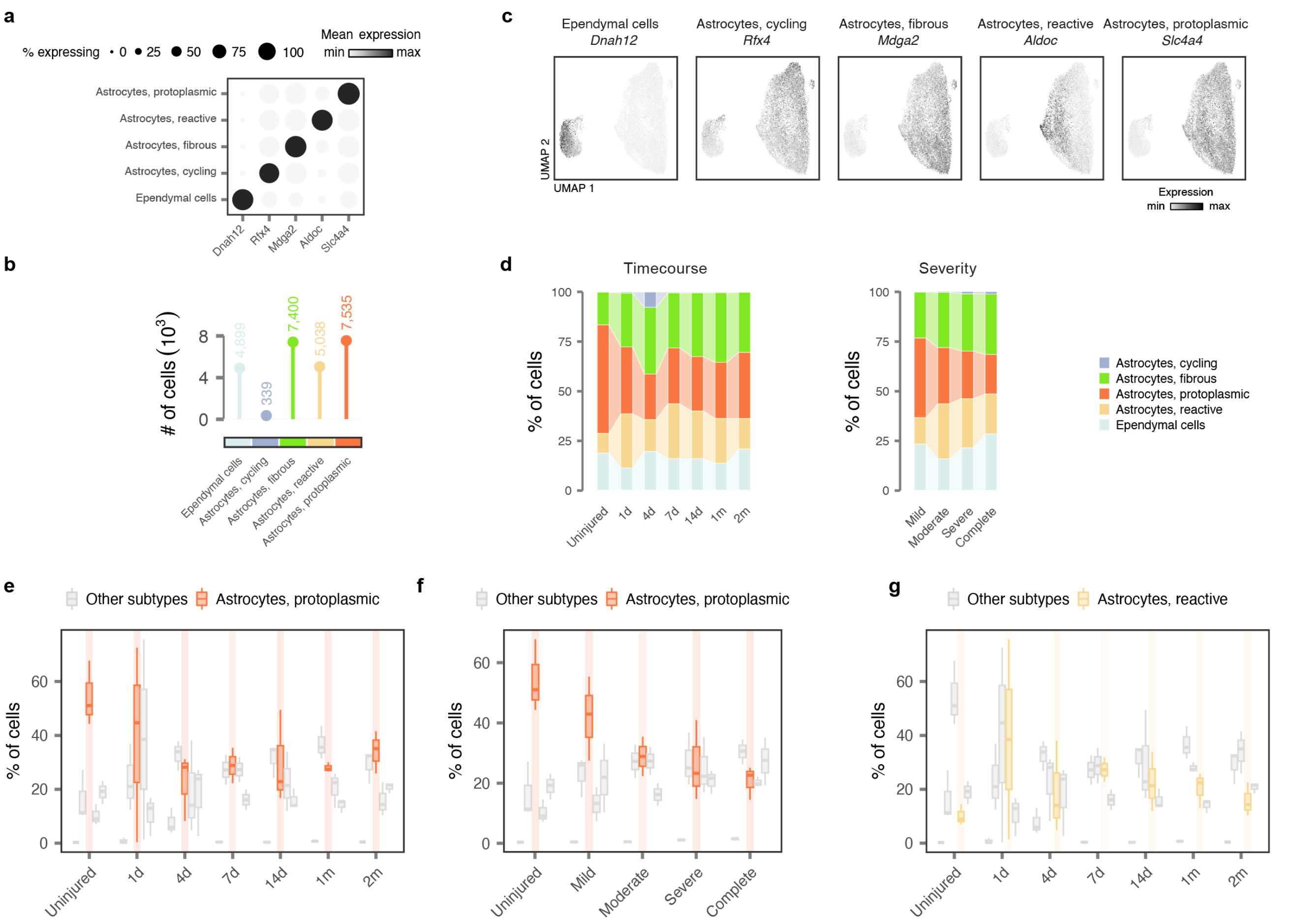


**Supplementary Fig. 8 | Annotation of astroependymal cell subtypes in the snRNA-seq atlas.**

**a,** Dot plot showing expression of key marker genes for astroependymal cell subtypes.

**b,** Number of nuclei from each astroependymal cell subtype identified across all experimental conditions.

**c,** UMAP visualization showing expression of key marker genes for astroependymal cell subtypes.

**d,** Sankey diagrams showing the proportions of each astroependymal cell subtype across timepoints, left, and injury severities, right.

**e,** Boxplot highlighting the proportions of protoplasmic astrocytes within individual libraries from the timecourse experiment, as compared to all other astroependymal cell subtypes.

**f,** Boxplot highlighting the proportion of protoplasmic astrocytes within individual libraries from the severity experiment, as compared to all other astroependymal cell subtypes.

**g,** Boxplot highlighting the proportion of reactive astrocytes within individual libraries from the timecourse experiment, as compared to all other astroependymal cell subtypes.

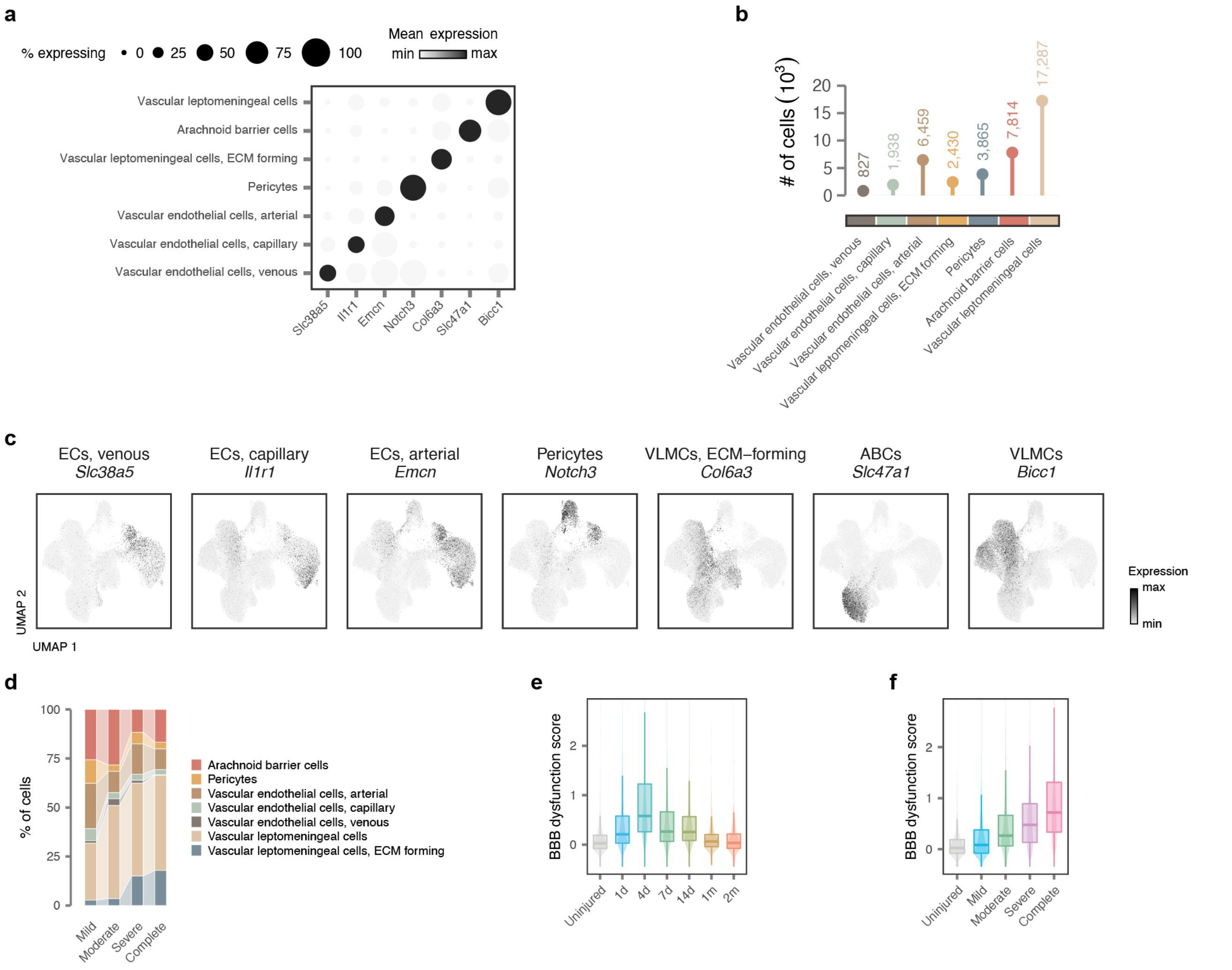


**Supplementary Fig. 9 | Annotation of vascular cell subtypes in the snRNA-seq atlas.**
**a,** Dot plot showing expression of key marker genes for vascular cell subtypes.
**b,** Number of nuclei from each vascular cell subtype identified across all experimental conditions.
**c,** UMAP visualization showing expression of key marker genes for vascular cell subtypes.
**d,** Sankey diagram showing the proportions of each vascular cell subtype across injury severities.
**e,** Average expression of the BBB dysfunction module in vascular cells across timepoints.
**f,** Average expression of the BBB dysfunction module in vascular cells across injury severities.

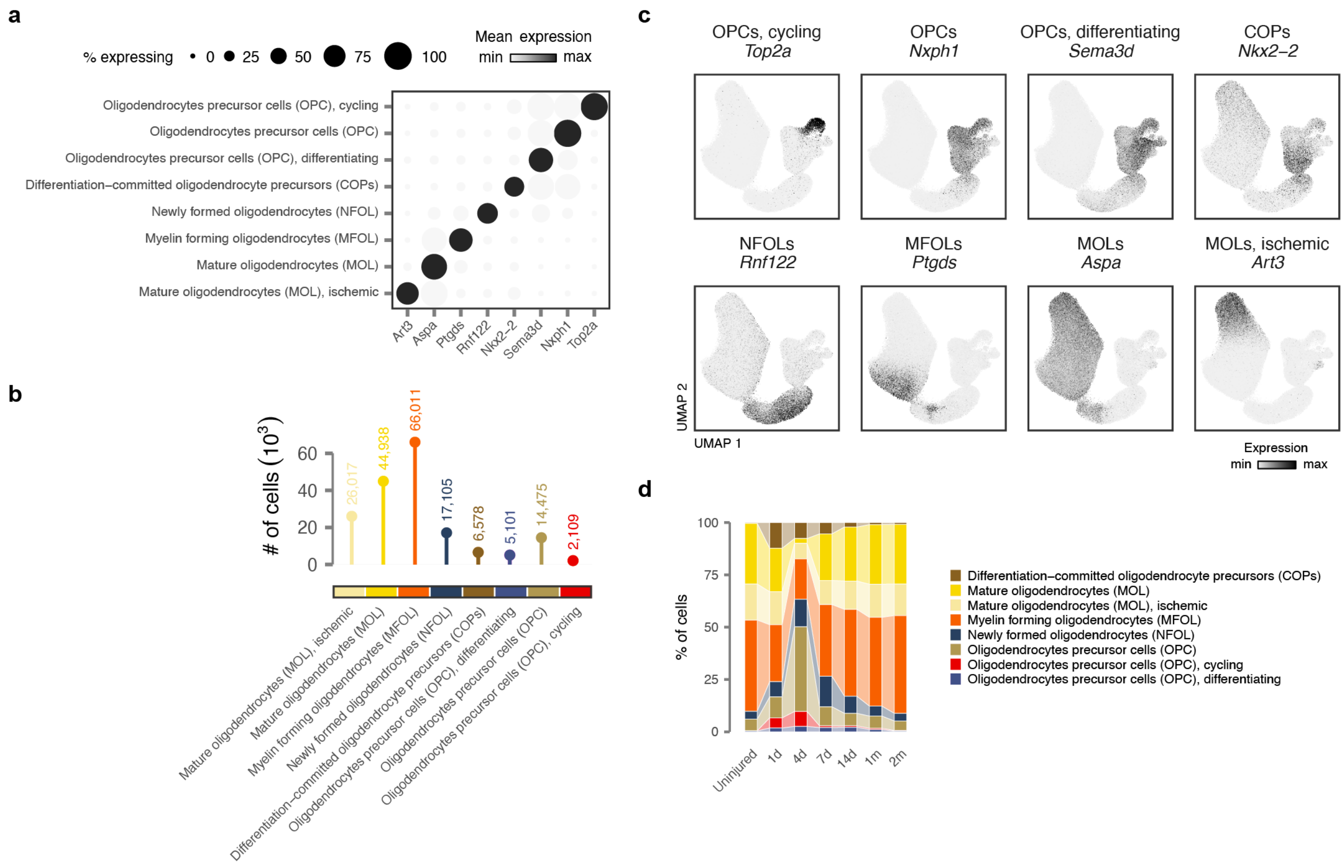


**Supplementary Fig. 10 | Annotation of oligodendrocyte subtypes in the snRNA-seq atlas.**
**a,** Dot plot showing expression of key marker genes for oligodendrocyte subtypes.
**b,** Number of nuclei from each oligodendrocyte subtype identified across all experimental conditions.
**c,** UMAP visualization showing expression of key marker genes for oligodendrocyte subtypes.
**d,** Sankey diagram showing the proportions of each oligodendrocyte subtype across timepoints.

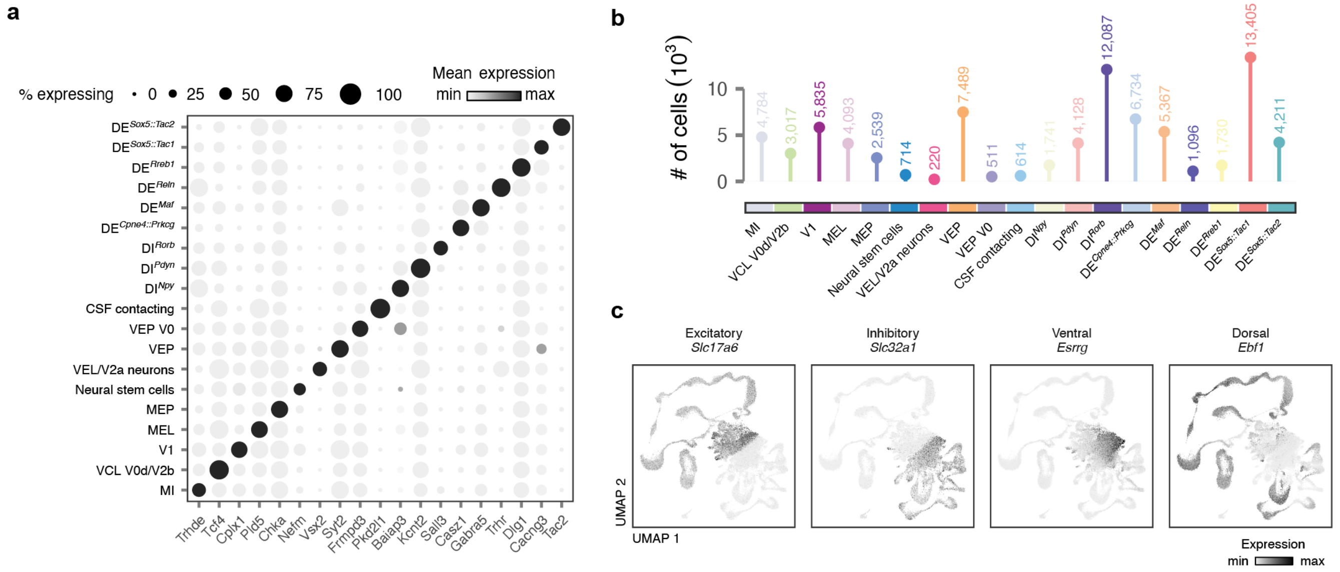


**Supplementary Fig. 11 | Annotation of neuron subtypes in the snRNA-seq atlas.**

**a,** Dot plot showing expression of one key marker gene for each level 4 neuron subtype in the mouse spinal cord.

**b,** Number of nuclei from each level 4 neuron subtype identified across all experimental conditions.

**c,** UMAP visualization showing expression of classical excitatory-inhibitory and dorsal-ventral marker genes within neurons in the mouse spinal cord.

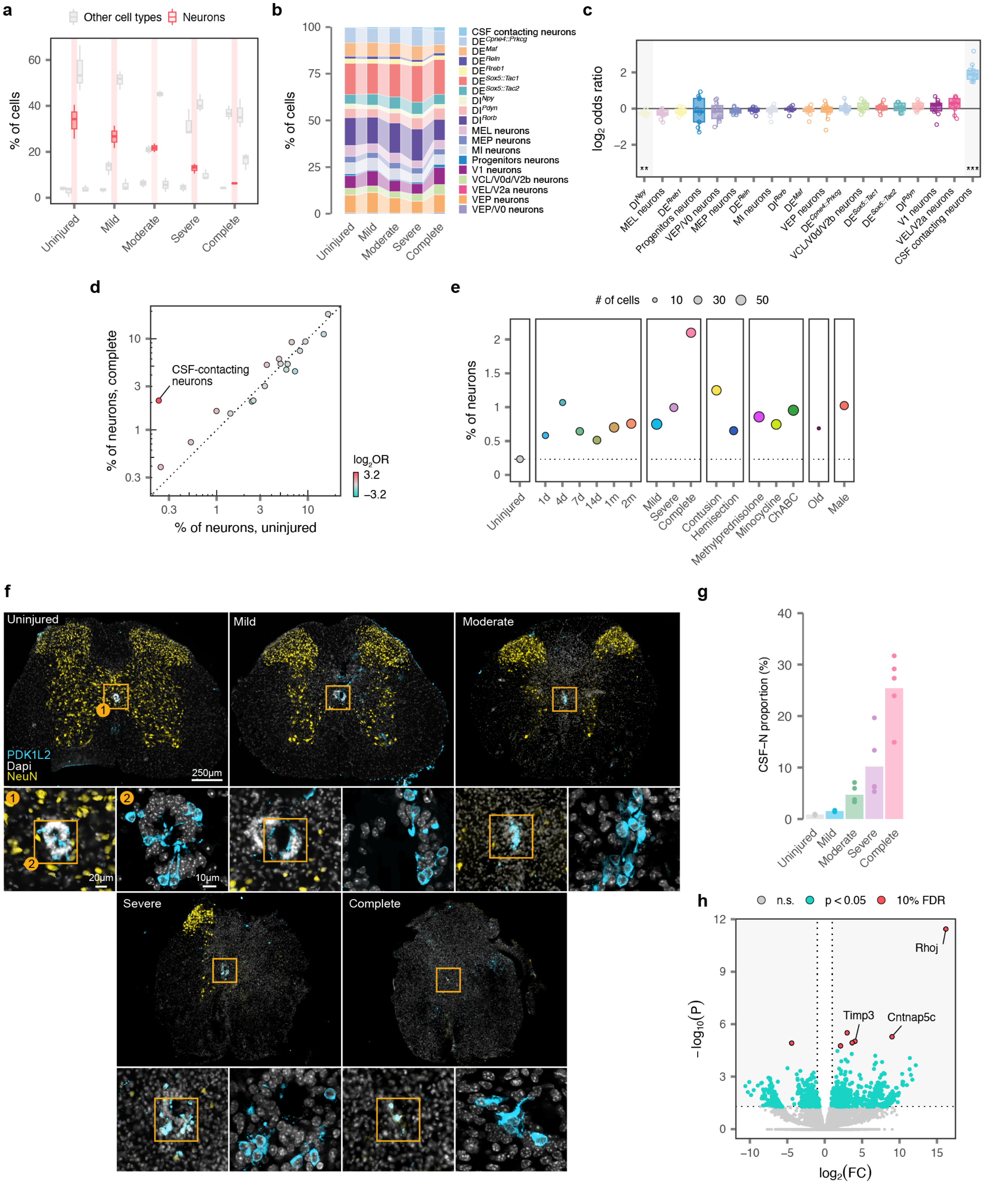

a
Other cell types
Neurons
% of cells
Uninjured
Mild
Moderate
Severe
Complete
b
% of cells
CSF contacting neurons
MEL neurons
MEP neurons
MI neurons
Progenitors neurons
V1 neurons
VCL/V0d/V2b neurons
VEL/V2a neurons
VEP neurons
VEP/V0 neurons
c
log2 odds ratio
d
CSF-contacting neurons
% of neurons, complete
% of neurons, uninjured
log2OR
e
### of cells
% of neurons
Contusion
Hemisection
Methylprednisolone
Minocycline
ChABC
Old
Male
f
PDK1L2
Dapi
NeuN
250µm
20µm
10µm
g
CSF–N proportion (%)
h
n.s.
p < 0.05
10% FDR
−log10(P)
log2(FC)
Rhoj
Timp3
Cntnap5c

**Supplementary Fig. 12 | Neuronal susceptibility and resilience to SCI.**
**a,** Boxplot highlighting the proportion of neurons within individual libraries from the severity experiment, as compared to other major cell types.
**b,** Sankey diagram showing the proportions of each level 4 neuron subtype across injury severities.
**c,** Boxplot showing the $\log_2$-odds ratio comparing the proportions of neurons from each level 4 subtype between the uninjured spinal cord, for all comparisons involving the injured spinal cord at 7 days post-injury. Cerebrospinal fluid-contacting neurons are the lone subpopulation to exhibit statistically significant resilience following SCI. **, $p < 0.01$; ***, $p < 0.001$.
**d,** Scatterplot highlighting an individual comparison from **c,** showing the proportions of neurons from each level 4 subtype in the uninjured spinal cord, x-axis, and 7 days after a complete injury, y-axis. Color shows the $\log_2$-odds ratio. Cerebrospinal fluid-contacting neurons are highlighted.
**e,** Proportion, y-axis, and absolute number, point size, of cerebrospinal fluid-contacting neurons recovered from each experimental condition. Dotted line shows the proportion of cerebrospinal fluid-contacting neurons in the uninjured spinal cord.
**f,** Representative histological photomicrographs show injured spinal cords across injury severities after staining for NeuN and PKD1L2, a marker of cerebrospinal fluid-contacting neurons.
**g,** Quantification of histological data demonstrating increasing proportions of cerebrospinal fluid-contacting neurons across injury severities.
**h,** Volcano plot showing differentially expressed genes in cerebrospinal fluid-contacting neurons following spinal cord injury, as compared to other neuron subtypes.

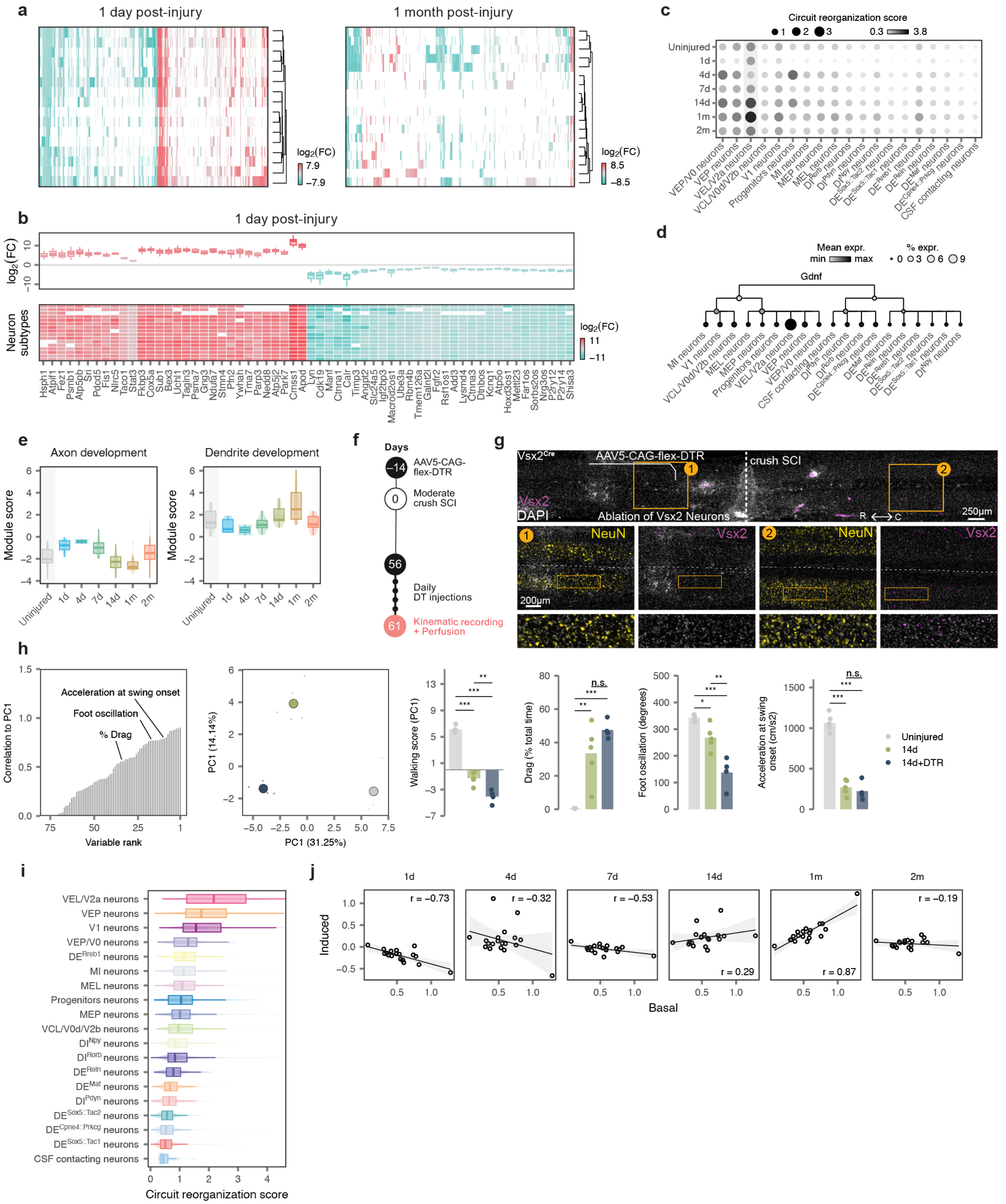
a
1 day post-injury
1 month post-injury
log2(FC)
7.9
−7.9
8.5
−8.5
b
1 day post-injury
Neuron subtypes
c
Circuit reorganization score
d
Mean expr.
% expr.
Gdnf
e
Axon development
Dendrite development
Module score
f
Days
AAV5-CAG-flex-DTR
Moderate crush SCI
Daily DT injections
Kinematic recording + Perfusion
g
crush SCI
Ablation of Vsx2 Neurons
NeuN
Vsx2
DAPI
h
Acceleration at swing onset
Foot oscillation
% Drag
Correlation to PC1
Variable rank
PC1 (31.25%)
PC2 (14.14%)
Walking score (PC1)
Drag (% total time)
Foot oscillation (degrees)
Acceleration at swing onset (cm/s2)
Uninjured
14d
14d+DTR
i
Circuit reorganization score
j
Induced
Basal

**Supplementary Fig. 13 | Conserved and divergent neuronal responses to SCI.**

**a,** Heatmap showing fold changes for all genes differentially expressed after SCI in at least one level 4 neuron subtype at 1 day, top, and 1 month, bottom, after injury. Patterns of differential expression are broadly conserved at 1 day, but more subtype-specific at 1 month.

**b,** Heatmap showing fold changes for selected genes with broadly conserved patterns of differential expression across level 4 neuron subtypes at 1 day post-injury.

**c,** Dot plot showing average expression of the circuit reorganization module in each level 4 neuron subtype across timepoints.

**d,** Dendrogram showing expression of the growth factor Gdnf across levels 1 to 4 of the neuron taxonomy. Point color shows mean expression in each neuron subtype, while point size reflects the proportion of neurons of that subtype with detectable expression.

**e,** Boxplots showing average expression of the axon development, left, and dendrite development, right modules in Vsx2-expressing neurons across timepoints.

**f,** Timeline of *Vsx2*$^{ON}$ neuron diphtheria toxin ablation experiments. Two weeks before complete crush SCI, animals received an injection of AAVs expressing DTR. At eight weeks, animals received daily injections of diphtheria toxin for 7 days. Kinematics were then recorded and tissue was collected for evaluation.

**g,** Histological verification of *Vsx2*$^{ON}$ neuron ablation in the lower thoracic region. Images show loss of *Vsx2*$^{ON}$ neurons in the thoracic spinal cord, above and below the level of the crush SCI. Bar graph shows the number of *Vsx2*$^{ON}$ neurons found in each animal ($n$ = 4 mice per group, independent samples two-tailed t-test, t = 11.7, p = $2.4 \times 10^{-5}$).

**h,** Locomotor performance in the *Vsx2*$^{ON}$ ablation experiment, as quantified in **Supplementary Fig. 2** ($n > 10$ gait cycles per mouse, $n$ = 4 mice per group; statistics indicate Tukey HSD tests following one-way ANOVA). n.s., not significant; *, $p < 0.05$; **, $p < 0.01$; ***, $p < 0.001$.

**i,** Boxplot showing average expression of the circuit reorganization module in each level 4 neuron subtype within the uninjured spinal cord. Vsx2-expressing neurons display the highest expression of the circuit reorganization module in the uninjured spinal cord.

**j,** Scatterplots comparing basal expression of the circuit reorganization module in the uninjured spinal cord, x-axis, with the SCI-induced upregulation of this module at each timepoint after injury, y-axis, for each level 4 neuron subtype. Inset text shows the Pearson correlation. Basal and induced expression of the circuit reorganization module is maximally correlated at 1 month post-injury, coinciding with the temporal window of opportunity for natural recovery after SCI.

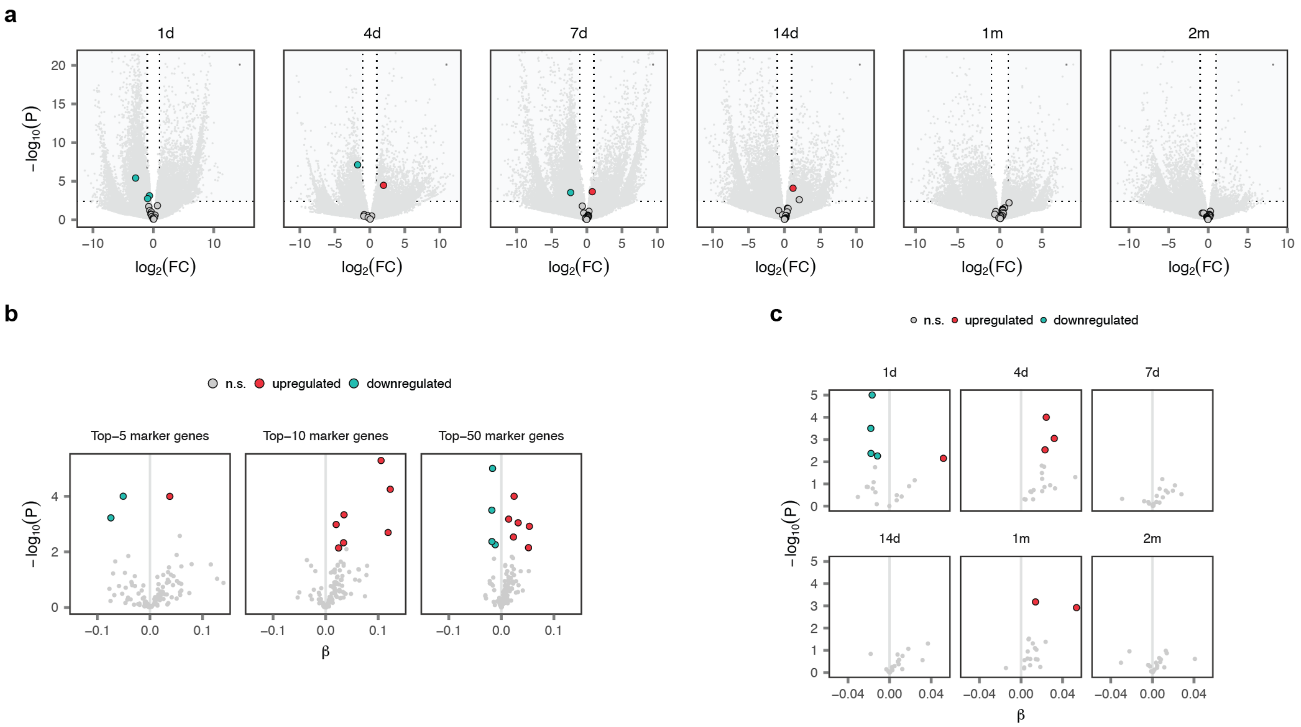


**Supplementary Fig. 14 | Neurons remain differentiated after SCI.**

**a,** Volcano plots showing differential expression for all level 4 neuron subtypes simultaneously across timepoints. Enlarged points represent the key marker genes for each neuron subtype shown in **Supplementary Fig. 11a**. Marker genes shown in grey show no evidence of differential expression after SCI in their respective neuron subtype.

**b,** Volcano plot showing differential expression for averaged gene expression modules comprising the top 5, top 10, or top 50 marker genes identified for each level 4 neuron subtype by unbiased comparisons with all other neurons, simultaneously across all timepoints. The vast majority of marker gene modules show no evidence of downregulation after SCI in their respective neuron subtype.

**c,** As in **b,** but showing differential expression for averaged gene expression modules comprising the top 50 marker genes for each level 4 neuron subtype, shown separately for each condition in the timecourse experiment.

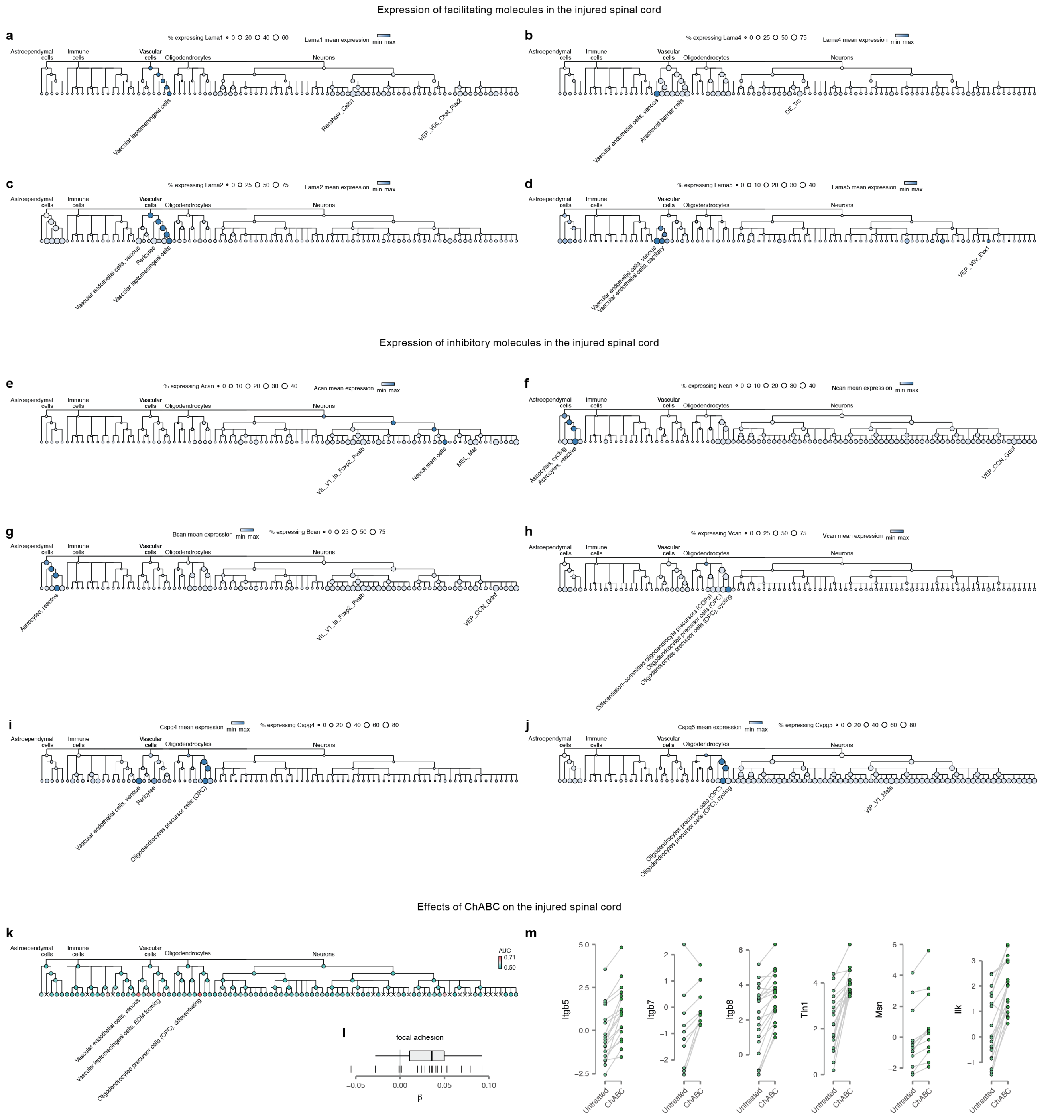


**Supplementary Fig. 15 | Facilitating and inhibiting molecule expression in the injured spinal cord.**

**a-j,** Dendrograms showing expression of selected facilitating and inhibiting molecules across the cellular taxonomy of the spinal cord. Point color shows mean expression in each cell type, while point size reflects the proportion of cells with detectable expression.

**k,** Dendrogram showing cell type prioritizations assigned by Augur across the cellular taxonomy of the spinal cord after treatment with ChABC. The three level 5 cell types with the highest AUCs are annotated.

**l,** Upregulation of genes associated with focal adhesion across level 4 neuron subtypes after ChABC treatment. Vertical lines show coefficients estimated by a linear mixed model within each neuron subtype.

**m,** Differential expression of individual genes associated with focal adhesion across level 4 neuron subtypes after ChABC treatment. Points show pseudobulk gene expression values in counts per million. Lines connect each neuron subtype across experimental conditions.

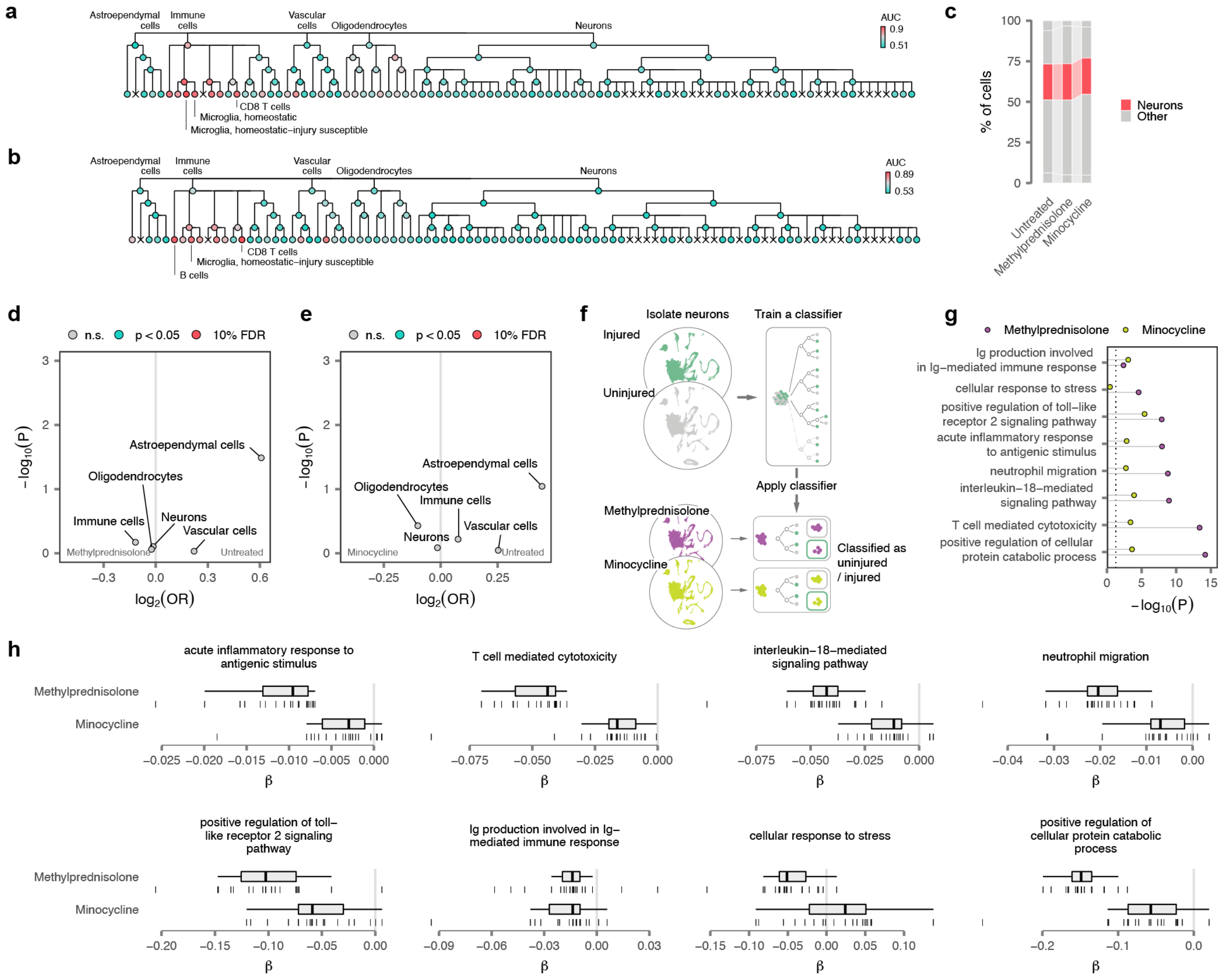


**Supplementary Fig. 16 | Immunomodulation does not confer neuroprotection after spinal cord injury.**

**a,** Dendrogram showing cell type prioritizations assigned by Augur across the cellular taxonomy of the spinal cord after treatment with methylprednisolone. The three level 5 cell types with the highest AUCs are annotated.

**b,** Dendrogram showing cell type prioritizations assigned by Augur across the cellular taxonomy of the spinal cord after treatment with minocycline. The three level 5 cell types with the highest AUCs are annotated.

**c,** Sankey diagram showing the proportions of neurons recovered from the injured spinal cord after treatment with methylprednisolone or minocycline, as compared to untreated controls.

**d,** Volcano plot showing the statistical significance of changes in major cell type proportions after treatment with methylprednisolone, as compared to untreated controls. No statistically significant differences in cell type proportions are detected after treatment.

**e,** Volcano plot showing the statistical significance of changes in major cell type proportions after treatment with minocycline, as compared to untreated controls. No statistically significant differences in cell type proportions are detected after treatment.

**f,** Schematic depicting the procedure by which neurons are assigned to uninjured versus injured transcriptional phenotypes by a machine-learning classifier.

**g,** Lollipop plot showing the statistical significance of downregulation for each of the gene modules shown in in **h** after methylprednisolone or minocycline treatment.

**h,** Downregulation of gene modules associated with cellular stress across level 4 neuron subtypes after methylprednisolone or minocycline treatment. Vertical lines show coefficients estimated by a linear mixed model within each neuron subtype.

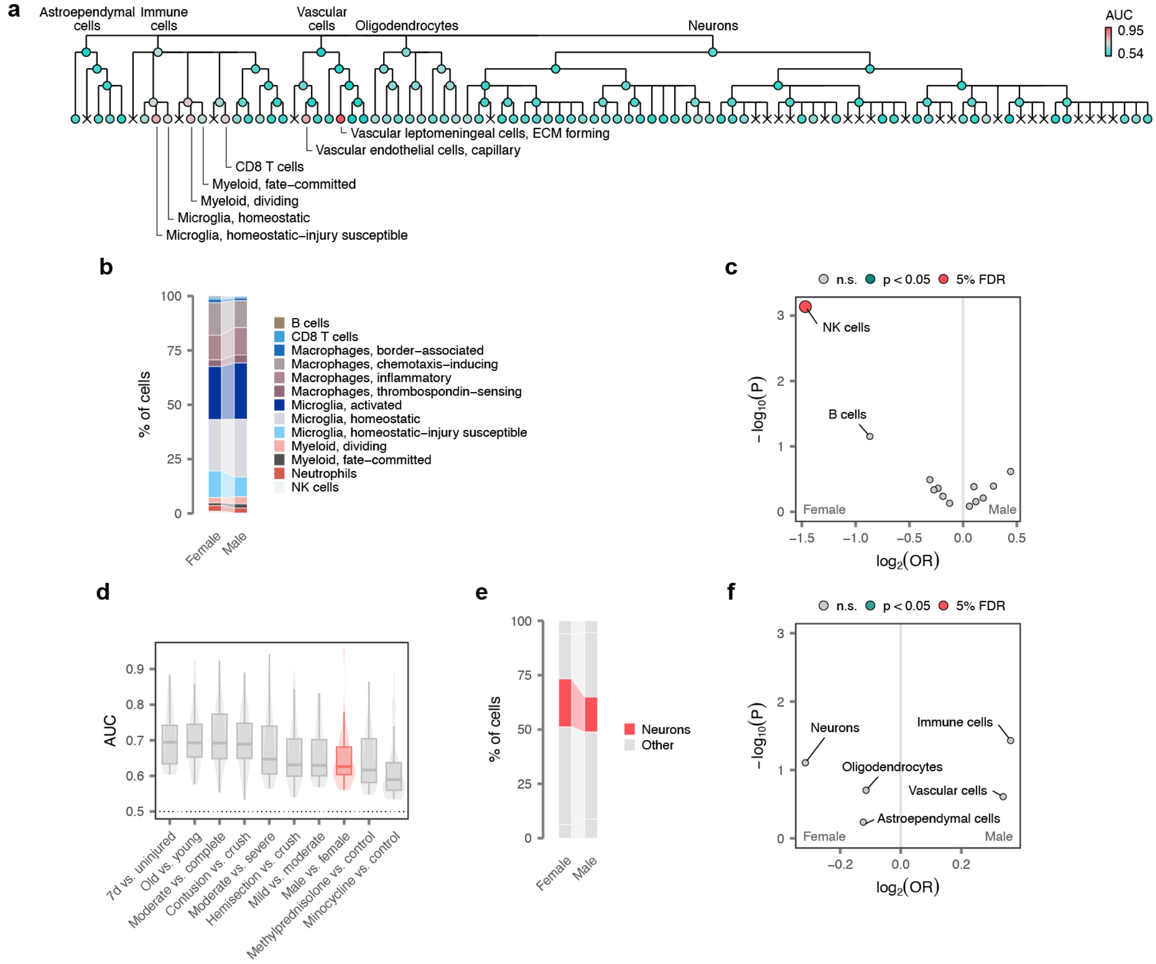


**Supplementary Fig. 17 | Sexual dimorphism in the response to SCI.**

**a,** Dendrogram showing cell type prioritizations assigned by Augur across the cellular taxonomy of the spinal cord in comparisons of male and female mice. The seven level 5 cell types with the highest AUCs are annotated.

**b,** Sankey diagram showing the proportions of immune cell subtypes recovered from the spinal cords of male versus female mice.

**c,** Volcano plot showing the statistical significance of changes in immune cell subtype proportions between male and female mice.

**d,** Boxplot showing the intensity of transcriptional perturbations within level 4 cell types, as quantified by Augur, in all comparisons involving the injured spinal cord at 7 days post-injury. The transcriptional perturbation between male and female mice is among the most subtle in the snRNA-seq atlas.

**e,** Sankey diagram showing the proportions of major cell types recovered from the spinal cords of male versus female mice.

**f,** Volcano plot showing the statistical significance of changes in major cell type proportions between male and female mice. No statistically significant differences in cell type proportions are detected.

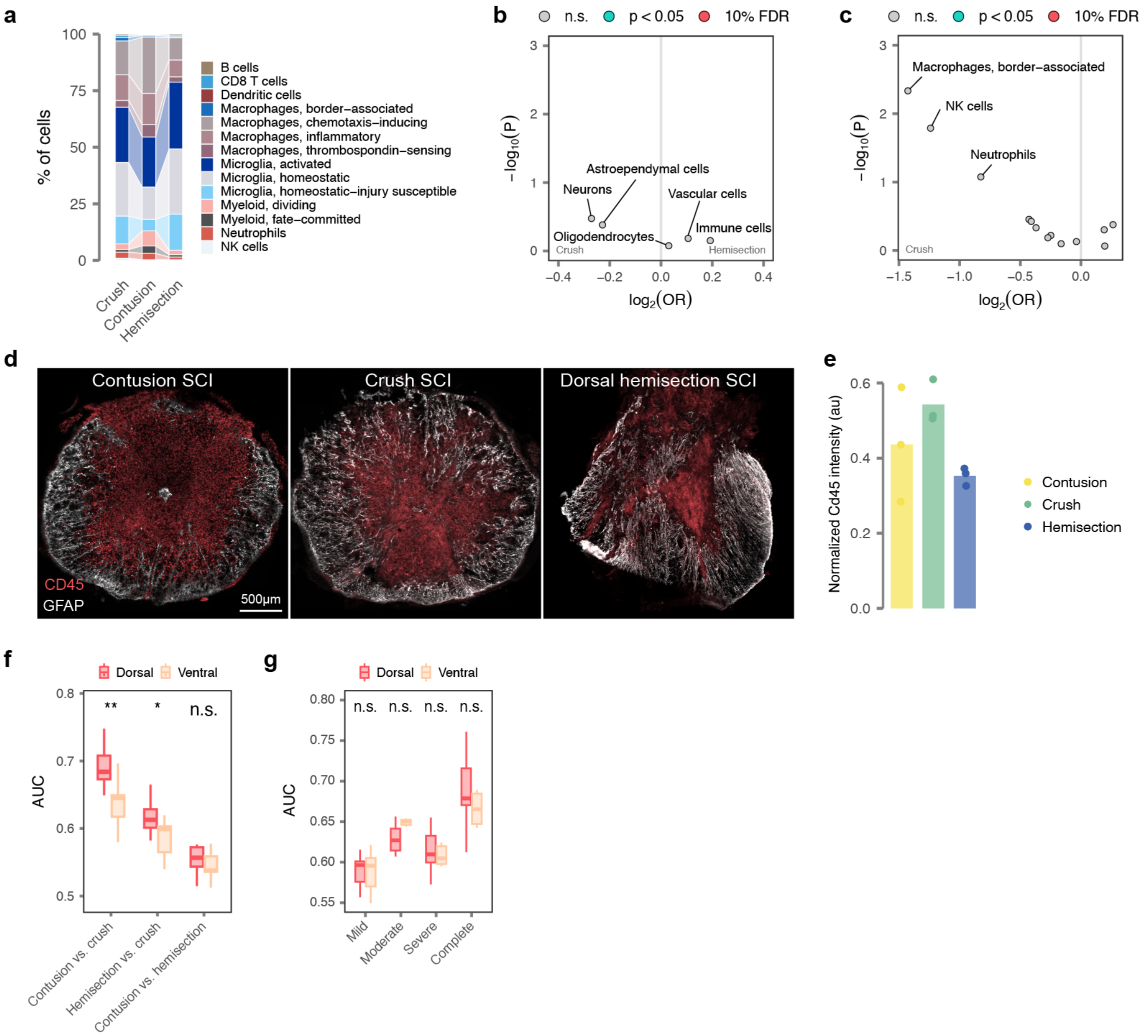


**Supplementary Fig. 18 | Cellular divergence between animal models of SCI.**

**a,** Sankey diagram showing the proportions of immune cell subtypes across crush, contusion, and hemisection injury models.

**b,** Volcano plot showing the statistical significance of changes in major cell type proportions between crush and hemisection injuries. No statistically significant differences in cell type proportions are detected.

**c,** Volcano plot showing the statistical significance of changes in immune cell subtype proportions between crush and hemisection injuries. No statistically significant differences in cell type proportions are detected.

**d,** Histological photomicrographs show injured spinal cords across injury models after staining for GFAP and CD45.

**e,** Quantification of immune infiltration at the lesion epicentre across injury models. The extent of immune infiltration was quantified from coronal sections immunolabeled for CD45. No comparisons were significant (all $p > 0.05$, Tukey HSD tests following one-way ANOVA).

**f,** Boxplots showing the average intensity of transcriptional perturbations in dorsal versus ventral neurons, as quantified by Augur, in comparisons of mice subjected to crush, contusion, and hemisection injuries. *, $p < 0.05$; **, $p < 0.01$; n.s., not significant.

**g,** Boxplots showing the average intensity of transcriptional perturbations in dorsal versus ventral neurons, as quantified by Augur, in comparisons of mice subjected to crush injuries of increasing severity. n.s., not significant.

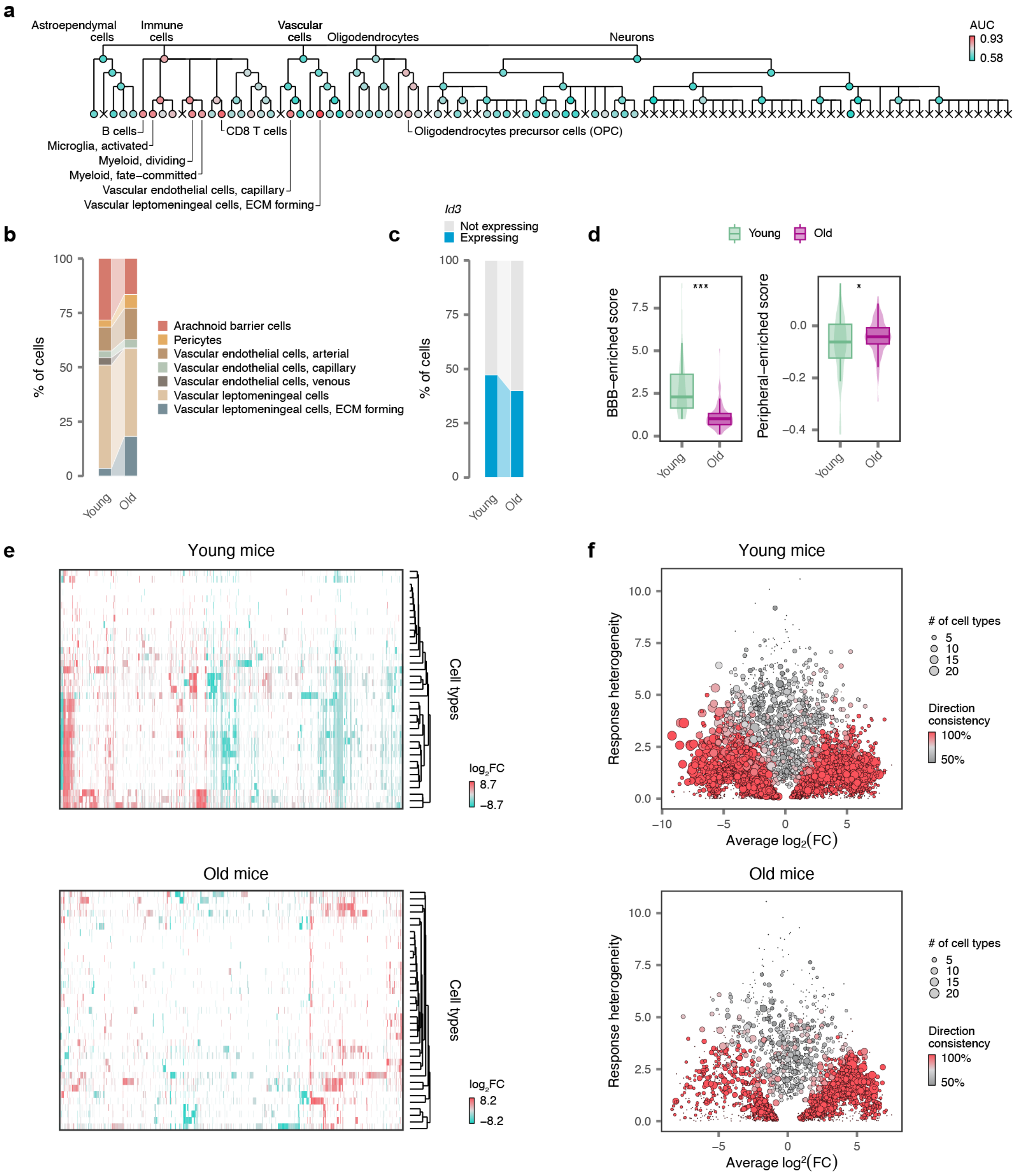
a
Astroependymal cells
Immune cells
Vascular cells
Oligodendrocytes
Neurons
AUC
0.93
0.58
B cells
Microglia, activated
Myeloid, dividing
Myeloid, fate–committed
Vascular endothelial cells, capillary
Vascular leptomeningeal cells, ECM forming
CD8 T cells
Oligodendrocytes precursor cells (OPC)
b
% of cells
Arachnoid barrier cells
Pericytes
Vascular endothelial cells, arterial
Vascular endothelial cells, capillary
Vascular endothelial cells, venous
Vascular leptomeningeal cells
Vascular leptomeningeal cells, ECM forming
Young
Old
c
Id3
Not expressing
Expressing
% of cells
d
Young
Old
BBB–enriched score
Peripheral–enriched score
***
*
e
Young mice
Old mice
Cell types
log2FC
8.7
–8.7
8.2
–8.2
f
Young mice
Old mice
Response heterogeneity
Average log2(FC)
### of cell types
5
10
15
20
Direction consistency
100%
50%

**Supplementary Fig. 19 | Catastrophic failure of tripartite barrier formation in old mice.**
**a,** Dendrogram showing cell type prioritizations assigned by Augur across the cellular taxonomy of the spinal cord in comparisons of young and old mice at seven days post-injury. The eight level 5 cell types with the highest AUCs are annotated.
**b,** Sankey diagram showing the proportions of vascular cell subtypes in young and old mice at seven days post-injury.
**c,** Sankey diagram showing the proportions of astroependymal cells expressing *Id3* in young and old mice at seven days post-injury.
**d,** Average expression of the BBB identity module, left, and the peripheral vascular identity module, right, in capillary endothelial cells from young and old mice at seven days post-injury.
**e,** Heatmap showing log-fold changes for all genes differentially expressed in at least one level 4 cell type in comparisons of injured versus uninjured mice, top, and old versus young mice, bottom.
**f,** Heterogeneity of differential expression in comparisons of injured versus uninjured mice, top, and old versus young mice, bottom. Each point shows a gene differentially expressed in at least one level 4 cell type. The x-axis shows the average log2-fold change across all cell types; the y-axis shows the standard deviation of the $\log_2$-fold change across cell types ("response heterogeneity"); point size reflects the total number of cell types in which the gene is differentially expressed; and point color reflects the proportion of cell types in which the sign of the log2-fold change was the same as the modal sign ("direction consistency").

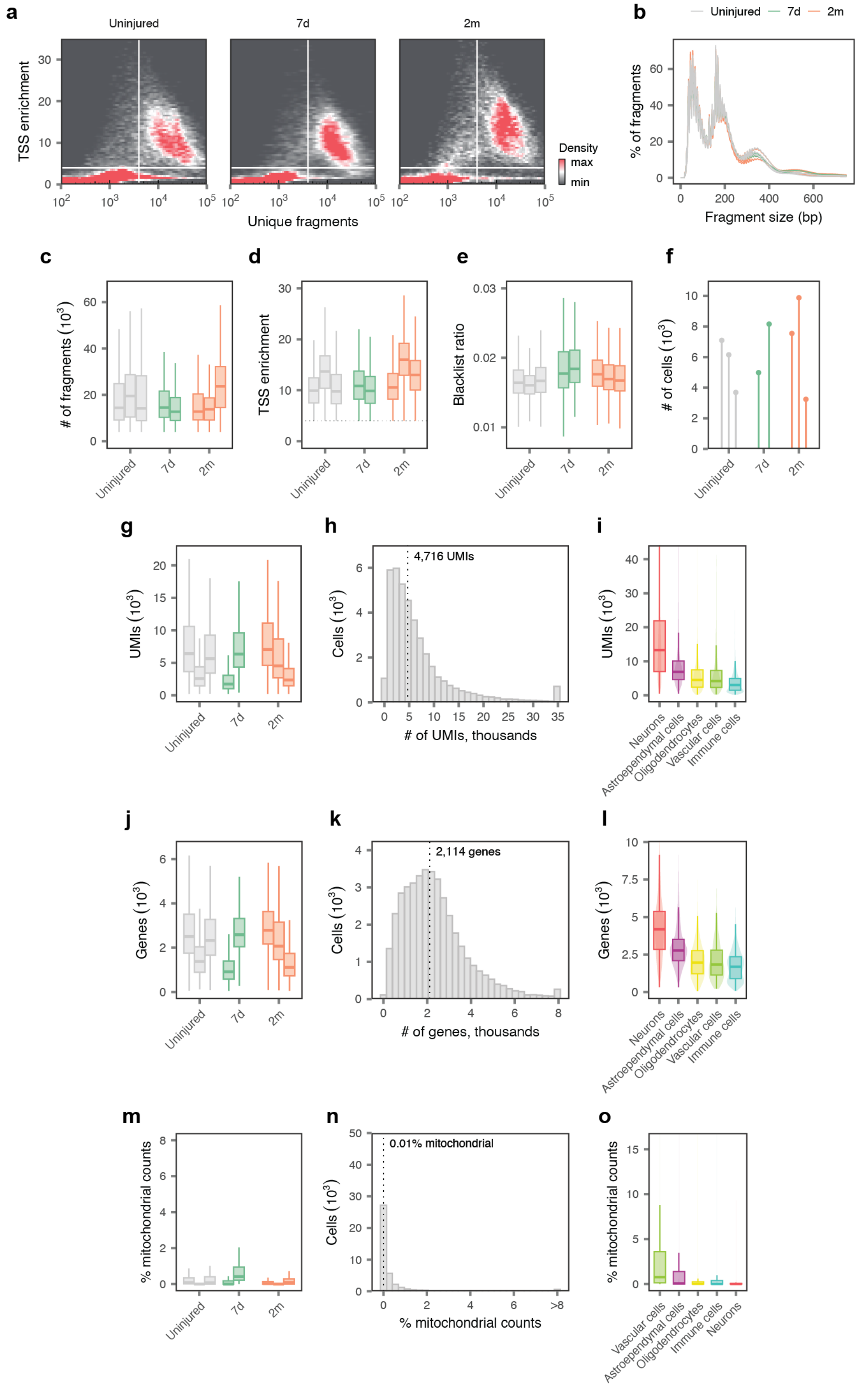

a
Uninjured
7d
2m
TSS enrichment
Unique fragments
Density
max
min
b
Uninjured 7d 2m
% of fragments
Fragment size (bp)
c
### of fragments (10³)
d
TSS enrichment
e
Blacklist ratio
f
### of cells (10³)
g
UMIs (10³)
h
4,716 UMIs
Cells (10³)
### of UMIs, thousands
i
UMIs (10³)
Neurons
Astroependymal cells
Oligodendrocytes
Vascular cells
Immune cells
j
Genes (10³)
k
2,114 genes
Cells (10³)
### of genes, thousands
l
Genes (10³)
m
% mitochondrial counts
n
0.01% mitochondrial
Cells (10³)
% mitochondrial counts
o
% mitochondrial counts

**Supplementary Fig. 20 | Quality control of the multiome atlas.**
**a-f,** Quality control of the ATAC modality.
**a,** TSS enrichment score versus the number of unique fragments per cell in barcodes from each experimental condition. Solid lines show the thresholds used for initial quality control of the ATAC modality in the multiome atlas.
**b,** Aggregate fragment size distributions for barcodes passing initial quality control thresholds in the ATAC modality of the multiome atlas.
**c,** Number of fragments per nucleus in individual libraries from each experimental condition.
**d,** TSS enrichment scores per nucleus in individual libraries from each experimental condition.
**e,** Proportion of reads mapping to blacklist regions per nucleus in individual libraries from each experimental condition.
**f,** Number of nuclei passing quality control for each individual library in the multiome atlas.
**g-o,** Quality control of the RNA modality.
**g,** Number of unique molecular identifiers (UMIs) per nucleus in individual libraries from each experimental condition.
**h,** Number of UMIs per nucleus, aggregated across all libraries. Inset text shows the median number of UMIs.
**i,** Number of UMIs per nucleus in each major cell type.
**j,** Number of genes detected per nucleus in individual libraries from each experimental condition.
**k,** Number of genes detected per nucleus, aggregated across all libraries. Inset text shows the median number of genes detected.
**l,** Number of genes detected per nucleus in each major cell type.
**m,** Proportion of mitochondrial counts per nucleus in individual libraries from each experimental condition.
**n,** Proportion of mitochondrial counts per nucleus, aggregated across all libraries. Inset text shows the median proportion of mitochondrial counts.
**o,** Proportion of mitochondrial counts per nucleus in each major cell type.

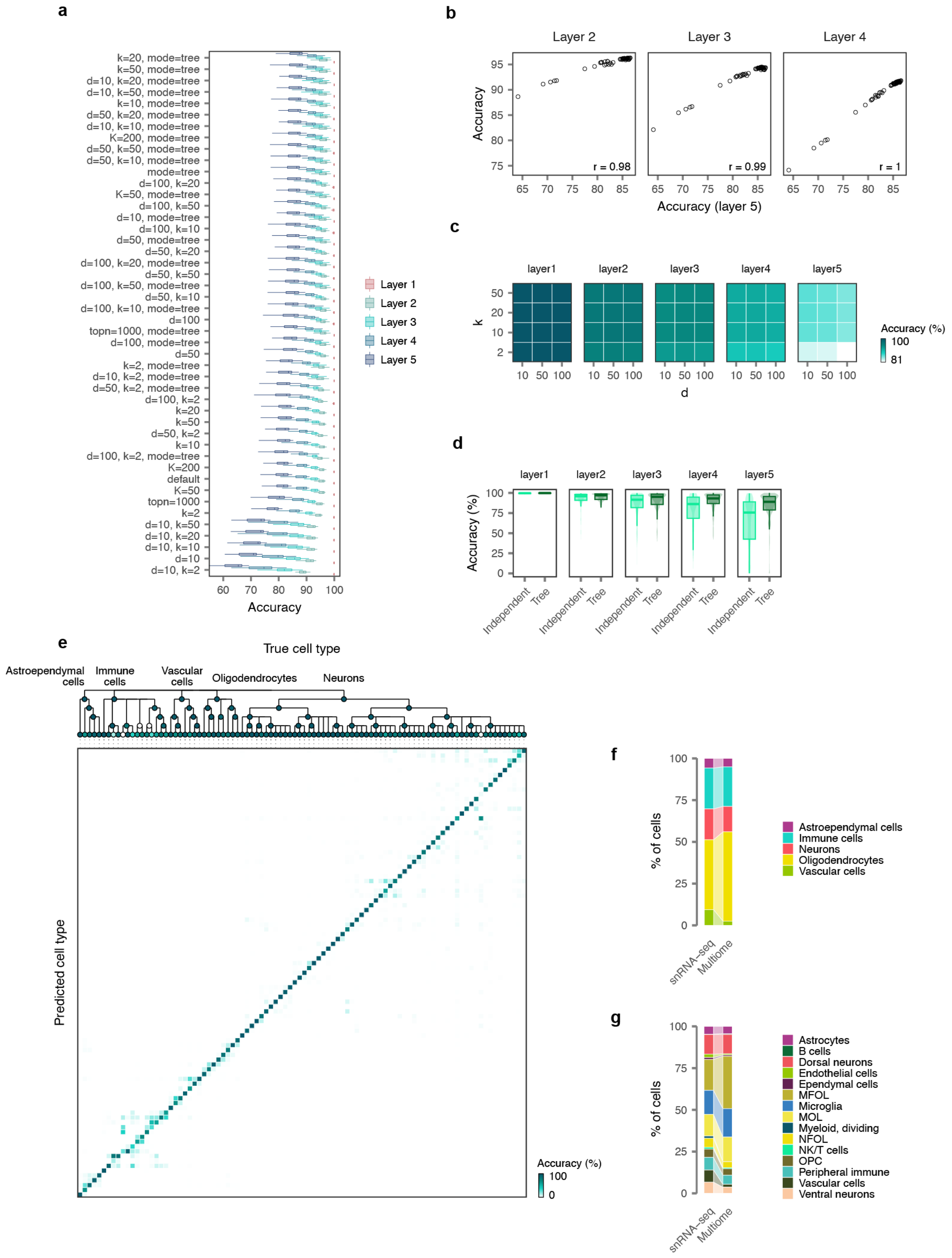
a
k=20, mode=tree
k=50, mode=tree
d=10, k=20, mode=tree
d=10, k=50, mode=tree
k=10, mode=tree
d=50, k=20, mode=tree
d=10, k=10, mode=tree
K=200, mode=tree
d=50, k=50, mode=tree
d=50, k=10, mode=tree
mode=tree
d=100, k=20
K=50, mode=tree
d=100, k=50
d=10, mode=tree
d=100, k=10
d=50, mode=tree
d=50, k=20
d=100, k=20, mode=tree
d=50, k=50
d=100, k=50, mode=tree
d=50, k=10
d=100, k=10, mode=tree
d=100
topn=1000, mode=tree
d=100, mode=tree
d=50
k=2, mode=tree
d=10, k=2, mode=tree
d=50, k=2, mode=tree
d=100, k=2
k=20
k=50
d=50, k=2
k=10
d=100, k=2, mode=tree
K=200
default
K=50
topn=1000
k=2
d=10, k=50
d=10, k=20
d=10, k=10
d=10
d=10, k=2
60 70 80 90 100
Accuracy
Layer 1
Layer 2
Layer 3
Layer 4
Layer 5
b
Layer 2
Layer 3
Layer 4
Accuracy
r = 0.98
r = 0.99
r = 1
Accuracy (layer 5)
c
layer1
layer2
layer3
layer4
layer5
k
d
Accuracy (%)
100
81
d
layer1
layer2
layer3
layer4
layer5
Accuracy (%)
Independent
Tree
e
True cell type
Astroependymal cells
Immune cells
Vascular cells
Oligodendrocytes
Neurons
Predicted cell type
Accuracy (%)
100
0
f
% of cells
snRNA–seq
Multiome
Astroependymal cells
Immune cells
Neurons
Oligodendrocytes
Vascular cells
g
% of cells
snRNA–seq
Multiome
Astrocytes
B cells
Dorsal neurons
Endothelial cells
Ependymal cells
MFOL
Microglia
MOL
Myeloid, dividing
NFOL
NK/T cells
OPC
Peripheral immune
Vascular cells
Ventral neurons

**Supplementary Fig. 21 | Hierarchical transfer of cell type annotations from the snRNA-seq atlas to the multiome atlas.**
**a,** Benchmarking the accuracy of automated cell type assignment by leave-library-out cross-validation on the snRNA-seq atlas. Boxplot shows per-cell-type accuracies in each held-out library after systematically varying the number of nearest-neighbors, k; the number of principal components, d; and the number of soft cluster centroids, K. In addition, we experimented with constraining the potential matches at finer levels of our cellular taxonomy on the basis of cell types assigned at higher levels (“mode=tree”).
**b,** Correlation between the accuracy of cell type annotation at the finest level of the clustering tree (level 5) and levels 2, 3, and 4, for each hyperparameter combination shown in a. The results for level 1 are not shown because accuracy at this level approximated 100**c,** Impact of varying the the number of nearest-neighbors, k, and the number of principal components, d, on the accuracy of automated cell type assignment in leave-library-out cross-validation.
**d,** Impact of the hierarchical adaptation of cell type annotation transfer, whereby all possible cell types are considered as potential matches at a given resolution (“independent”), or potential matches are constrained based on the cell type annotation at the previous resolution of the clustering tree (“tree”). The hierarchical approach consistently improves accuracy, particularly for lower levels of the clustering tree.
**e,** Performance of automated cell type annotation under the specific parameters used for cell type annotation in the multiome dataset. Top, dendrogram showing the average accuracy of cell type annotation for the complete cellular taxonomy in leave-library-out cross-validation on the snRNA-seq atlas. Bottom, confusion matrix at level 5 of the cellular taxonomy.
**f,** Sankey diagram showing the proportions of major (level 1) cell types in the snRNA-seq versus multiome atlases.
**g,** Sankey diagram showing the proportions of level 2 cell types in the snRNA-seq versus multiome atlases.

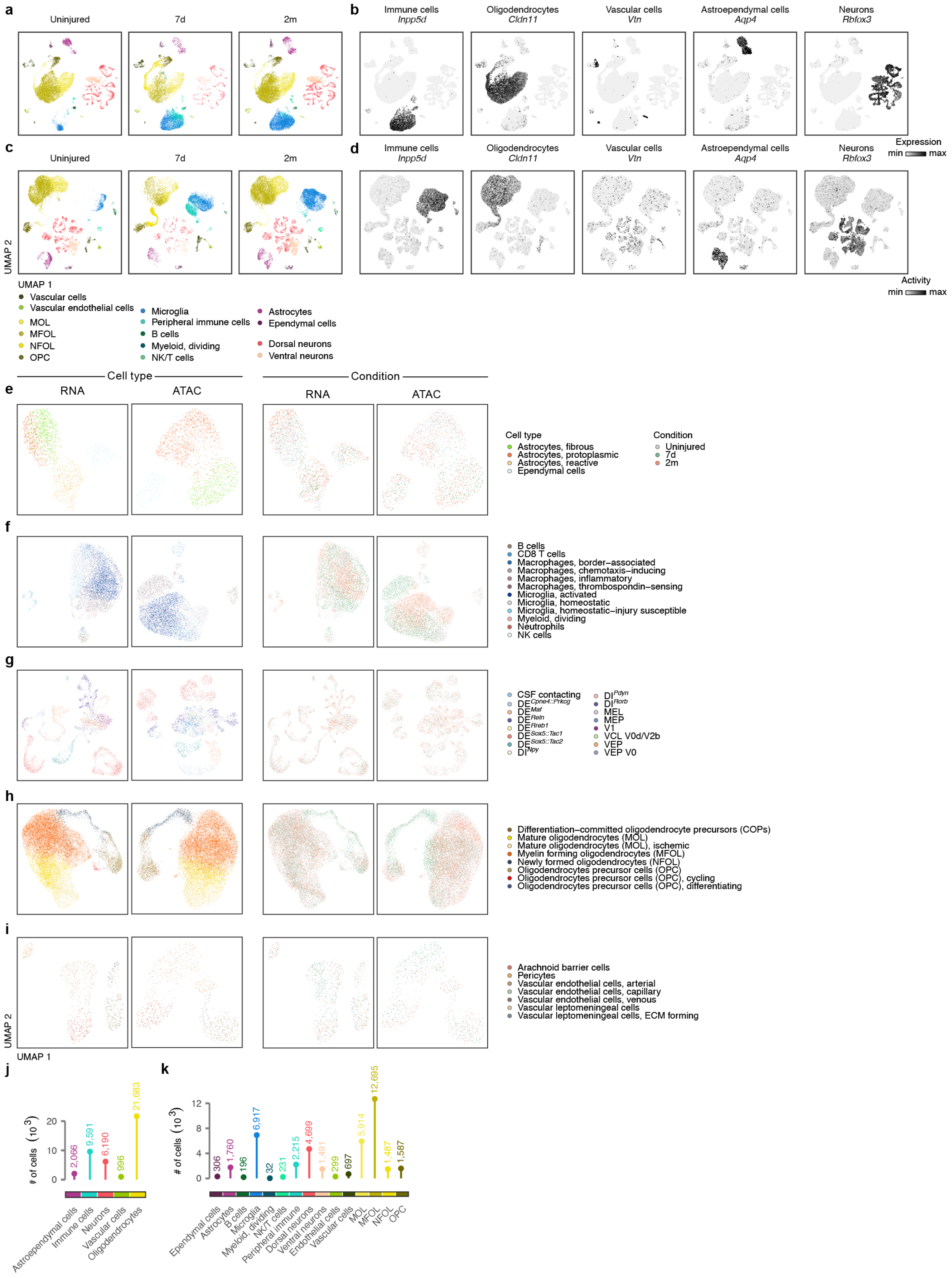
a
Uninjured
7d
2m
b
Immune cells Inpp5d
Oligodendrocytes Cldn11
Vascular cells Vtn
Astroependymal cells Aqp4
Neurons Rbfox3
Expression
min max
c
d
Activity
UMAP 2
UMAP 1
Vascular cells
Vascular endothelial cells
MOL
MFOL
NFOL
OPC
Microglia
Peripheral immune cells
B cells
Myeloid, dividing
NK/T cells
Astrocytes
Ependymal cells
Dorsal neurons
Ventral neurons
Cell type
Condition
RNA
ATAC
e
Cell type
Astrocytes, fibrous
Astrocytes, protoplasmic
Astrocytes, reactive
Ependymal cells
Condition
Uninjured
7d
2m
f
B cells
CD8 T cells
Macrophages, border–associated
Macrophages, chemotaxis–inducing
Macrophages, inflammatory
Macrophages, thrombospondin–sensing
Microglia, activated
Microglia, homeostatic
Microglia, homeostatic–injury susceptible
Myeloid, dividing
Neutrophils
NK cells
g
CSF contacting
DE Cpne4::Prkcg
DE Maf
DE Reln
DE Rreb1
DE Sox5::Tac1
DE Sox5::Tac2
DI Npy
DI Pdyn
DI Rorb
MEL
MEP
V1
VCL V0d/V2b
VEP
VEP V0
h
Differentiation–committed oligodendrocyte precursors (COPs)
Mature oligodendrocytes (MOL)
Mature oligodendrocytes (MOL), ischemic
Myelin forming oligodendrocytes (MFOL)
Newly formed oligodendrocytes (NFOL)
Oligodendrocytes precursor cells (OPC)
Oligodendrocytes precursor cells (OPC), cycling
Oligodendrocytes precursor cells (OPC), differentiating
i
Arachnoid barrier cells
Pericytes
Vascular endothelial cells, arterial
Vascular endothelial cells, capillary
Vascular endothelial cells, venous
Vascular leptomeningeal cells
Vascular leptomeningeal cells, ECM forming
j
### of cells (10^3)
2,066
9,591
6,190
996
21,683
Astroependymal cells
Immune cells
Neurons
Vascular cells
Oligodendrocytes
k
306
1,760
196
6,917
32
231
2,215
4,699
1,491
299
697
5,914
12,695
1,487
1,587
Ependymal cells
Astrocytes
B cells
Microglia
Myeloid, dividing
NK/T cells
Peripheral immune
Dorsal neurons
Ventral neurons
Endothelial cells
Vascular cells
MOL
MFOL
NFOL
OPC

**Supplementary Fig. 22 | Cell types and subtypes in the multiome atlas.**
**a,** UMAP visualization showing nuclei separately for each experimental condition in the multiome atlas, based on gene expression in the RNA modality and colored by level 2 cell type.
**b,** UMAP visualization showing expression of key marker genes for major cell types in the RNA modality of the multiome atlas.
**c,** UMAP visualization showing nuclei separately for each experimental condition in the multiome atlas, based on chromatin accessibility in the ATAC modality and colored by level 2 cell type.
**d,** UMAP visualization showing gene activity scores of key marker genes for major cell types in the ATAC modality of the multiome atlas.
**e,** UMAP visualization of immune cells in the multiome atlas based on gene expression in the RNA modality or chromatin accessibility in the ATAC modality, and colored by level 2 cell type or experimental condition.
**f,** As in **e,** but for astroependymal cells.
**g,** As in **e,** but for vascular cells.
**h,** As in **e,** but for oligodendrocytes.
**i,** As in **e,** but for neurons.
**j,** Number of nuclei from each major (level 1) cell type identified in the multiome atlas.
**k,** Number of nuclei from each level 2 cell type identified in the multiome atlas.

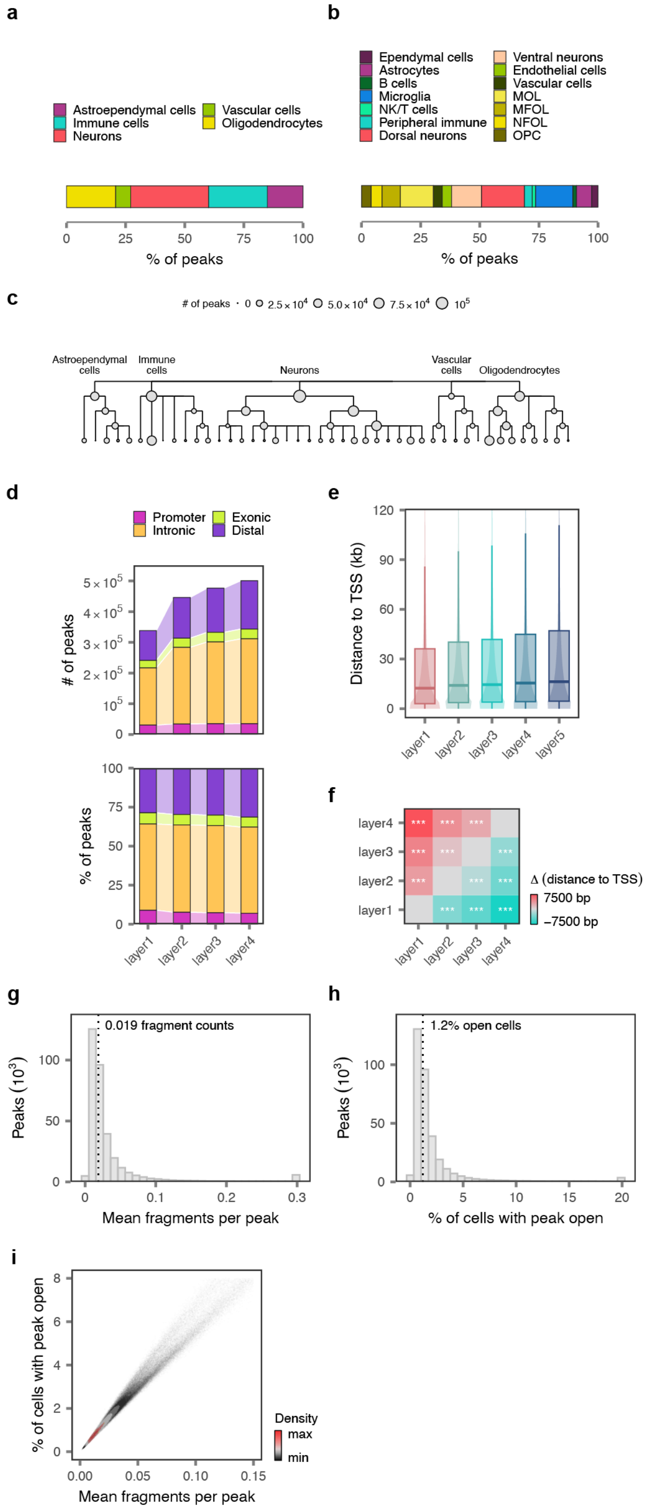


**Supplementary Fig. 23** | **Peak calling in the multiome atlas.**

**a,** Number of peaks attributed to each major (level 1) cell type by the iterative overlap-merging procedure in ArchR.

**b,** Number of peaks attributed to each level 2 cell type by the iterative overlap-merging procedure in ArchR.

**c,** Number of peaks attributed to each cell type throughout the cellular taxonomy of the spinal cord by the iterative overlap-merging procedure in ArchR.

**d,** Number, top, and proportion, bottom, of promoter, exonic, intronic, or distal peaks when calling peaks at levels 1 to 4 of the cellular taxonomy.

**e,** Distance to the nearest TSS for peaks called at levels 1 to 4 of the cellular taxonomy.

**f,** Statistical comparison of distance to the nearest TSS for peaks called at levels 1 to 4 of the cellular taxonomy, as shown in **e.** Calling peaks for cell subtypes defined at more granular resolutions leads to improved detection of distal regulatory elements.

**g,** Distribution of the mean fragment counts per cell, for all peaks called at level 4 of the cellular taxonomy.

**h,** Distribution of the mean proportion of cells in which each peak is accessible, for all peaks called at level 4 of the cellular taxonomy.

**i,** Scatterplot showing the relationship between the mean fragment counts per cell and the mean, for all peaks called at level 4 of the cellular taxonomy.

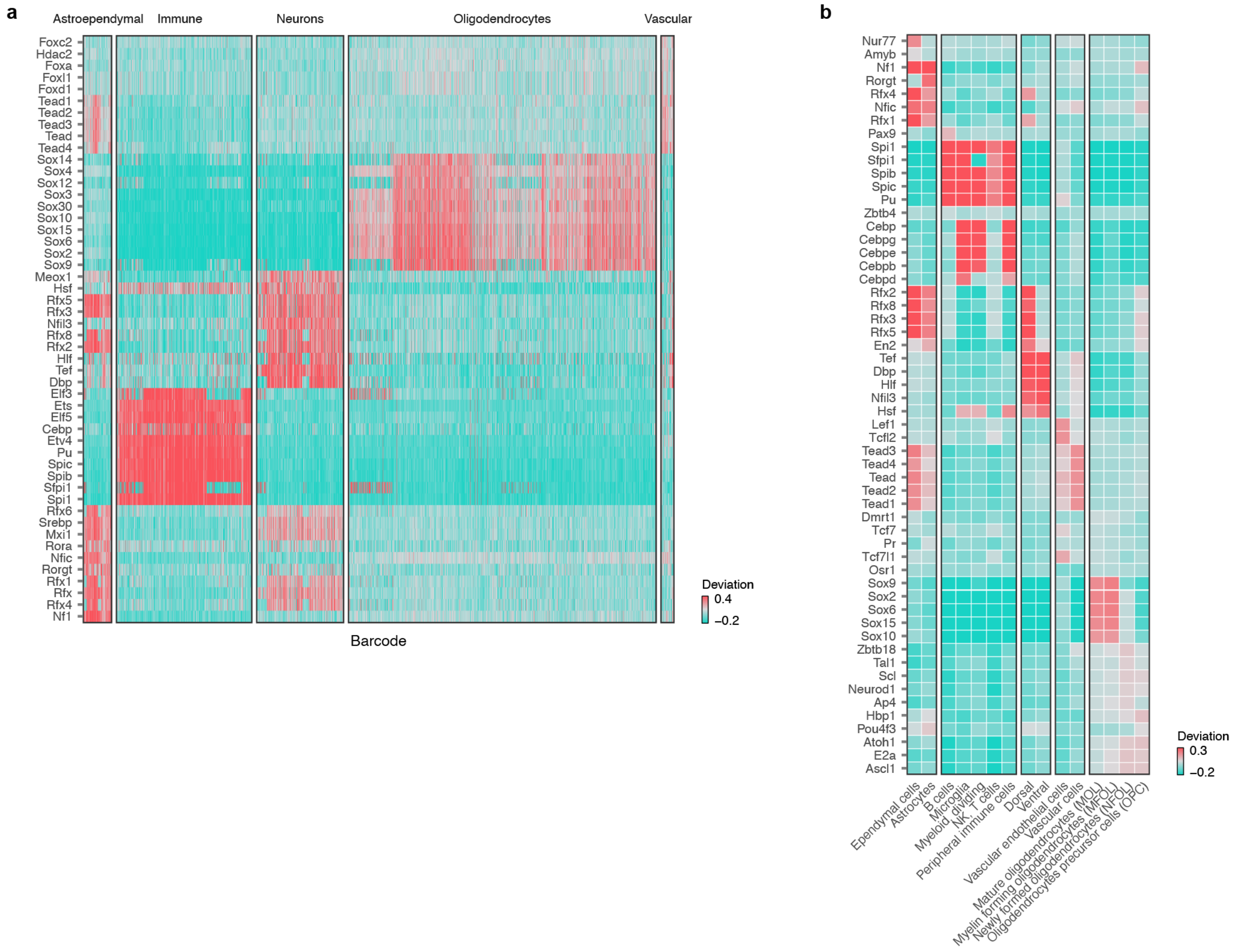


**Supplementary Fig. 24 | Cell-type-specific transcription factors in the multiome atlas.**

**a,** Deviations of the top ten cell-type-specific transcription factors for major (level 1) cell types identified by chromVAR, shown for each individual barcode in the multiome atlas.

**b,** Median deviations of the top five cell-type-specific transcription factors for level 2 cell types identified by chromVAR.

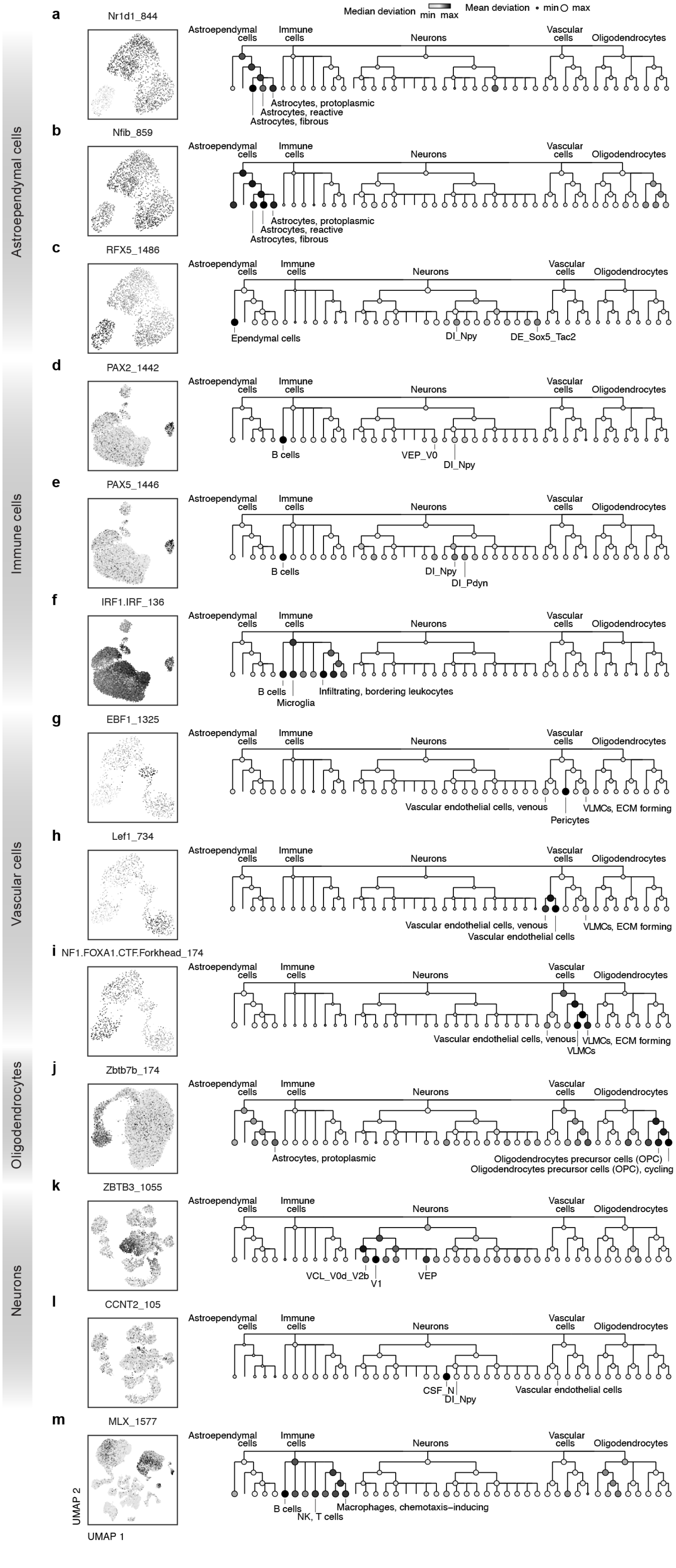

Median deviation min max
Mean deviation min max
Astroependymal cells
Immune cells
Vascular cells
Oligodendrocytes
Neurons
a
Nr1d1_844
Astrocytes, protoplasmic
Astrocytes, reactive
Astrocytes, fibrous
b
Nfib_859
Astrocytes, protoplasmic
Astrocytes, reactive
Astrocytes, fibrous
c
RFX5_1486
Ependymal cells
DI_Npy
DE_Sox5_Tac2
d
PAX2_1442
B cells
VEP_V0
DI_Npy
e
PAX5_1446
B cells
DI_Npy
DI_Pdyn
f
IRF1.IRF_136
B cells
Microglia
Infiltrating, bordering leukocytes
g
EBF1_1325
Vascular endothelial cells, venous
Pericytes
VLMCs, ECM forming
h
Lef1_734
Vascular endothelial cells, venous
Vascular endothelial cells
VLMCs, ECM forming
i
NF1.FOXA1.CTF.Forkhead_174
Vascular endothelial cells, venous
VLMCs
VLMCs, ECM forming
j
Zbtb7b_174
Astrocytes, protoplasmic
Oligodendrocytes precursor cells (OPC)
Oligodendrocytes precursor cells (OPC), cycling
k
ZBTB3_1055
VCL_V0d_V2b
V1
VEP
l
CCNT2_105
CSF_N
DI_Npy
Vascular endothelial cells
m
MLX_1577
B cells
NK, T cells
Macrophages, chemotaxis–inducing
UMAP 2
UMAP 1

**Supplementary Fig. 25 | Cell type- and subtype-specific transcription factors in the multiome atlas.**
Cell-type-specific transcription factors discussed in the main text. Left, UMAP visualizations of chromVAR motif deviations for cells subset to the relevant major (level 1) cell type, **a-l,** or for all cells in the dataset, **m.** Right, dendrograms showing mean and median chromVAR motif deviations per cell type for levels 1-4 of the clustering tree.

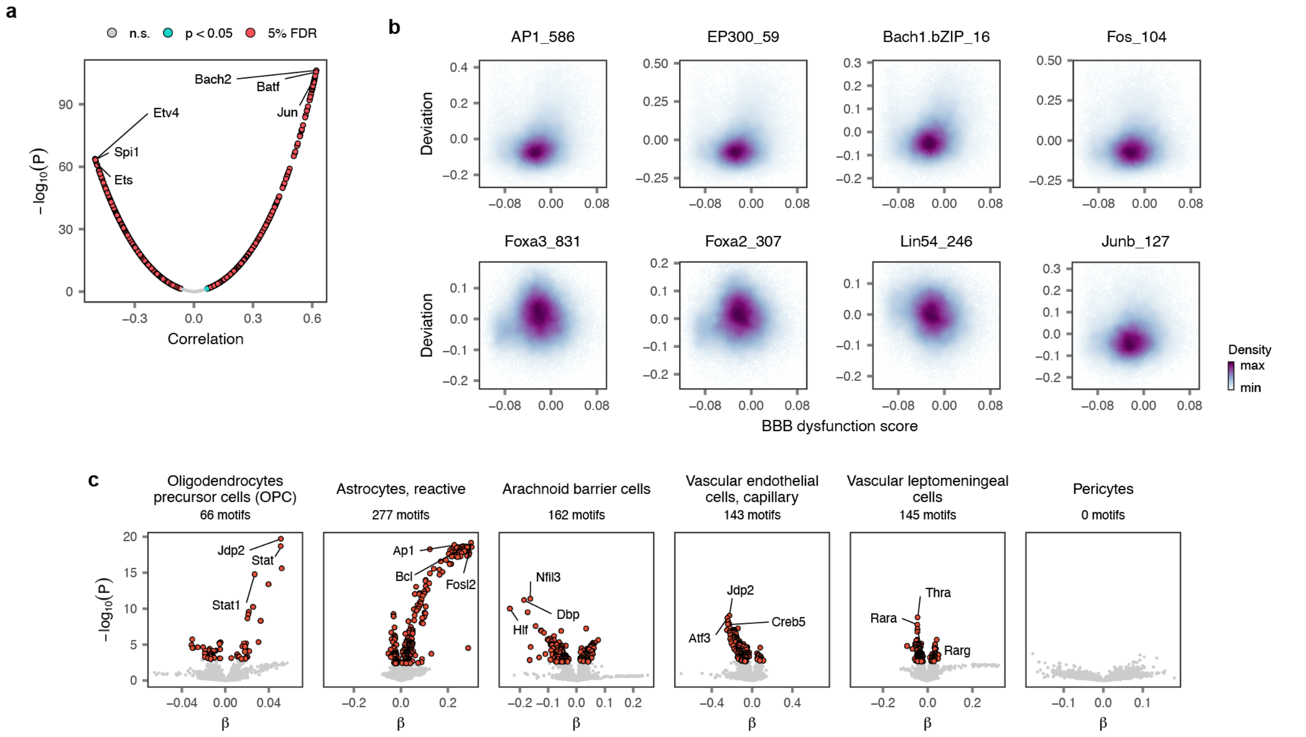


**Supplementary Fig. 26 | Regulatory programs underlying tripartite barrier formation.**

**a,** Volcano plot showing correlations between transcription factor binding motif accessibility, in the ATAC modality, and expression of the blood-brain barrier dysfunction score, in the RNA modality, across all vascular cells. The top three correlated and anti-correlated motifs are annotated.

**b,** Scatterplots highlighting relationships between expression of the blood-brain barrier dysfunction score and transcription factor binding motif accessibility for selected motifs discussed in the main text. Points are colored by the density of individual nuclei.

**c,** Volcano plots showing differentially accessible transcription factor binding motifs in the other cell types that establish the tripartite barrier.

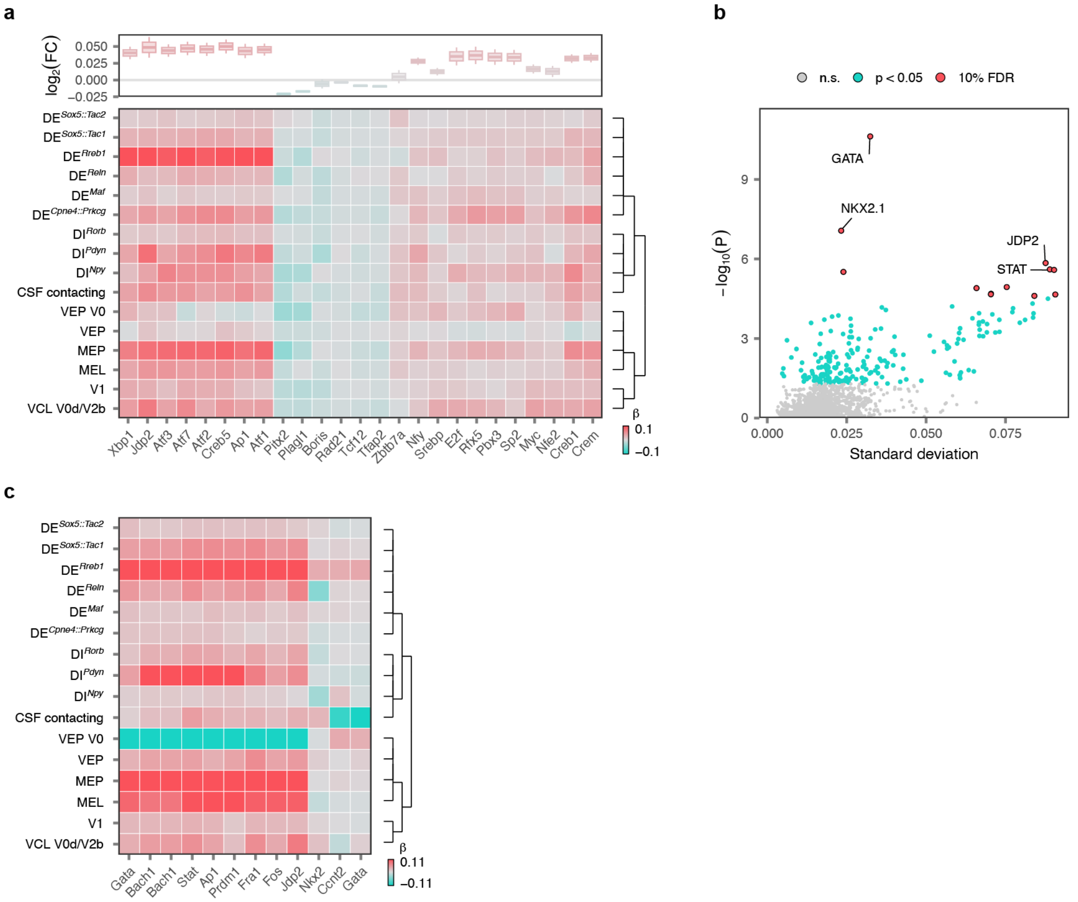


**Supplementary Fig. 27 | Conserved and divergent regulatory programs in injured neurons.**

**a,** Top 25 transcription factor binding motifs with conserved patterns of up- or downregulation across level 4 neuron subtypes at 7 days post-injury. Top, boxplot showing the estimated effect of SCI on transcription factor binding motif accessibility across all level 4 neuron subtypes. Bottom, heatmap showing the estimated effect of SCI on transcription factor binding motif accessibility within each level 4 neuron subtype individually.

**b,** Identification of transcription factor binding motifs with significant variation across neuron subtypes in their patterns of accessibility at 2 months post-injury. The x-axis shows the standard deviation of coefficients estimated by linear mixed models for each neuron subtype independently, while the y-axis shows the statistical significance of the standard deviation compared to linear mixed models fit with randomized neuron subtype assignments.

**c,** Heatmap showing the estimated effect of SCI on transcription factor binding motif accessibility across level 4 neuron subtypes for the significantly variable motif shown in **b.**

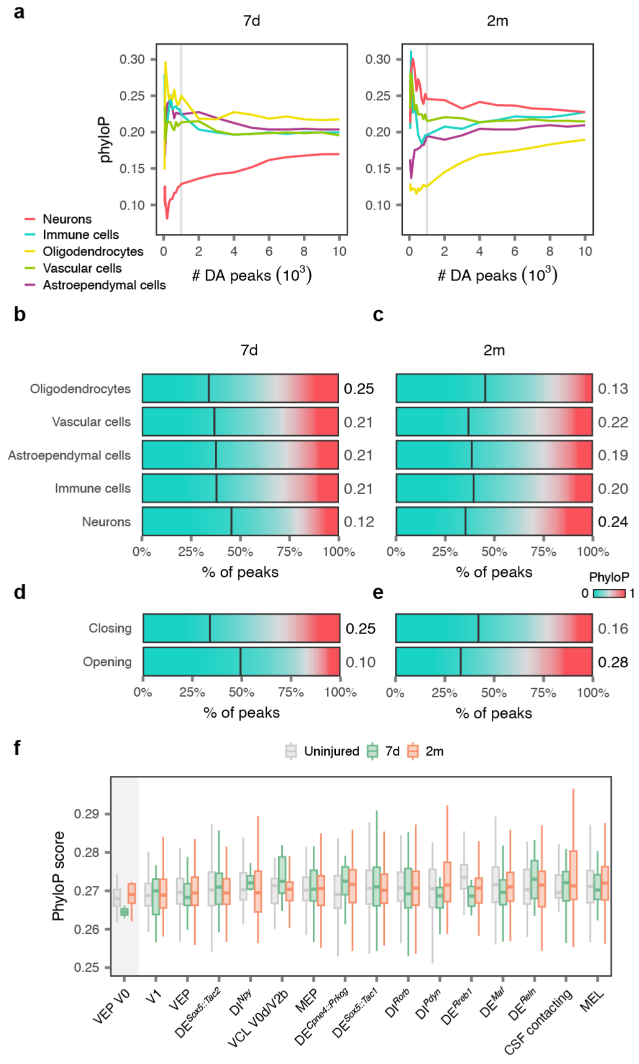


**Supplementary Fig. 28 | Evolutionary conservation and acceleration of differentially accessible regions.**

**a,** Line charts showing the median phyloP score of the top-*n* differentially accessible peaks, for values of *n* between 50 and 10,000, within major cell types of the spinal cord at 7 days and 2 months after SCI. Vertical lines highlight the top-1,000 differentially accessible peaks within each cell type.

**b-c,** Mean phyloP scores over all nucleotides within each of the top 1,000 differentially accessible peaks in each major cell type at 7 days, **b,** and 2 months, **c,** after SCI. Vertical line shows the median phyloP score for all peaks in the dataset.

**d-e,** Mean phyloP scores over all nucleotides within each of the top 1,000 peaks that are opening versus closing in neurons at 7 days, **d,** and 2 months, **e,** after SCI. Vertical line shows the median phyloP score for all peaks in the dataset.

**f,** Mean phyloP score over all accessible peaks in nuclei from each level 4 neuron subtype in the multiome atlas, split by experimental condition. Ventral excitatory projection neurons are highlighted.

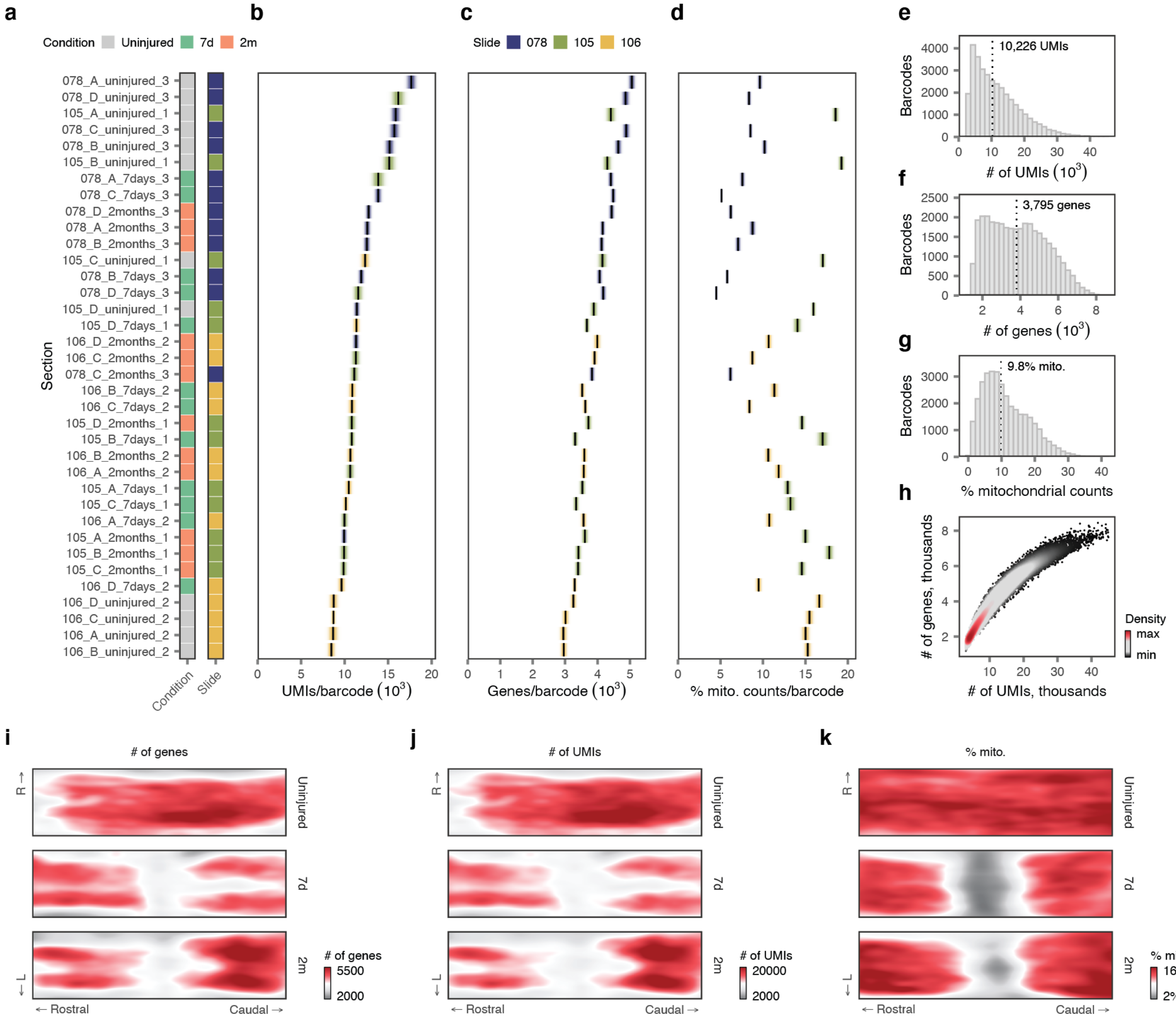


**Supplementary Fig. 29 | Quality control of the 2D spatial atlas.**

**a-d,** Quality control statistics for 36 sections profiled by spatial transcriptomics (**a,** legend showing the experimental condition and slide number for each section; **b,** mean number of UMIs per barcode; **c,** mean number of genes detected per barcode; **d,** mean proportion of mitochondrial counts per barcode). Shaded areas show the standard deviation. Sections are colored by the slide on which each section was sequenced.

**e-g,** Aggregate distributions of quality control statistics over all sections (**e,** number of UMIs per barcode; **f,** number of genes detected per barcode; **g,** proportion of mitochondrial counts per barcode).

**h,** Relationship between the number of UMIs and number of genes detected per spatial barcode.

**i-k,** Quality control statistics visualized on the common coordinate system of the spinal cord, separately for each experimental condition (**i,** mean number of UMIs per barcode; **j,** mean number of genes detected per barcode; **k,** mean proportion of mitochondrial counts per barcode).

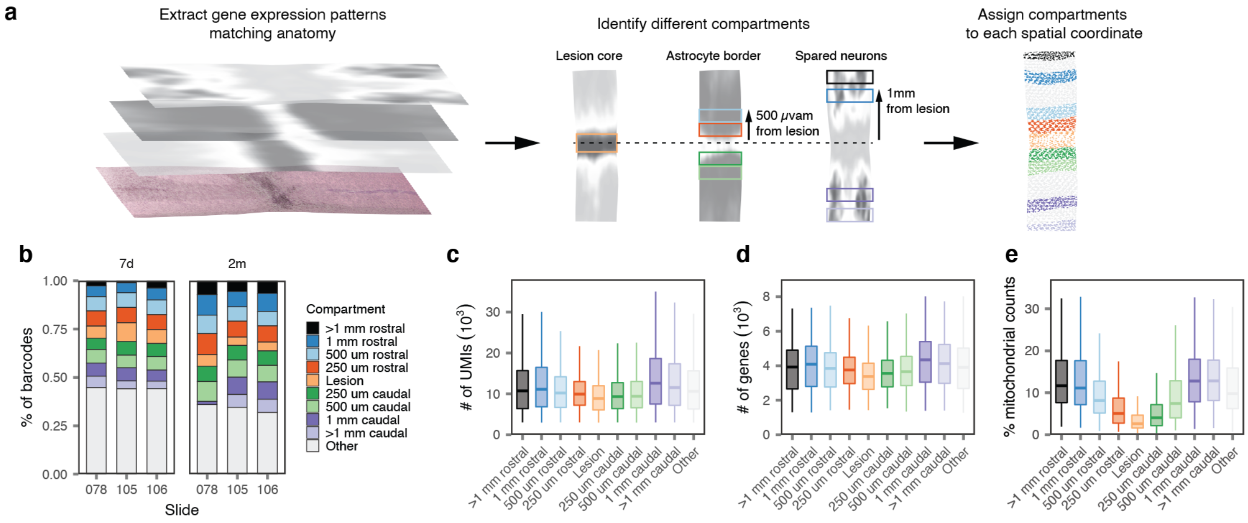


**Supplementary Fig. 30** | **Definition and quality control of lesion compartments.**

**a,** Schematic overview of the procedure for definition of lesion compartments on the common coordinate system of the spinal cord.

**b,** Proportion of spatial barcodes assigned to each lesion compartment within each slide.

**c-e,** Distributions of quality control statistics (**c,** number of UMIs per barcode; **d,** number of genes detected per barcode; **e,** proportion of mitochondrial counts per barcode) for spatial barcodes within each lesion compartment.

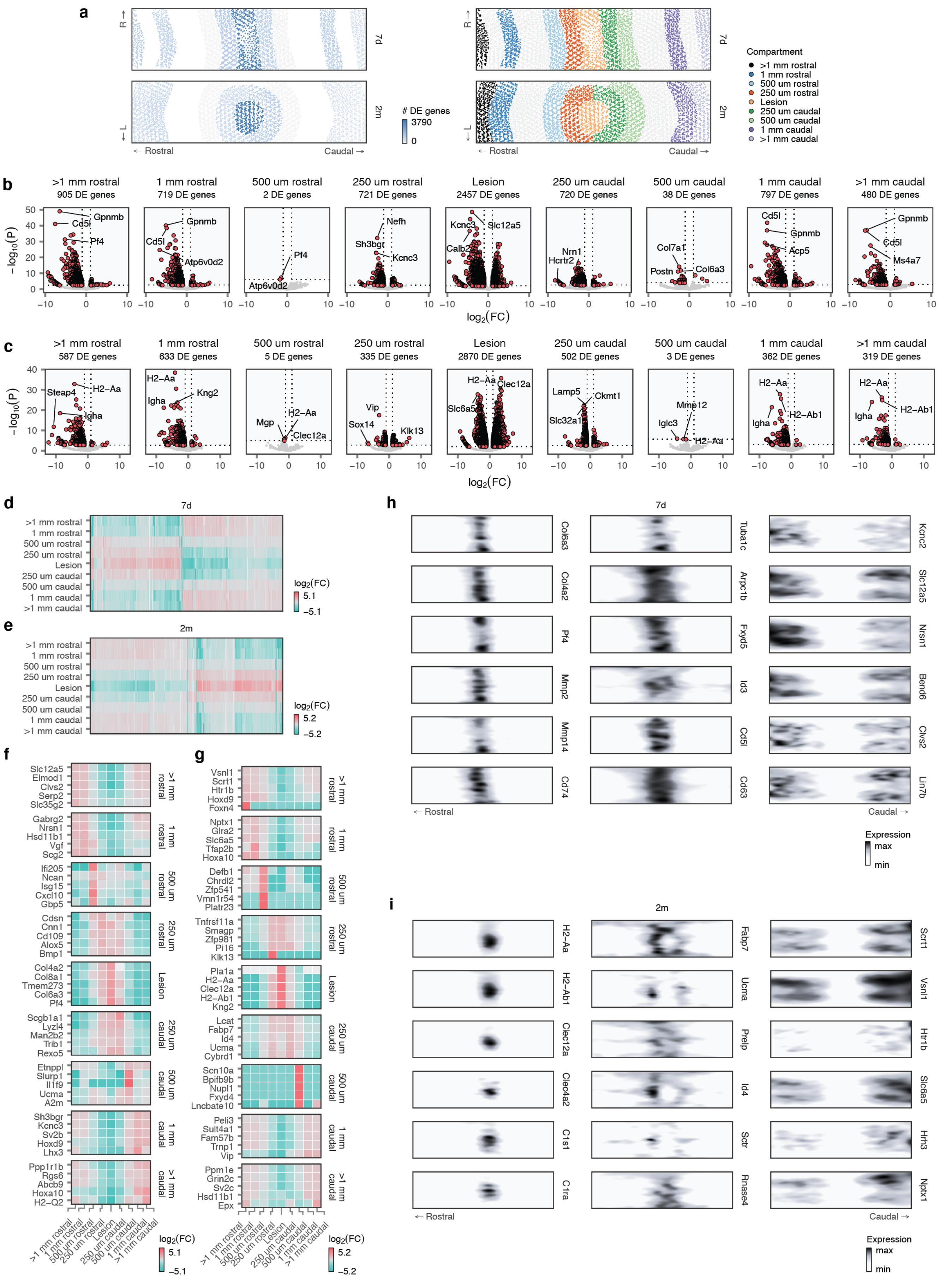
a
R →
← L
7d
2m
← Rostral
Caudal →
### DE genes
3790
0
Compartment
>1 mm rostral
1 mm rostral
500 um rostral
250 um rostral
Lesion
250 um caudal
500 um caudal
1 mm caudal
>1 mm caudal
b
>1 mm rostral 905 DE genes
1 mm rostral 719 DE genes
500 um rostral 2 DE genes
250 um rostral 721 DE genes
Lesion 2457 DE genes
250 um caudal 720 DE genes
500 um caudal 38 DE genes
1 mm caudal 797 DE genes
>1 mm caudal 480 DE genes
−log10(P)
log2(FC)
c
>1 mm rostral 587 DE genes
1 mm rostral 633 DE genes
500 um rostral 5 DE genes
250 um rostral 335 DE genes
Lesion 2870 DE genes
250 um caudal 502 DE genes
500 um caudal 3 DE genes
1 mm caudal 362 DE genes
>1 mm caudal 319 DE genes
−log10(P)
log2(FC)
d
7d
log2(FC)
5.1
−5.1
e
2m
log2(FC)
5.2
−5.2
f
g
h
7d
← Rostral
Caudal →
Expression
max
min
i
2m
← Rostral
Caudal →
Expression
max
min

**Supplementary Fig. 31 | Shared and distinct programs of gene expression across lesion compartments.**
**a,** Left, total number of differentially expressed genes detected within each lesion compartment at 7 days and 2 months after SCI. Right, legend showing the position of each lesion compartment, as in **Fig. 6c.**
**b-c,** Volcano plots showing differentially expressed genes for all lesion compartments at 7 days, **b,** and 2 months, **c.** The top three genes per lesion compartment, as ranked by the product of the $\log_2$-fold change and the $-\log_{10}$ p-value, are annotated.
**d-e,** Heatmap showing $\log_2$-fold changes for all genes differentially expressed in at least one lesion compartment at 7 days, **d,** and 2 months, **e.**
**f-g,** Heatmap showing $\log_2$-fold changes for the top five genes differentially expressed in each lesion compartment at 7 days, **f,** and 2 months, **g.**
**h-i,** Visualization of selected differentially expressed genes specific to individual lesion compartments at 7 days, **h,** and 2 months, **i,** on the two-dimensional coordinate system of the injured spinal cord.

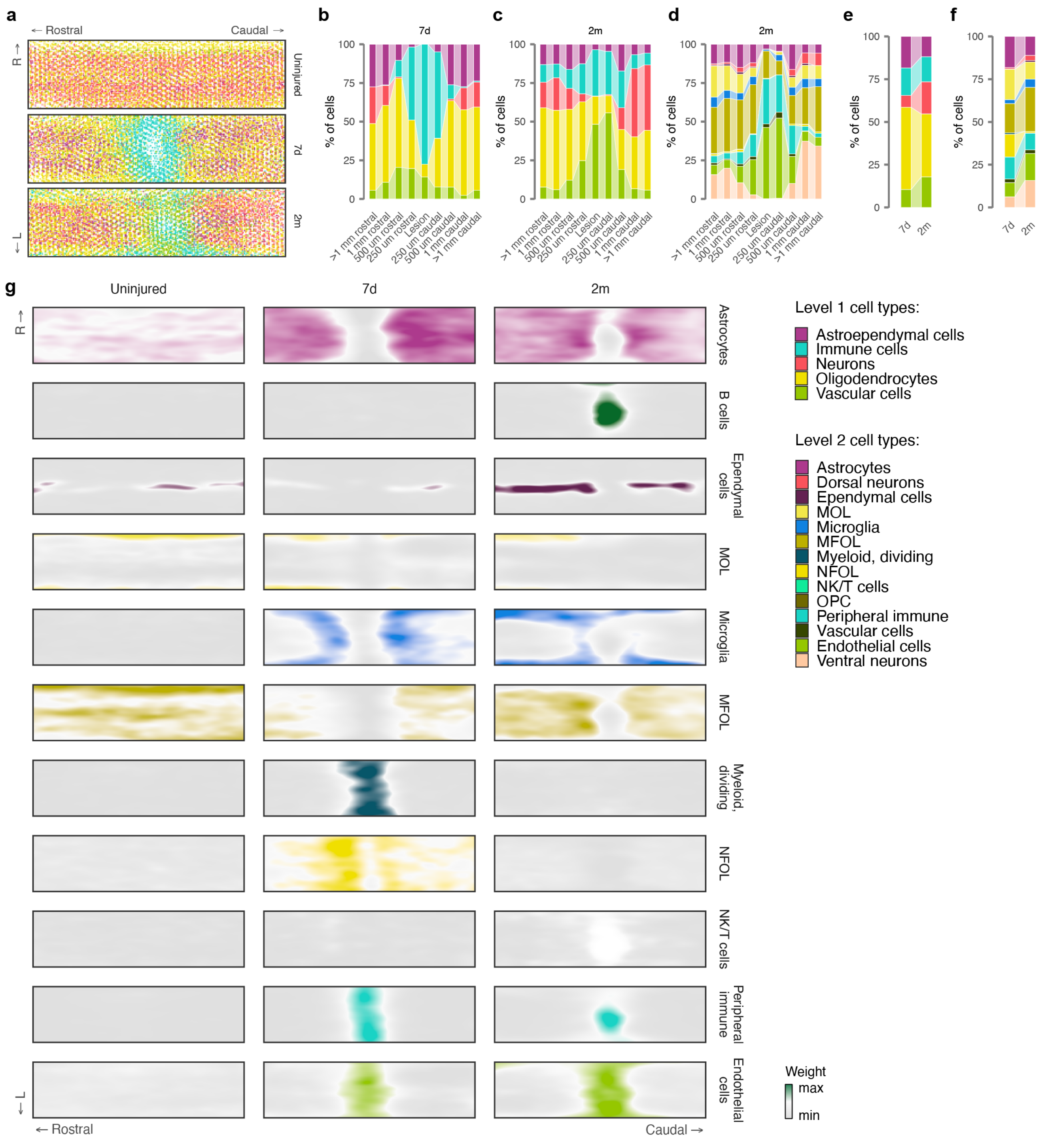


**Supplementary Fig. 32 | Cell type deconvolution of the 2D spatial atlas.**

**a,** Major cell types assigned to each spatial barcode, visualized for each experimental condition on the two-dimensional coordinate system of the injured spinal cord.

**b,** Sankey diagram showing the cellular composition of each lesion compartment at 7 days, for major (level 1) cell types.

**c,** Sankey diagram showing the cellular composition of each lesion compartment at 2 months, for level 1 cell types.

**d,** Sankey diagram showing the cellular composition of each lesion compartment at 2 months, for level 2 cell types.

**e,** Sankey diagram showing the evolution of the cellular composition of the entire injured spinal cord between 7 days and 2 months, for level 1 cell types.

**f,** Sankey diagram showing the evolution of the cellular composition of the entire injured spinal cord between 7 days and 2 months, for level 2 cell types.

**g,** Visualization of the deconvolution weights assigned by RCTD for selected level 2 cell types at each timepoint, on the two-dimensional coordinate system of the injured spinal cord.

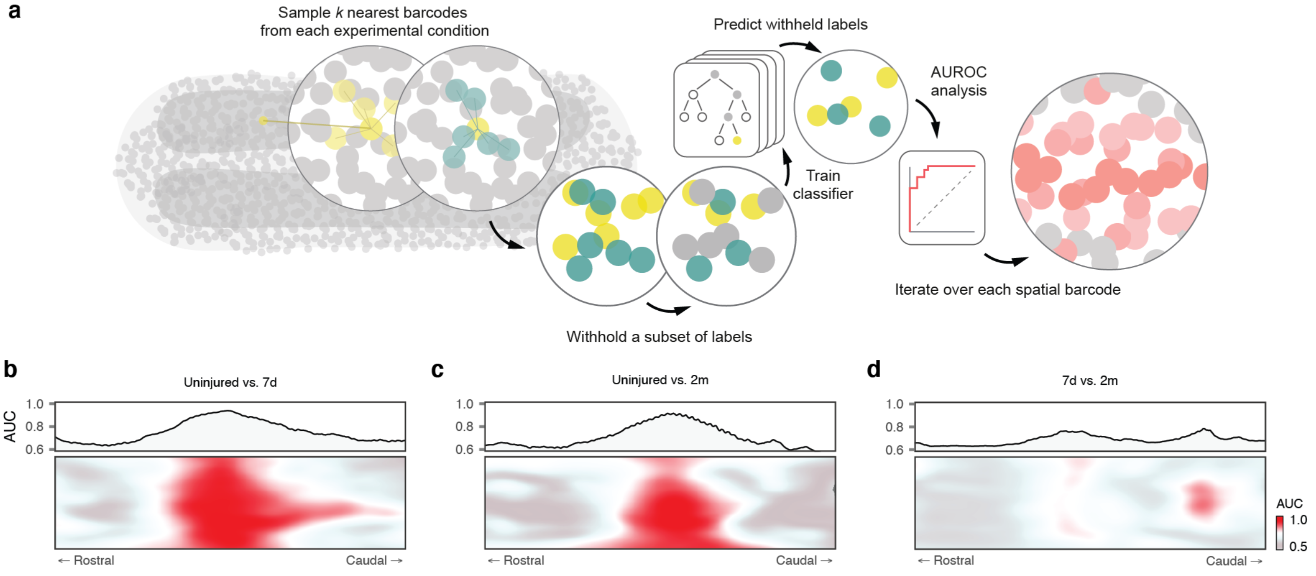


**Supplementary Fig. 33** | **Spatial prioritization of the 2D spatial atlas.**
**a,** Schematic overview illustrating the concept of spatial prioritization, as captured in Magellan.
**b-d,** Spatial prioritizations assigned by Magellan to each pairwise comparison of experimental conditions in the 2D spatial atlas (**b,** 7 days versus uninjured; **c,** 2 months versus uninjured, **d,** 7 days versus 2 months). Bottom, AUCs of spatial prioritization visualized on the two-dimensional coordinate system of the injured spinal cord. Top, average AUCs of spatial prioritization along the rostrocaudal axis, highlighting prioritization of the lesion site.

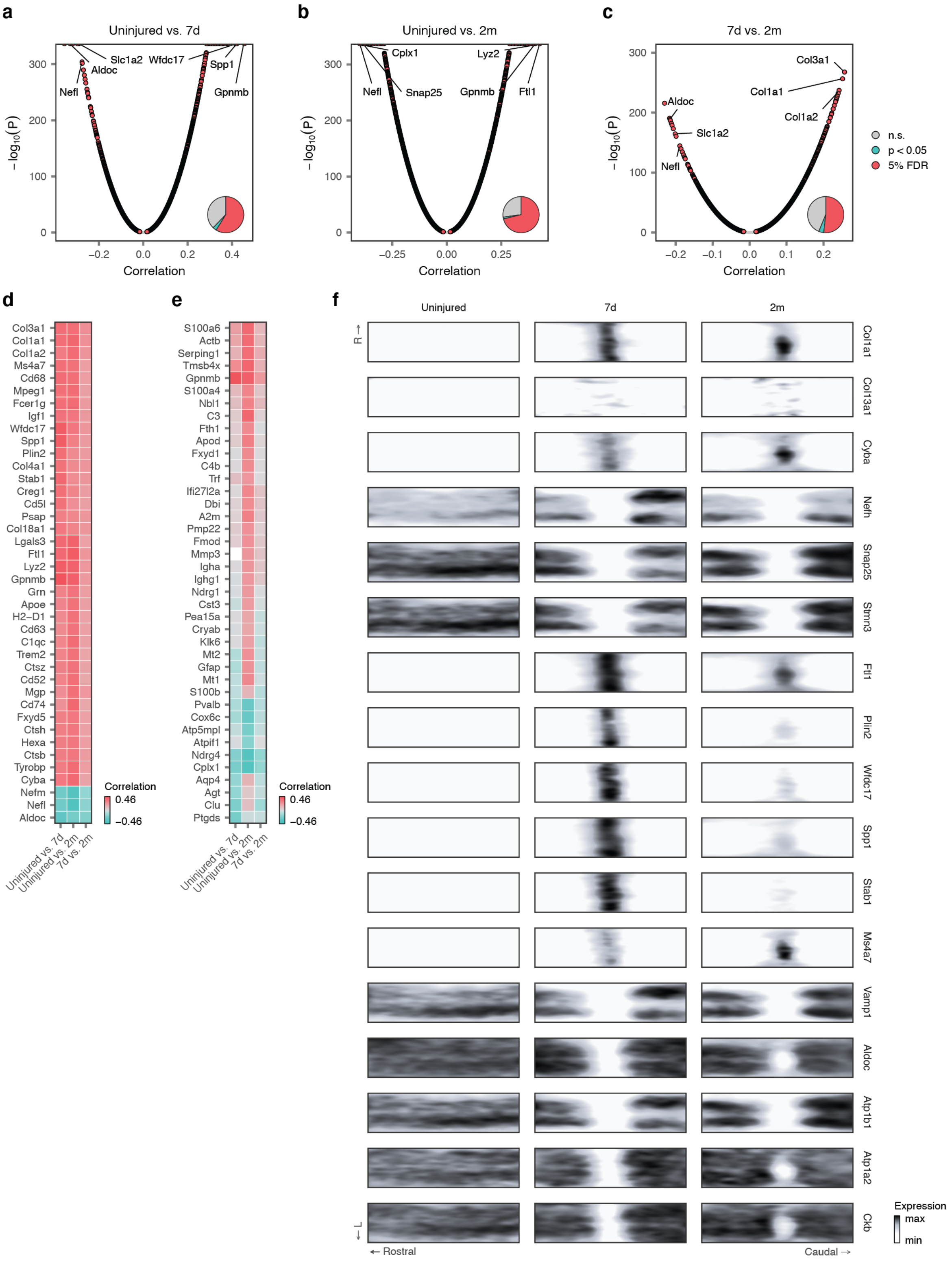


**Supplementary Fig. 34 | Molecular basis of spatial prioritization at the gene level.**
**a-c,** Volcano plots showing correlations between the AUCs assigned by Magellan at each spatial barcode and transcriptome-wide gene expression across the same spatial barcodes (**a,** 7 days versus uninjured; **b,** 2 months versus uninjured; **c,** 7 days versus 2 months). Inset pie charts show the proportions of all tested genes that are significantly correlated with the spatial prioritizations.
**d,** Heatmap showing Pearson correlations between spatial prioritizations and gene expression for each pairwise comparison of experimental conditions, for the top 40 most recurrently correlated genes across all comparisons.
**e,** Heatmap showing Pearson correlations between spatial prioritizations and gene expression for each pairwise comparison of experimental conditions, for the top 40 most variably correlated genes across all comparisons.
**f,** Visualization of selected genes prioritized by their correlation to spatial prioritizations on the two-dimensional coordinate system of the injured spinal cord.

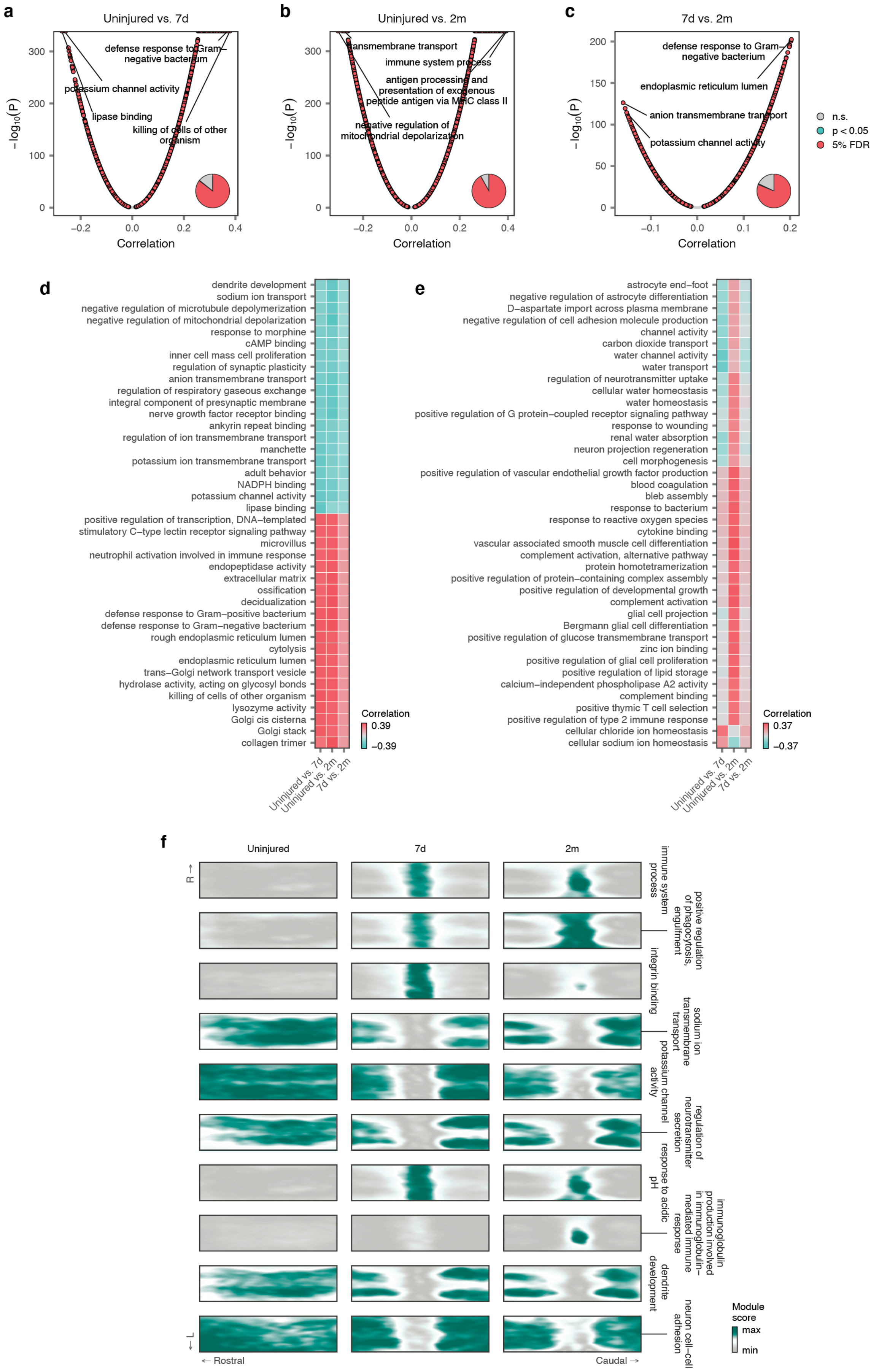
a
Uninjured vs. 7d
defense response to Gram−negative bacterium
potassium channel activity
lipase binding
killing of cells of other organism
b
Uninjured vs. 2m
transmembrane transport
immune system process
antigen processing and presentation of exogenous peptide antigen via MHC class II
negative regulation of mitochondrial depolarization
c
7d vs. 2m
defense response to Gram−negative bacterium
endoplasmic reticulum lumen
anion transmembrane transport
potassium channel activity
−log10(P)
Correlation
n.s.
p < 0.05
5% FDR
d
dendrite development
sodium ion transport
negative regulation of microtubule depolymerization
negative regulation of mitochondrial depolarization
response to morphine
cAMP binding
inner cell mass cell proliferation
regulation of synaptic plasticity
anion transmembrane transport
regulation of respiratory gaseous exchange
integral component of presynaptic membrane
nerve growth factor receptor binding
ankyrin repeat binding
regulation of ion transmembrane transport
manchette
potassium ion transmembrane transport
adult behavior
NADPH binding
potassium channel activity
lipase binding
positive regulation of transcription, DNA−templated
stimulatory C−type lectin receptor signaling pathway
microvillus
neutrophil activation involved in immune response
endopeptidase activity
extracellular matrix
ossification
decidualization
defense response to Gram−positive bacterium
defense response to Gram−negative bacterium
rough endoplasmic reticulum lumen
cytolysis
endoplasmic reticulum lumen
trans−Golgi network transport vesicle
hydrolase activity, acting on glycosyl bonds
killing of cells of other organism
lysozyme activity
Golgi cis cisterna
Golgi stack
collagen trimer
Correlation
0.39
−0.39
Uninjured vs. 7d
Uninjured vs. 2m
7d vs. 2m
e
astrocyte end−foot
negative regulation of astrocyte differentiation
D−aspartate import across plasma membrane
negative regulation of cell adhesion molecule production
channel activity
carbon dioxide transport
water channel activity
water transport
regulation of neurotransmitter uptake
cellular water homeostasis
water homeostasis
positive regulation of G protein−coupled receptor signaling pathway
response to wounding
renal water absorption
neuron projection regeneration
cell morphogenesis
positive regulation of vascular endothelial growth factor production
blood coagulation
bleb assembly
response to bacterium
response to reactive oxygen species
cytokine binding
vascular associated smooth muscle cell differentiation
complement activation, alternative pathway
protein homotetramerization
positive regulation of protein−containing complex assembly
positive regulation of developmental growth
complement activation
glial cell projection
Bergmann glial cell differentiation
positive regulation of glucose transmembrane transport
zinc ion binding
positive regulation of glial cell proliferation
positive regulation of lipid storage
calcium−independent phospholipase A2 activity
complement binding
positive thymic T cell selection
positive regulation of type 2 immune response
cellular chloride ion homeostasis
cellular sodium ion homeostasis
Correlation
0.37
−0.37
Uninjured vs. 7d
Uninjured vs. 2m
7d vs. 2m
f
Uninjured
7d
2m
R →
← L
← Rostral
Caudal →
immune system process
positive regulation of phagocytosis, engulfment
integrin binding
sodium ion transmembrane transport
potassium channel activity
regulation of neurotransmitter secretion
response to acidic pH
immunoglobulin production involved in immunoglobulin−mediated immune response
dendrite development
neuron cell−cell adhesion
Module score
max
min

**Supplementary Fig. 35 | Molecular basis of spatial prioritization at the gene module level.**
**a-c,** Volcano plots showing correlations between the AUCs assigned by Magellan at each spatial barcode and the average expression of genes associated with a given GO term ("GO modules") across the same spatial barcodes (**a,** 7 days versus uninjured; **b,** 2 months versus uninjured; **c,** 7 days versus 2 months). Inset pie charts show the proportions of all tested GO modules that are significantly correlated with the spatial prioritizations.
**d,** Heatmap showing Pearson correlations between spatial prioritizations and GO module scores for each pairwise comparison of experimental conditions, for the top 40 most recurrently correlated GO modules across all comparisons.
**e,** Heatmap showing Pearson correlations between spatial prioritizations and GO module scores for each pairwise comparison of experimental conditions, for the top 40 most variably correlated GO modules across all comparisons.
**f,** Visualization of selected GO modules prioritized by their correlation to spatial prioritizations on the two-dimensional coordinate system of the injured spinal cord.

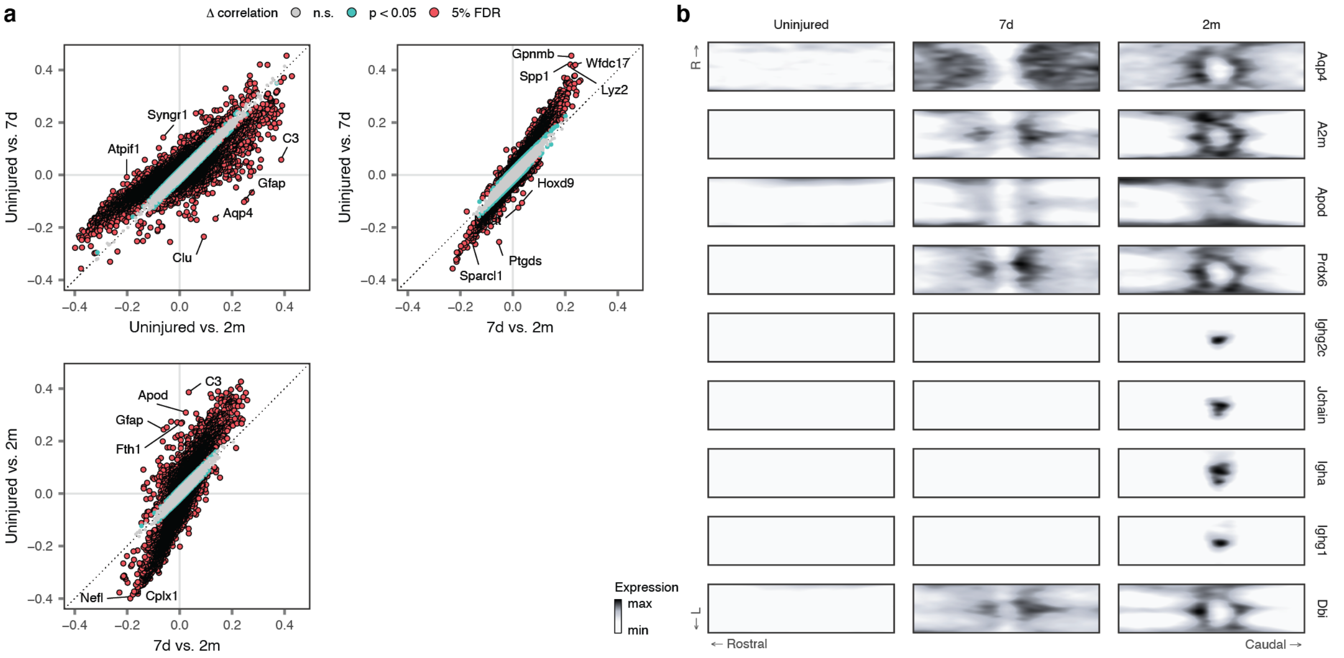


**Supplementary Fig. 36 | Molecular basis of differential spatial prioritization at the gene level.**

**a,** Scatterplots highlighting genes with differential correlations to spatial prioritizations between pairwise comparisons of experimental conditions.

**b,** Visualization of selected genes prioritized by their differential correlation to spatial prioritizations between comparisons on the two-dimensional coordinate system of the injured spinal cord.

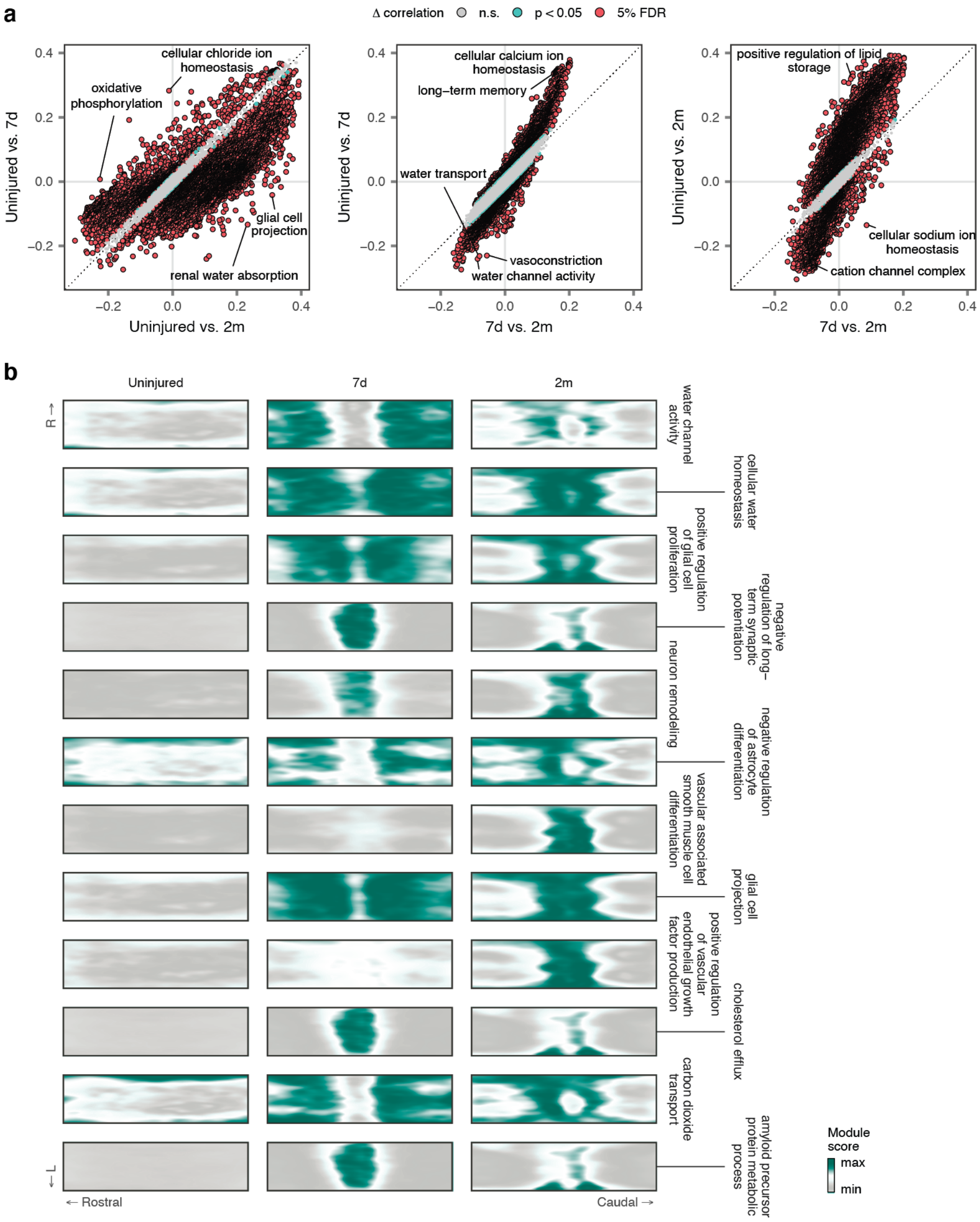


**Supplementary Fig. 37 | Molecular basis of differential spatial prioritization at the gene module level.**
**a,** Scatterplots highlighting GO modules with differential correlations to spatial prioritizations between pairwise comparisons of experimental conditions. **b,** Visualization of selected GO modules prioritized by their differential correlation to spatial prioritizations between comparisons on the two-dimensional coordinate system of the injured spinal cord.

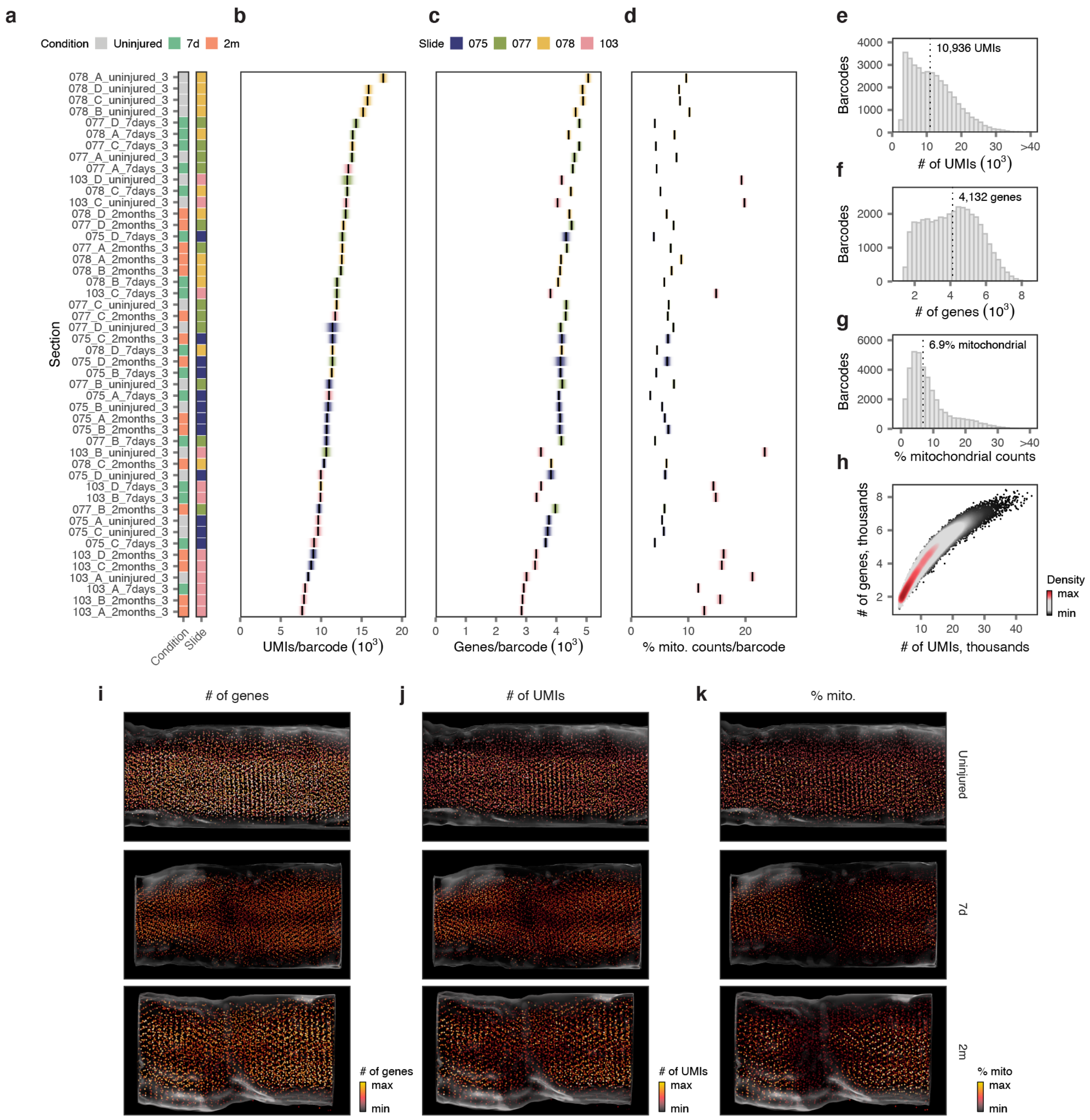

**Supplementary Fig. 38 | Quality control of the 3D spatial atlas.**

**a-d,** Quality control statistics for 16 sections profiled by three-dimensional spatial transcriptomics (**a,** legend showing the experimental condition and slide number for each section; **b,** mean number of UMIs per barcode; **c,** mean number of genes detected per barcode; **d,** mean proportion of mitochondrial counts per barcode). Shaded areas show the standard deviation. Sections are colored by the slide on which each section was sequenced.

**e-g,** Aggregate distributions of quality control statistics over all sections in the 3D spatial atlas (**e,** number of UMIs per barcode; **f,** number of genes detected per barcode; **g,** proportion of mitochondrial counts per barcode).

**h,** Relationship between the number of UMIs and number of genes detected per spatial barcode in the 3D spatial atlas.

**i-k,** Quality control statistics visualized on the three-dimensional common coordinate system of the spinal cord, separately for each experimental condition (**i,** mean number of UMIs per barcode; **j,** mean number of genes detected per barcode; **k,** mean proportion of mitochondrial counts per barcode).

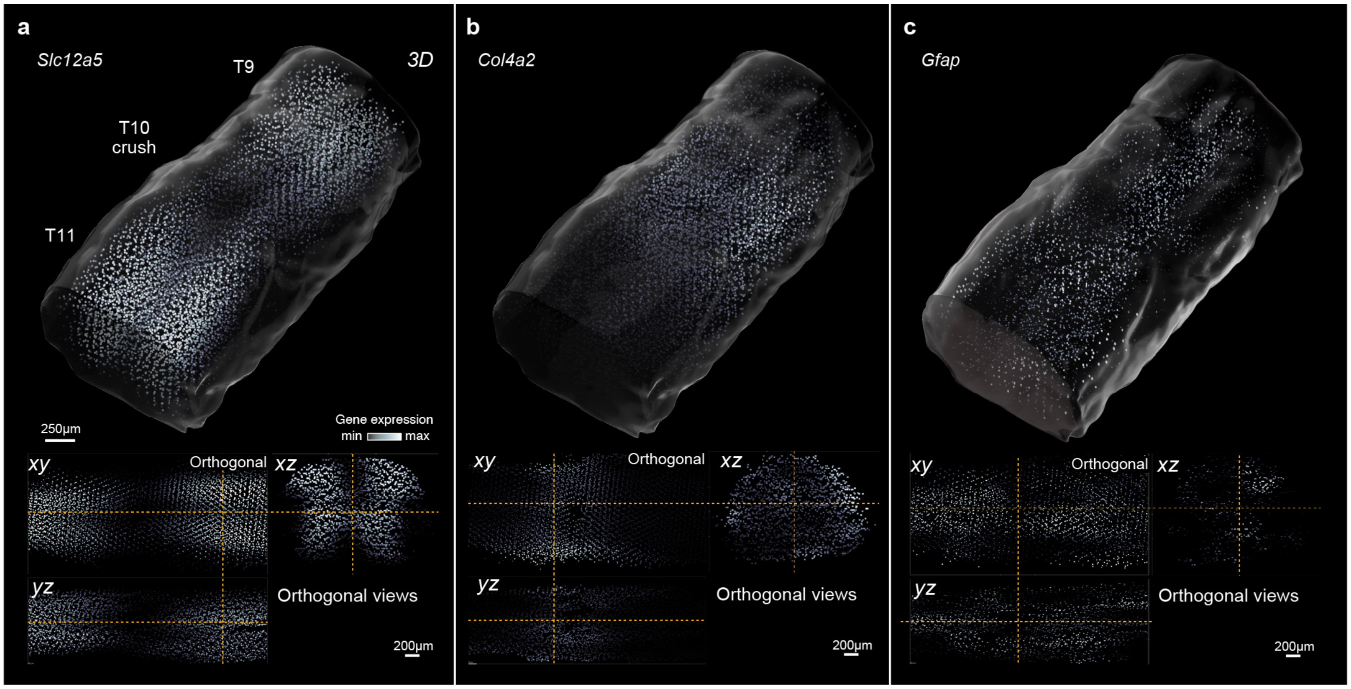


**Supplementary Fig. 39 | Lesion compartments in the 3D spatial atlas.**
Expression of marker genes associated with distinct lesion compartments across the 3D spatial transcriptomic atlas at 7 days, including spared but reactive neural tissue (*Slc12a5*, **a**) the fibrotic scar (*Col4a2*, **b**), and the astrocyte barrier (*Gfap*, **c**), as shown in **Fig. 8b** but here visualized separately for each gene.

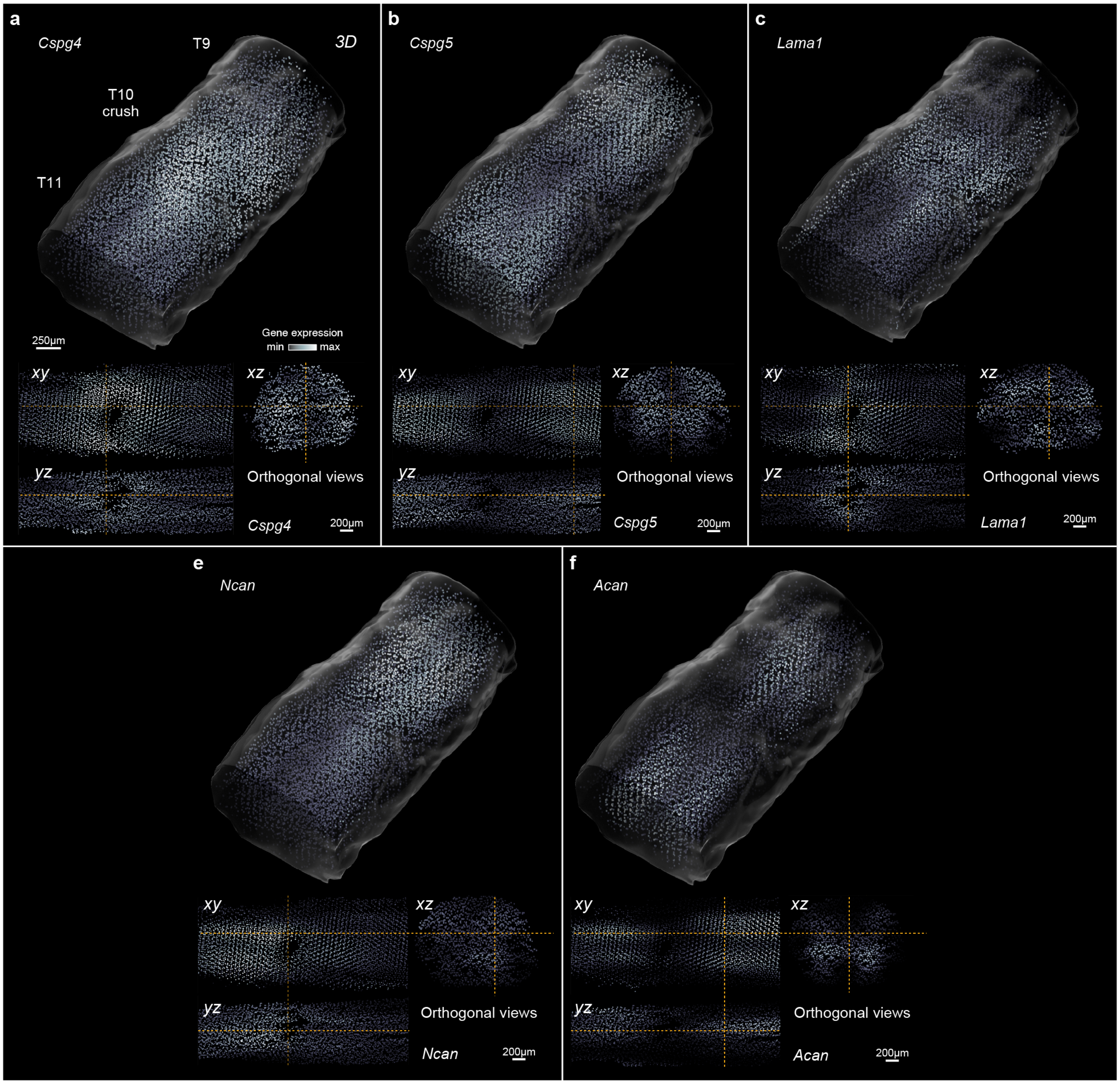


**Supplementary Fig. 40 | Inhibitory and facilitating molecules in the 3D spatial atlas.**
Expression of selected inhibitory and facilitating molecules across the 3D spatial transcriptomic atlas at 7 days. **a,** *Cspg4*; **b,** *Cspg5*; **c,** *Ncan*; **d,** *Acan*; **e,** *Lama1*.

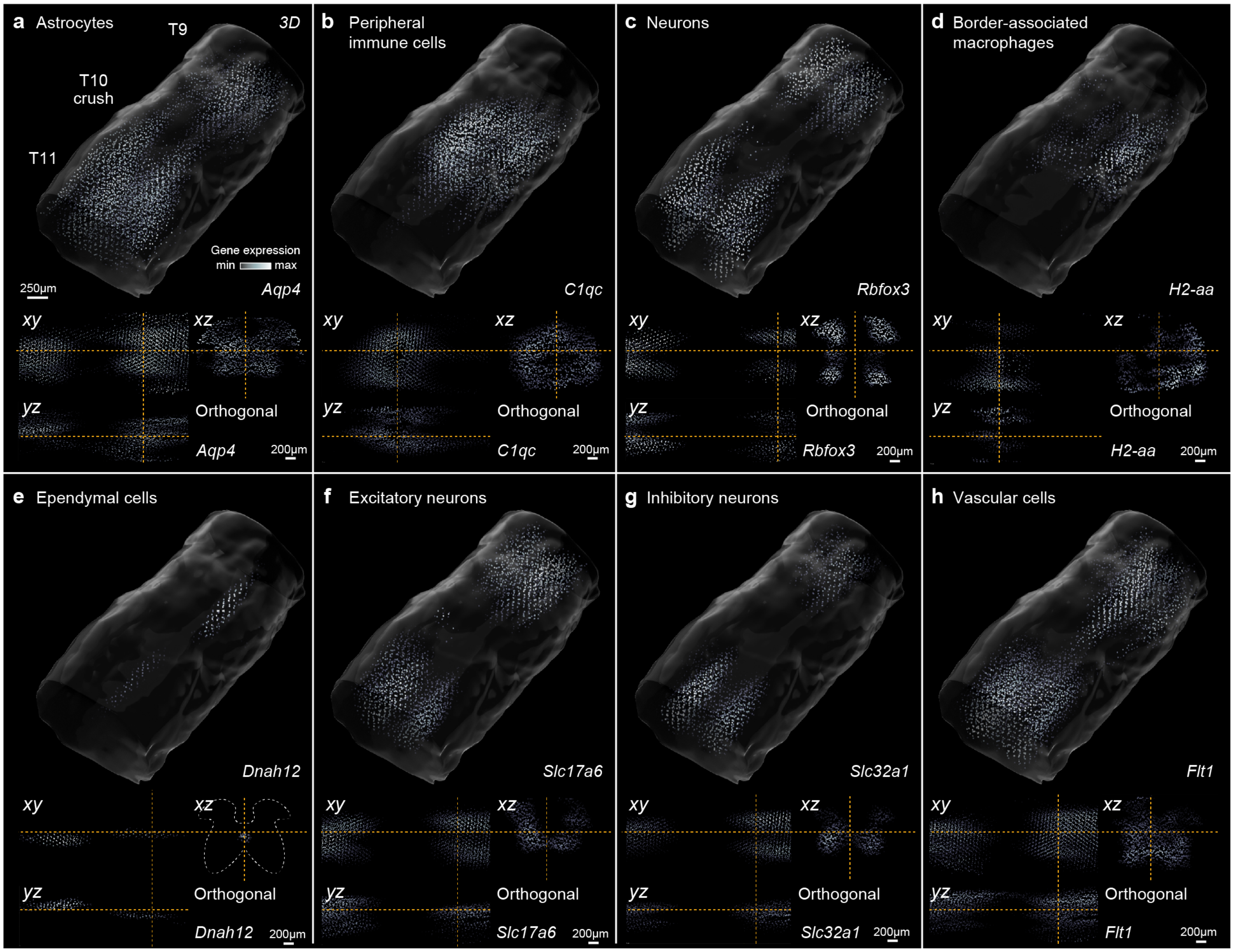


**Supplementary Fig. 41 | Cell type marker gene expression in the 3D spatial atlas.**
Expression of selected cell type marker genes across the 3D spatial transcriptomic atlas at 7 days. **a,** *Aqp4* (astrocytes); **b,** *C1qc* (peripheral immune cells); **c,** *Rbfox3* (neurons); **d,** *H2-aa* (border-associated macrophages); **e,** *Dnah12* (ependymal cells); **f,** *Slc17a6*, excitatory neurons; **g,** *Slc32a1* (inhibitory neurons); **h,** *Flt1* (vascular cells).

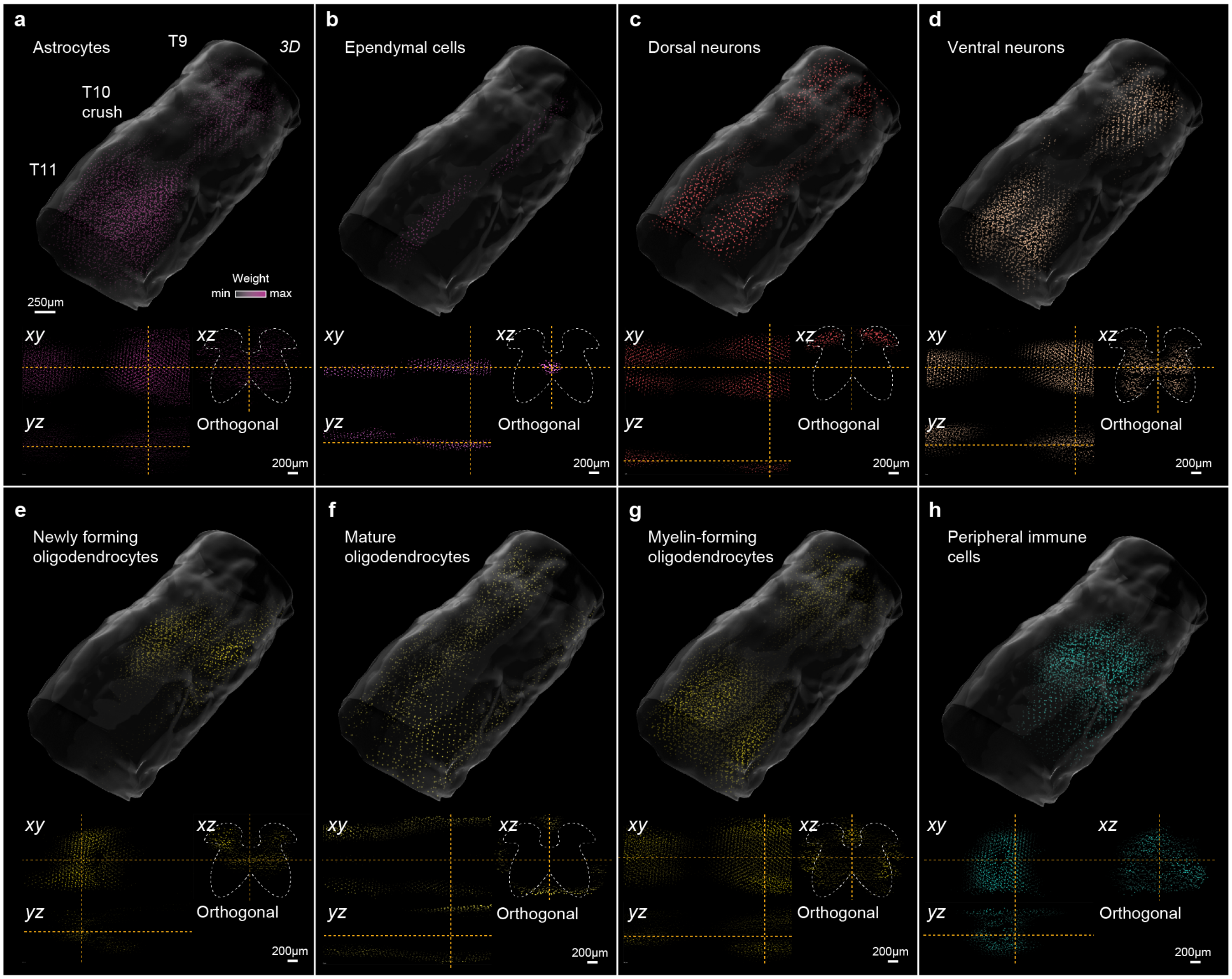


**Supplementary Fig. 42 | Cell type deconvolution of the 3D spatial atlas.**
Visualization of the deconvolution weights assigned by RCTD for selected level 2 cell types at 7 days, on the two-dimensional coordinate system of the injured spinal cord. **a,** Astrocytes; **b,** ependymal cells; **c,** dorsal neurons; **d,** ventral neurons; **e,** newly forming oligodendrocytes (NFOL); **f,** mature oligodendrocytes (MOL); **g,** myelin-forming oligodendrocytes (MFOL); **h,** peripheral immune cells.

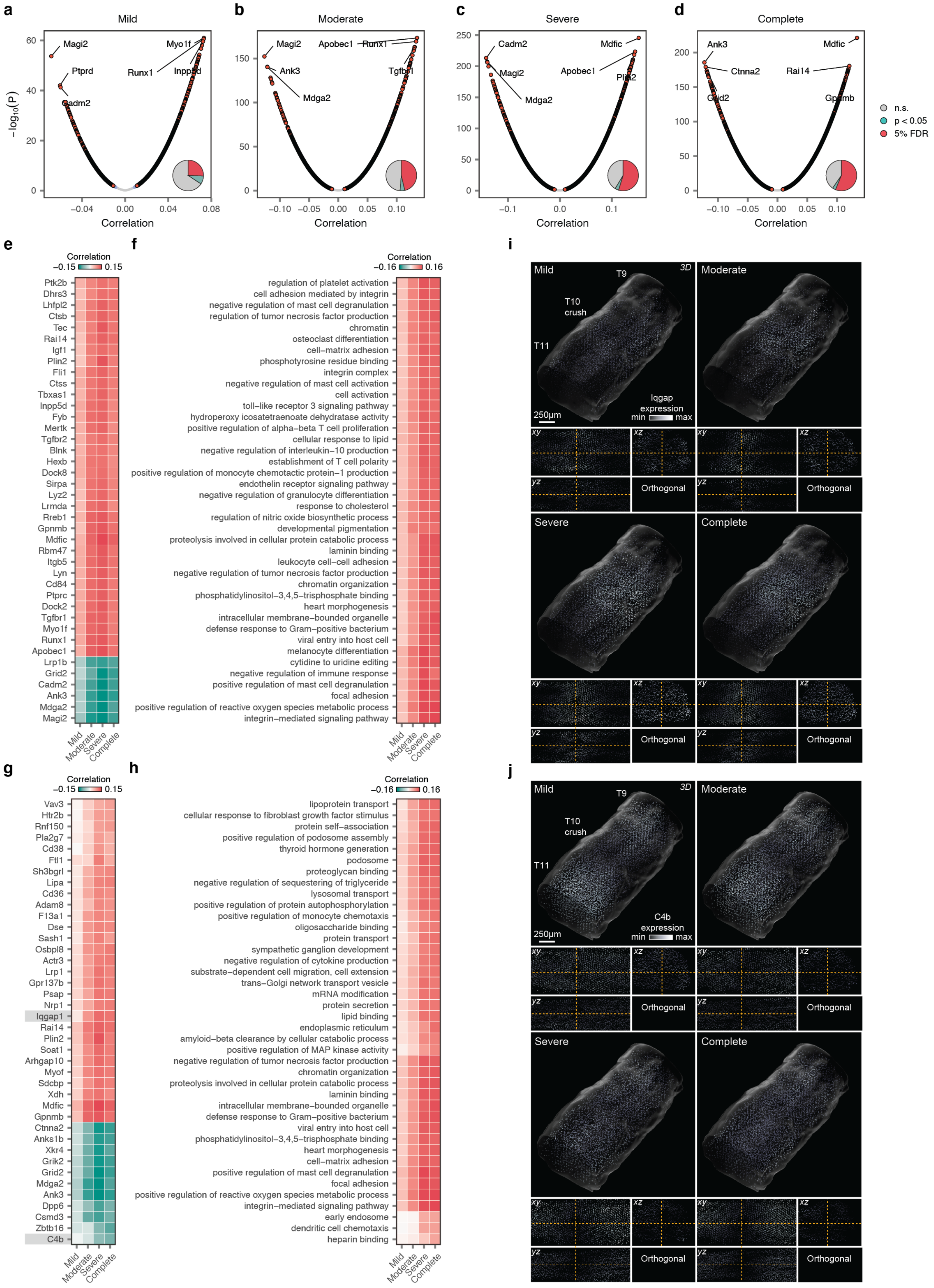
a
Mild
Magi2
Myo1f
Ptprd
Runx1
Inpp5d
Cadm2
−log10(P)
Correlation
b
Moderate
Magi2
Apobec1
Runx1
Ank3
Tgfbr1
Mdga2
c
Severe
Cadm2
Mdfic
Magi2
Apobec1
Plin2
Mdga2
d
Complete
Ank3
Mdfic
Ctnna2
Rai14
Grid2
Gpnmb
n.s.
p < 0.05
5% FDR
e
Correlation
Ptk2b
Dhrs3
Lhfpl2
Ctsb
Tec
Rai14
Igf1
Plin2
Fli1
Ctss
Tbxas1
Inpp5d
Fyb
Mertk
Tgfbr2
Blnk
Hexb
Dock8
Sirpa
Lyz2
Lrmda
Rreb1
Gpnmb
Mdfic
Rbm47
Itgb5
Lyn
Cd84
Ptprc
Dock2
Tgfbr1
Myo1f
Runx1
Apobec1
Lrp1b
Grid2
Cadm2
Ank3
Mdga2
Magi2
Mild
Moderate
Severe
Complete
f
regulation of platelet activation
cell adhesion mediated by integrin
negative regulation of mast cell degranulation
regulation of tumor necrosis factor production
chromatin
osteoclast differentiation
cell−matrix adhesion
phosphotyrosine residue binding
integrin complex
negative regulation of mast cell activation
cell activation
toll−like receptor 3 signaling pathway
hydroperoxy icosatetraenoate dehydratase activity
positive regulation of alpha−beta T cell proliferation
cellular response to lipid
negative regulation of interleukin−10 production
establishment of T cell polarity
positive regulation of monocyte chemotactic protein−1 production
endothelin receptor signaling pathway
negative regulation of granulocyte differentiation
response to cholesterol
regulation of nitric oxide biosynthetic process
developmental pigmentation
proteolysis involved in cellular protein catabolic process
laminin binding
leukocyte cell−cell adhesion
negative regulation of tumor necrosis factor production
chromatin organization
phosphatidylinositol−3,4,5−trisphosphate binding
heart morphogenesis
intracellular membrane−bounded organelle
defense response to Gram−positive bacterium
viral entry into host cell
melanocyte differentiation
cytidine to uridine editing
negative regulation of immune response
positive regulation of mast cell degranulation
focal adhesion
positive regulation of reactive oxygen species metabolic process
integrin−mediated signaling pathway
i
Mild
T9
3D
T10
crush
T11
Iqgap
expression
min max
250µm
Moderate
Severe
Complete
xy
xz
yz
Orthogonal
g
Vav3
Htr2b
Rnf150
Pla2g7
Cd38
Ftl1
Sh3bgrl
Lipa
Cd36
Adam8
F13a1
Dse
Sash1
Osbpl8
Actr3
Lrp1
Gpr137b
Psap
Nrp1
Iqgap1
Rai14
Plin2
Soat1
Arhgap10
Myof
Sdcbp
Xdh
Mdfic
Gpnmb
Ctnna2
Anks1b
Xkr4
Grik2
Grid2
Mdga2
Ank3
Dpp6
Csmd3
Zbtb16
C4b
h
lipoprotein transport
cellular response to fibroblast growth factor stimulus
protein self−association
positive regulation of podosome assembly
thyroid hormone generation
podosome
proteoglycan binding
negative regulation of sequestering of triglyceride
lysosomal transport
positive regulation of protein autophosphorylation
positive regulation of monocyte chemotaxis
oligosaccharide binding
protein transport
sympathetic ganglion development
negative regulation of cytokine production
substrate−dependent cell migration, cell extension
trans−Golgi network transport vesicle
mRNA modification
protein secretion
lipid binding
endoplasmic reticulum
amyloid−beta clearance by cellular catabolic process
positive regulation of MAP kinase activity
negative regulation of tumor necrosis factor production
chromatin organization
proteolysis involved in cellular protein catabolic process
laminin binding
intracellular membrane−bounded organelle
defense response to Gram−positive bacterium
viral entry into host cell
phosphatidylinositol−3,4,5−trisphosphate binding
heart morphogenesis
cell−matrix adhesion
positive regulation of mast cell degranulation
focal adhesion
positive regulation of reactive oxygen species metabolic process
integrin−mediated signaling pathway
early endosome
dendritic cell chemotaxis
heparin binding
j
C4b
expression

**Supplementary Fig. 43 | Molecular basis of three-dimensional spatial prioritization at the gene and gene module levels.**
**a-d,** Volcano plots showing correlations between the AUCs assigned by Magellan to each spatialized barcode from the snRNA-seq and transcriptome-wide gene expression across the same snRNA-seq barcodes (**a,** mild; **b,** moderate; **c,** severe; **d,** complete). Inset pie charts show the proportions of all tested genes that are significantly correlated with the spatial prioritizations.
**e,** Heatmap showing Pearson correlations between spatial prioritizations and gene expression for the top 40 most recurrently correlated genes across injury severities.
**f,** Heatmap showing Pearson correlations between spatial prioritizations and gene expression for the top 40 most variably correlated genes across injury severities.
**g,** As in **e,** but for gene modules.
**h,** As in **f,** but for gene modules.
**i,** Expression of selected genes from **f** across injury severities, visualized on the 3D spatial transcriptomic atlas (**i,** *Iqgap*; **j,** *C4b*).

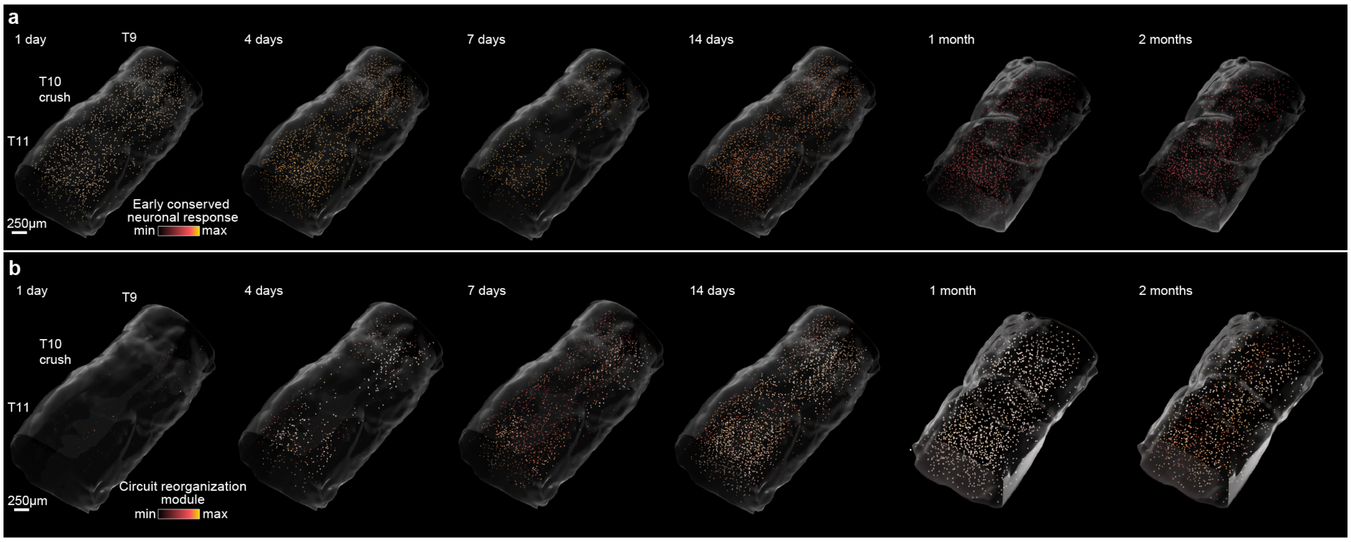


**Supplementary Fig. 44 | Early-conserved and late-diverging neuronal responses to SCI in the 3D spatial atlas.**
**a,** Expression of the early conserved neuronal response module in spatialized neurons from the snRNA-seq atlas, visualized on the coordinate system of the 3D spatial transcriptomic atlas.
**b,** Expression of the circuit reorganization module in spatialized neurons from the snRNA-seq atlas, visualized on the coordinate system of the 3D spatial transcriptomic atlas.

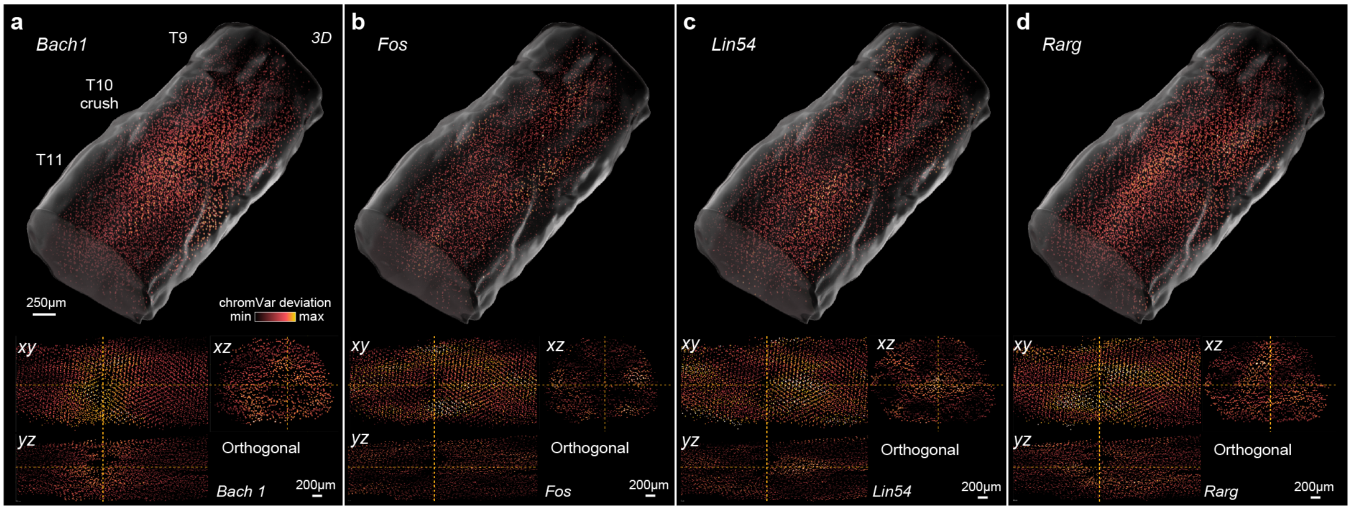


**Supplementary Fig. 45 | Three-dimensional mapping of chromatin accessibility.**
Accessibility of motifs associated with tripartite barrier formation in spatialized vascular cells from the multiome atlas, visualized on the coordinate system of the 3D spatial transcriptomic atlas. **a,** *Bach1*; **b,** *Fos*; **c,** *Lin54*; **d,** *Rarg*.

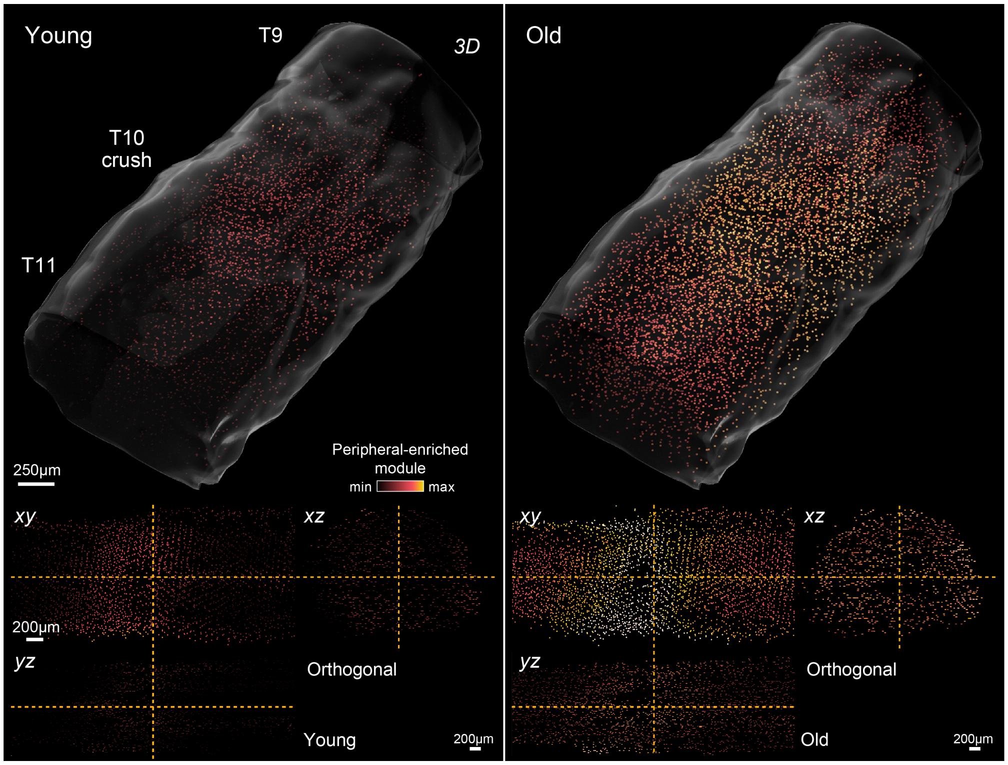


**Supplementary Fig. 46 | Loss of blood-brain barrier transcriptional identity in old mice.**
Expression of genes that distinguish peripheral endothelial cells from cells of the blood-brain barrier in spatialized vascular cells from young versus old mice, visualized on the coordinate system of the 3D spatial transcriptomic atlas.

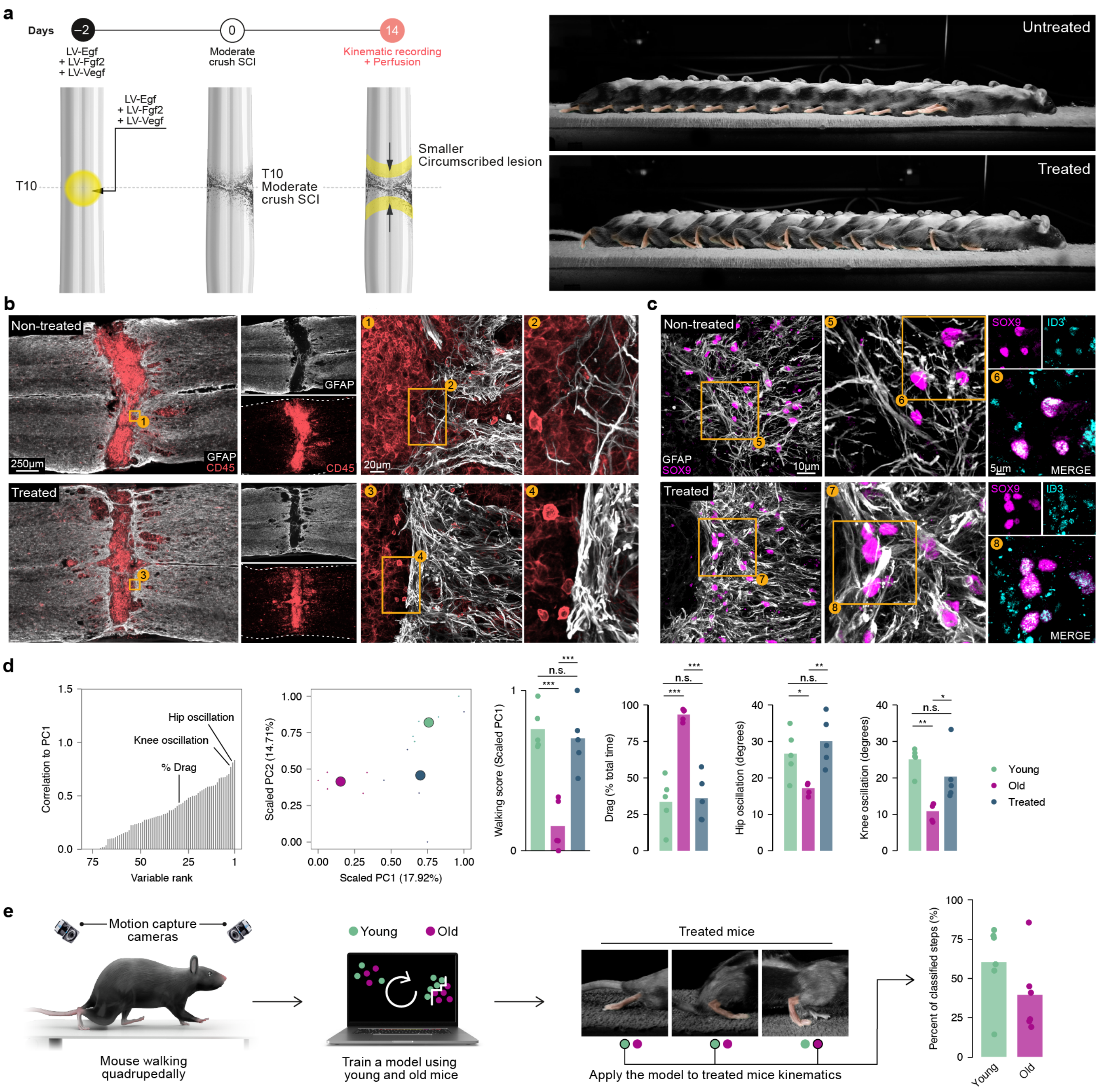


**Supplementary Fig. 47 | A rejuvenative gene therapy reestablishes the tripartite barrier to restore walking.**
**a,** Left, experimental design of a gene therapy intervention to promote the formation of the tripartite barrier, reproduced from **Fig. 8a**. Right, a second chronophotography series showing walking in old mice without (top) and with (bottom) a gene therapy intervention to promote the formation of the tripartite barrier.
**b,** Composite tiled scans of GFAP and CD45 in horizontal sections from representative old and treated mice.
**c,** Horizontal sections from representative old and treated mice identifying a restoration of Sox9$^{ON}$Id3$^{ON}$ cells in the astrocyte border region in treated mice.
**d,** Locomotor performance in the gene therapy experiment, as quantified in **Supplementary Fig. 2** (*n* > 10 gait cycles per mouse, (*n* = 5 mice per group; statistics indicate Tukey HSD tests following one-way ANOVA). *, p < 0.05; **, p < 0.01; ***, p < 0.001.
**e,** Left, schematic overview of the classification pipeline using high-resolution kinematics data from young and old mice. Right, experimental conditions assigned to individual steps in old mice that received gene therapy by a machine-learning model trained on kinematics data from untreated animals.
